# Revealing the Auxetic Behavior of Biomimetic Multi-material and Region-specific Nanofibrous Scaffolds via Synchrotron Multiscale Digital Volume Correlation: Innovative Building Blocks for the Enthesis Regeneration

**DOI:** 10.1101/2024.08.12.607645

**Authors:** Alberto Sensini, Francesca Giacomini, Olga Stamati, Bratislav Lukic, Julie Villanova, Henry Proudhon, Maryse Gille, Zeinab Tahmasebi Birgani, Roman Truckenmüller, Gianluca Tozzi, Martijn van Griensven, Lorenzo Moroni

**Affiliations:** MERLN Institute for Technology-Inspired Regenerative Medicine Complex Tissue Regeneration Department Maastricht University P.O. Box 616, 6200 MD, Maastricht, The Netherlands; MERLN Institute for Technology-Inspired Regenerative Medicine Cell Biology-Inspired Tissue Engineering Department Maastricht University P.O. Box 616, 6200 MD, Maastricht, The Netherlands; The European Synchrotron Radiation Facility CS 40220, 38043 Grenoble Cedex 9, Grenoble, France; University Grenoble Alpes, CNRS Grenoble INP, 3SR, F-38000 Grenoble, France; Henry Royce Institute, Department of Materials, The University of Manchester, UK Manchester, M13 9PL, United Kingdom; MINES Paris, PSL University, MAT – Centre des matériaux, CNRS UMR 7633, BP 87, F-91003 Evry, France; School of Engineering, Faculty of Engineering & Science University of Greenwich, ME4 4TB, Chatham Maritime, United Kingdom

**Keywords:** electrospinning, nanofibers, enthesis, digital volume correlation, synchrotron nanoCT, mesenchymal stromal cells spheroids, mineralization

## Abstract

Enthesis lesions are one of the prevalent causes of injuries in the tendon tissue. The gradient of mineralization, extracellular matrix organization and auxetic mechanical properties, make enthesis regeneration challenging. Innovative electrospun fascicle-inspired nanofibrous poly(L-lactic)acid/collagen type I blend scaffolds were developed. Specifically, a mineralized fibrocartilage-inspired region (with/without nano-mineralization with hydroxyapatite), where random and aligned nanofibers coexist, is connected to a tendon-like region made of aligned nanofibers, through a conical non-mineralized fibrocartilage-inspired junction. Scanning electron microscopy and synchrotron nano-tomography show the morphological biomimicry of scaffolds with the natural tendon fascicles. Human mesenchymal stromal cells spheroids cultures confirm a balanced expression of both tendon, cartilage and bone markers on the non-mineralized scaffolds compared with the mineralized ones. Mechanical tests, at different physiological strain-rates, reveal a biomimetic mechanical behavior of scaffolds and the ability of junctions to tune the mechanics of their surrounding sites. Multiscale synchrotron *in situ* tensile tests, coupled with Digital Volume Correlation, elucidate the full-field strain distribution of scaffolds from the structural down to the nanofiber level, highlighting the auxetic mechanical behavior of junctions typical of the natural enthesis. The findings and cutting-edge investigations of our study suggest the suitability of these enthesis-inspired fascicles as innovative scaffolds for enhanced enthesis regeneration.

## 1. Introduction

The regeneration of injured tendons and ligaments (T/L) is a crucial societal need worldwide accounting for approximately 30 million cases annually.[1–3] Among these, a relevant role is conferred to the enthesis (i.e. tendon/ligament-to-bone insertion). The most common enthesis sites of injury include the rotator cuff, the anterior cruciate ligament, the flexor, the patellar and the Achilles tendon.[4] Moreover, the failure rates after surgery for enthesis repair are extremely high (e.g. rotator cuff: 20-94%, anterior cruciate ligament 10-25%).[5,6] The reason behind these criticalities in healing the enthesis lies in its complex structure and biomechanical properties.[7] In fact, the main biological function of the enthesis is to guarantee a progressive attachment between the T/L tissue and the trabecular bone, reducing the risk of undesired interfacial stress concentrations.[8] The stress deconcentrating nature of the enthesis is also confirmed by the typical interdigitated nature of its attachment with the bone tissue.[8–10] For these reasons, the enthesis morphology is based on a structural gradient of fibrillar collagen organization and mineralization of the extracellular matrix (ECM).[7] Typically, four different, but structurally continuous, regions can be distinguished within the fibrocartilaginous enthesis **(Figure 1A, 1B)**: the T/L tissue, the non-mineralized fibrocartilage, the mineralized fibrocartilage and the bone tissue.[11] The T/L zone of the enthesis consists of axially aligned fibrils with elongated tenocytes/fibroblasts in between.[12] Here, the ECM is mostly composed of collagen type I and a small amount of proteoglycans.[7] The following region is the non-mineralized fibrocartilage, rich of fibrochondrocytes, mostly collagen type II, III (less percentages of collagen types I, X) and proteoglycans.[8,13] Then, the mineralized fibrocartilage which is populated by hypertrophic fibrochondrocytes and rich of collagen type II, X, aggrecan and small amounts of hydroxyapatite. Finally, the mineralized fibrocartilage connects the bone tissue with osteoblasts, osteocytes, osteoclasts and an ECM rich in collagen type I and hydroxyapatite.[8,13] Transitioning from the fibrocartilage to the bone tissue, the ECM matrix progressively loses its anisotropy becoming more disorganized.[7] The morphological changes in the ECM are followed by a progressive mineralization from the enthesis up to the bone tissue, causing a consequent gradient of mechanical properties along the enthesis tissue.[11] To verify these strain gradients, full-field 2D and 3D imaging techniques coupled with mechanical tests, such as Digital Image Correlation (DIC),[14–16] Digital Volume Correlation (DVC)[17,18] and confocal microscopy[10] were adopted. Interestingly, DIC and DVC were also able to detect the auxetic behavior of tendon tissue at the enthesis side both in human and animal tissues.[17,19]

**Figure 1.**
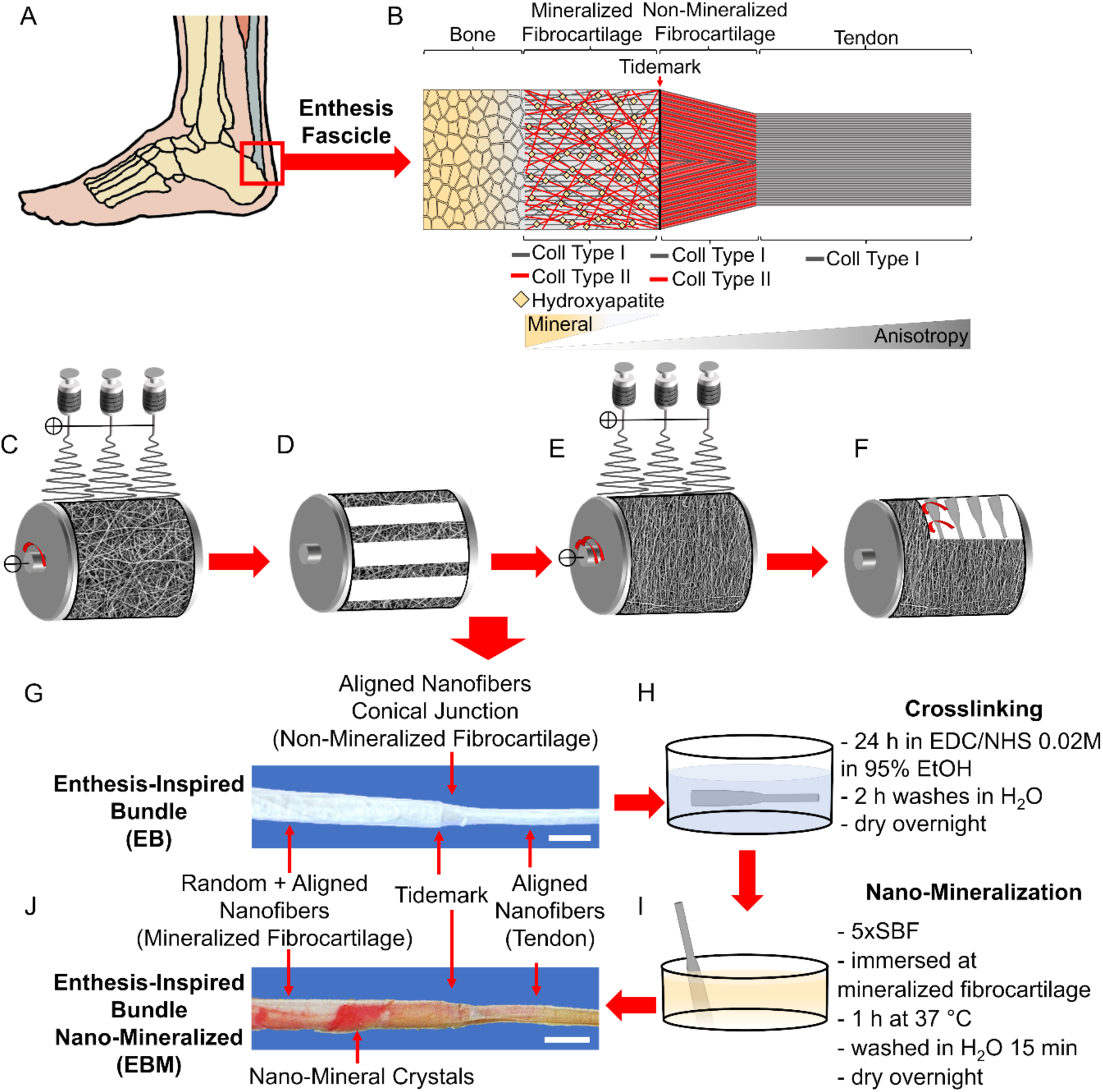
Structure of natural enthesis fascicles and enthesis-inspired electrospun bundles biofabrication. A) Example of anatomical site of natural enthesis; B) schematic of structure and composition of a natural enthesis fascicle. Fabrication of enthesis fascicle-inspired bundles: C) electrospinning of a mat of random nanofibers on a low-speed rotating drum collector; D) removal of axial stripes of mat; E) electrospinning of a mat of aligned nanofibers on a high-speed rotating drum collector; F) Axial and circumferential cuts of the mat on the drum and rolling up of rectangular stripes to obtain bundles; G) enthesis fascicle-inspired bundle (sale bar = 1 mm); H) crosslinking of collagen I done with a solution of N-(3-Dimethylaminopropyl)-N’-ethylcarbodiimide hydrochloride (EDC), N-Hydroxysuccinimide (NHS) and ethanol (EtOH); I) production of nano-mineralization on nanofibers by immersing bundles inside simulated body fluid (SBF) at the mineralized fibrocartilage region; J) enthesis fascicle-inspired bundle nano-mineralized (sale bar = 1 mm).

Aiming to regenerate the enthesis tissue with biomimetic devices, several synthetic scaffolds were developed using a panel of different biofabrication techniques.[7,20–24] Some studies also focused on the development of auxetic mechanical metamaterials,[25–27] by using additive manufacturing techniques, also in the attempt to mimic the auxetic behavior of the T/L tissue.[28] Among the different biofabrication methods, electrospinning has demonstrated to be suitable for this scope.[29] Researchers have proposed several electrospun scaffolds to mimic the complex environment of the tendon/ligament-to-bone interface by using mineralization gradients (random or aligned mats of nanofibers),[30–35] region specific nanofiber orientations[36–38] or a mix of them.[39,40] DIC investigations during tensile tests on electrospun mats, with mineral gradients, have preliminarily demonstrated the presence of strain gradients on the surface of scaffolds mimicking the natural behavior of the enthesis tissue.[30,33] Moreover, stem cells have been stimulated to produce enthesis specific markers when cultured on these morphological and mineral patterns.[34,35,37,38,40–43] Despite the promising results obtained so far, several open questions persist. Even if some preliminary electrospun partially mineralized yarns were proposed,[44,45] the design of scaffolds able to faithfully mimic enthesis fascicles, reproducing their typical stress-concentration reducer auxetic conical shape, as well as their morphological and mechanical gradients is still an unsolved open challenge. In fact, even though some auxetic behaviors of electrospun random mats were recently reported,[46,47] the development of mechanically relevant electrospun auxetic scaffolds remains undocumented so far. Although some works verified a superficial strain gradient in their electrospun mats, a full-field X-ray 3D imaging mechanical investigation coupled with DVC inside the volume of enthesis-inspired electrospun scaffolds is totally unexplored so far. This is mainly due to challenges in obtaining an adequate nanometric voxel size, mandatory to properly visualize the electrospun nanofibers, while performing fast tomography, to prevent relaxation of scaffolds during the stepwise tests. In fact, after years of unsuccessful attempts, only recently a group of researchers managed to design a laboratory micro-computed tomography (microCT) stepwise tensile test and DVC to investigate the mechanics of electrospun materials.[48] Despite the promising preliminarily results, the micrometric voxel size adopted was not sufficient to visualize the evolution of the nanofibers inside the scaffolds during the tests.

With this in mind, we developed an innovative electrospun enthesis-inspired conical fascicle, made of a blend of high molecular weight poly(L-lactic) acid (PLLA) and collagen type I (COL-I), with region-specific nanofibers orientations. Moreover, to study the benefits of mineralization, a group of scaffolds was nano-mineralized with hydroxyapatite nanocrystals. Their morphology was first investigated via scanning electron microscopy (SEM) and the macroscopic mechanical behavior was then assessed via mechanical tensile tests at three different physiological strain rates. Spheroids of human mesenchymal stromal cells (hMSCs) were cultured on the enthesis-inspired bundles up to 28 days revealing that the best candidates for a contemporary tendon, cartilage and bone markers expression over time were the non-mineralized scaffolds. These were then used to produce a comprehensive synchrotron multiscale micro-(SµCT) to nano-(SnCT) X-ray computed tomography investigation coupled with *in situ* tensile testing and DVC revealing, for the first time, the full-field strain distribution at the nanofiber level and a surprising auxetic enthesis-inspired mechanical behavior at the conical junction.

## 2. Results and Discussion

### 2.1 Production and Morphology of Enthesis-Inspired Scaffolds

To faithfully reproduce electrospun scaffolds with region-specific nanofibers orientations, mimicking the morphology of the enthesis (Figure 1A, 1B) (i.e. mineralized fibrocartilage, non-mineralized fibrocartilage and tendon tissue)[49], innovative electrospun enthesis fascicle inspired bundles (EB) made of a blend of high-molecular weight medical grade PLLA and Coll in a percentage of 75/25 (w/w) were produced. In brief, the biofabrication method consists in two consecutive electrospinning steps involving a drum collector rotating at different speeds (see methods section). In this way, a first mat of randomly arranged nanofibers was produced (Figure 1C). Subsequently the mat was cut in axial stripes, which were alternatively removed obtaining empty bands between two random nanofibers stripes (Figure 1D). A subsequent mat of axially aligned nanofibers was electrospun on top of the random stripes (Figure 1E). Further axial cuts were produced obtaining stripes with the contemporary presence of sections of random + aligned nanofibers and fully aligned nanofibers along the drum circumference. Finally, by producing additional circumferential cuts and rolling up the obtained stripes, it was possible to fabricate scaffolds with region-specific diameters and nanofibers’ orientations: a mineralized fibrocartilage-inspired region made of random + aligned nanofibers (FB) (nanofibers’ diameter random (d_R_) and aligned (d_A_): d_R_ = 0.39 ± 0.19 µm, d_A_ = 0.30 ± 0.16 µm; FB diameter: D_FB_ = 733 ± 20 µm) and a tendon-inspired region consisting of axially aligned nanofibers (TB) (same d_A_ reported above; TB diameter: D_TB_ = 450±30 µm). Moreover, by connecting them, a biomimetic non-mineralized fibrocartilage-inspired conical junction (J) of aligned nanofibers was generated (EB diameter: D_EB_ = 635 ± 10 µm) (Figure 1F, 1G and **Figure 2**). To closely mimic the natural mineralized fibrocartilage-bone tissue side, the FB region of a group of bundles was mineralized with amorphous nano-hydroxyapatite crystals (crystals diameter = 0.29±0.28 µm) producing a nano-mineralized fibrocartilage region (FBM) (mineralized nanofibers’ diameter (d_M_): d_M_ = 0.41 ± 0.16 µm; FBM diameter: D_FBM_ = 733 ± 30 µm) (Figure 1J and Figure 2AIV, 2BI, 2BIII, 2BIV). This allowed obtaining enthesis-inspired mineralized bundles (EBM) (d_M_ for the FBM side and d_A_ for the TB and J side; EBM diameter: D_EBM_ = 612±10 µm) (**Table S1**).

**Figure 2.**
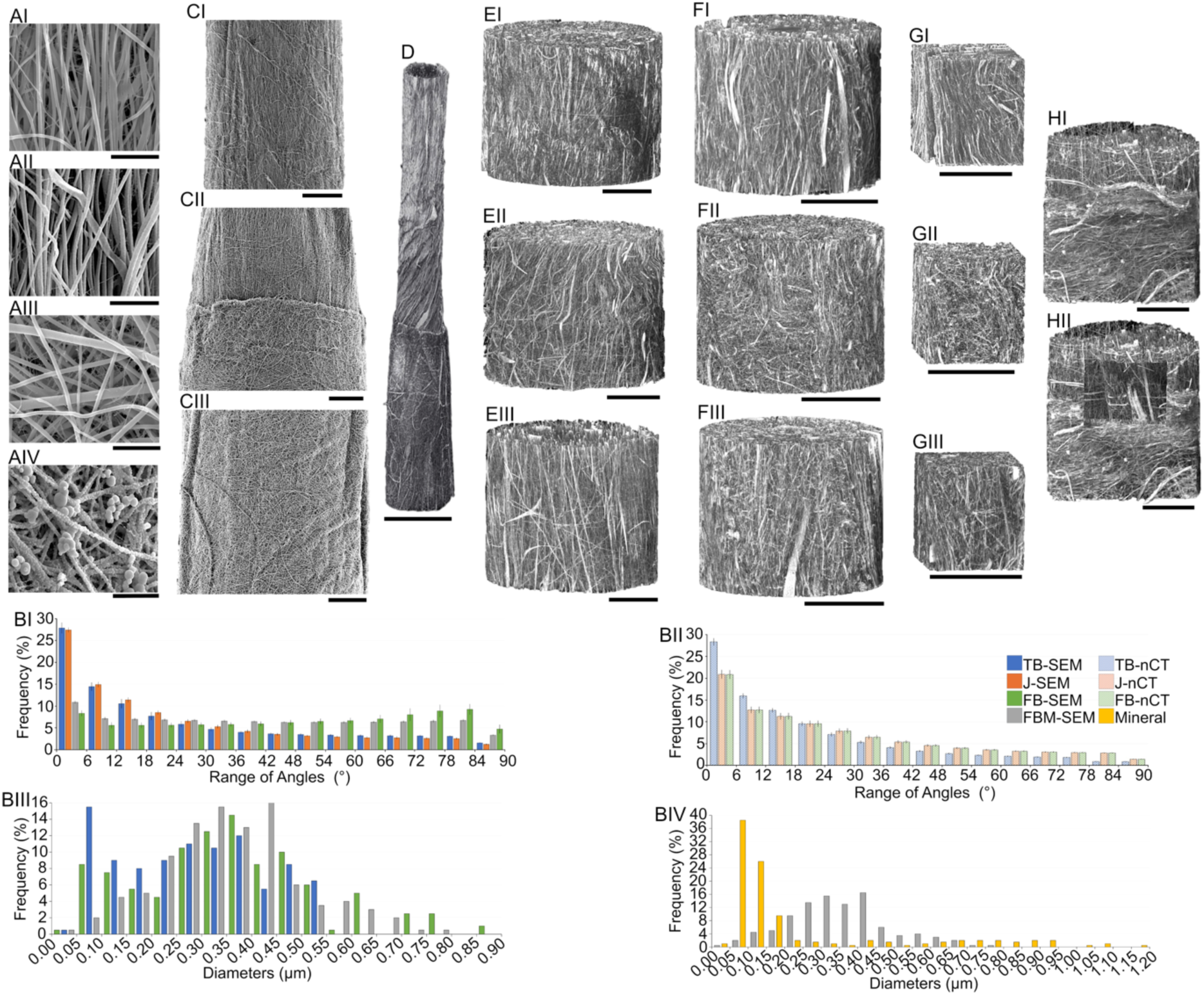
SEM and SnCT morphological investigation of enthesis-inspired bundles. A) SEM of PLLA/Coll nanofibers (scale bar = 5 µm, magnification = x5000): AI) tendon side; AII) non mineralized fibrocartilage side (junction); AIII) FB side without nano-mineralization; AIV) FBM side with nano-mineralization. B) Nanofibers and nano-mineral orientation and diameter distribution: BI) nanofibers orientation, via SEM images, on the surface of the scaffolds (0° = axial orientation; 90° = transversal orientation; images at x5000); BII) nanofibers orientation inside the scaffolds, from SnCT reconstructed sub-volumes (voxel size = 100 nm); BIII) diameter distribution of the nanofibers in the different regions of the scaffolds; BIV) diameter distribution of the nano-mineral crystals compared with mineralized nanofibers diameters. C) SEM images of the different regions of an enthesis-inspired bundle (scale bar = 100 µm): CI) TB side (magnification = x200); CII) J side (magnification = x190); CIII) and FB side (magnification = x190). Synchrotron X-ray 3D imaging multiscale investigation of D) the whole EB scaffold (SµCT, scale bar = 600 µm; voxel size = 650 nm) and its different regions (i.e. I = TB side; II = J side; III = FB side) at different voxel sizes (SnCT): E) voxel size = 300 nm (scale bar = 100 µm); F) voxel size = 100 nm (scale bar = 50 µm); G) internal crops of the different regions at voxel size = 100 nm (scale bar = 50 µm). H) SnCT images of the junction (voxel size = 100 nm; scale bar = 50 µm): HI) external overview; HII) central crop showing an internal layer of aligned nanofibers under the external random layer of the FB region.

The biomimetic diameters and orientations of nanofibers led the scaffolds to achieve region-specific porosity (P) that was for the TB side of P_TB_ = 51 ± 3 %, for the FB of P_FB_ = 56 ± 6 % and for FBM P_FBM_ = 55 ± 5 %. These values conferred a global porosity at EB of P_EB_ = 61 ± 4 % that slightly decreased for EBM resulting in P_EBM_ = 55 ± 5 % (Table S1).

Concerning the morphology of scaffolds and nanofibers (Figure 2A and 2C), the SEM images highlighted the smoothness of nanofibers and the overall biomimicry of the different regions of EB and EBM, compared to the natural counterparts.[10,11,50–52] Moreover, the length of conical junction (∼500 µm) and the net interface between the non-mineralized and mineralized fibrocartilage inspired regions (i.e. tidemark) (Figure 2CII) were consistent with the length of the natural enthesis reported in literature.[10]

The nanofibers’ orientations, calculated from the SEM images on the surface of crosslinked scaffolds, closely mimic the progressive loss of anisotropy, from the tendon to the bone tissue, described in literature.[7] Specifically, the strong axial alignment, with a small transversal dispersion, of nanofibers at TB (range: 0° - 18° = 53 ± 3 %; 78° - 90° = 4.7 ± 0.3 %) and at J (range: 0° - 18° = 54 ± 2 %; 78° - 90° = 3.8 ± 0.4 %) regions, were in contrast with the randomic orientation at FB (range: 0° - 18° = 20 ± 2 %; 78° - 90° = 14 ± 2 %) and FBM (range: 0° - 18° = 25 ± 1 %; 78° - 90° = 10.1 ± 0.4 %) with an isotropic distribution in the different ranges of angles (Figure 2BI). This organization offered an immediate biomimetic region-specific pattern useful to guide hMSC in the production enthesis-specific markers and in their differentiation.[53] Furthermore, the crosslinking process caused a heterogeneous shrinkage (sk) of scaffolds in their different regions (i.e. s_TB_ = 14 ± 1 %; s_FB_ = 16 ± 1 %; s_FBM_ = 16 ± 2 %) resulting in a total sk of EB and EBM of s_EB_ = 19 ± 1 % and s_EBM_ = 19 ± 3 %. The higher level of sk in FB and FBM regions was attributed to the contemporary presence of overlapped layers of random and aligned nanofibers. In fact, even if only random nanofibers were visible by SEM in the most external layer (Figure 2AIII, 2AIV), the internal layers of aligned nanofibers were clearly visible from the later SnCT and SµCT multiscale investigation on EB (Figure 2D-2H). This analysis allowed also to confirm the internal presence of aligned layers of nanofibers in the FB region (Figure 2BII). We were able to confirm the full-field maintenance of the strong axial alignment of the TB nanofibers of EB (range: 0° - 18° = 57 ± 2 %; 78° - 90° = 1.8 ± 0.1 %) and an internal partial alignment of the FB region of EB (range: 0° - 18° = 45 ± 2 %; 78° - 90° = 4.3 ± 0.1 %) with similar values as the J region (range: 0° - 18° = 45 ± 2 %; 78° - 90° = 4.3 ± 2 %). This effect, for the contemporary presence at the FB region of random and aligned nanofibers, contributed to generate a pre-strain of the FB/FBM region, driven by the internal axially aligned nanofibers, which caused a progressive alignment of hMSCs also in this part of the scaffold (see **Figure 4** and **Figure 5**). 3D videos of the SnCTs and SµCT reconstructions can be found in **Video S1-S8**.

### 2.2 Mechanical Properties of Enthesis-Inspired Scaffolds

To assess the mechanical performances of EB and EBM, both the whole scaffolds and their single parts (i.e. TB, FB and FBM) were tested via a tensile test at different physiological strain rates of natural tendon tissue: a quasi-static task (0.4% s^-1^), a fast walk or slow run (10 % s^-1^), and a worst case scenario of failure (100% s^-1^)[54–56] (**Figure 3**, **Table S3-S8**). The mechanical tests on scaffolds revealed a nonlinear ductile mechanical behavior with large deformations (Figure 3A-C). All bundles showed an initial toe-region, up to around 1 - 2% of strain, typical of the natural tissue counterpart (Figure 3AI-CI).[52] The strain-rate dependency of bundles contributed to progressively increase their mechanical properties reaching values that fell into the range of the tendon fascicles reported in literature (Figure 3, Table S3-S8) and in line with the natural viscoelastic nature of tendon fascicles.[57] It is worth mentioning that, despite the presence of the nano-mineralization which contributes to increase the strength and stiffness of FBM and EBM, all bundles preserved their ductile region reaching similar ranges of values of yield/failure strain (Figure 3F-3G). In fact, no significant differences in terms of yield and failure strain were detected between the different categories of scaffolds, confirming the maintenance of their ductility both in presence of mineralization and at different strain rates. This is a desired characteristic of scaffolds and orthopedic devices since it prevents them from an unexpected failure in case of overload. Moreover, EMB and FBM maintained their ductility, even in presence of the nano-mineralization, while slightly increasing their mechanical properties (i.e. F_Y_, ο_Y_, K and E), making them easy to be handle also in case of surgical scenarios. Structurally speaking, aiming to simulate the stiffness and morphology of the mineralized fibrocartilage, the simultaneous presence of layers of random and aligned nanofibers at the FB and FBM, contributed to increase the amount of resisting material at these regions/scaffolds. In addition, the adopted biofabrication strategy allowed to immediately expose hMSCs to a surface of random nanofibers suitable to enhance their osteogenic/chondrogenic differentiation.[58,59] In terms of mechanical performances, FB and FBM sides showed the highest values, in a strain-rate dependent fashion of yield/failure force and stiffness compared to the other sample categories, also in a statistically significant manner (Figure 3D, 3E and EH, Table S5). The stiffness and force developed by FB/FBM further corroborate the potential of hMSCs to express osteogenic and chondrogenic markers.[60]

**Figure 3.**
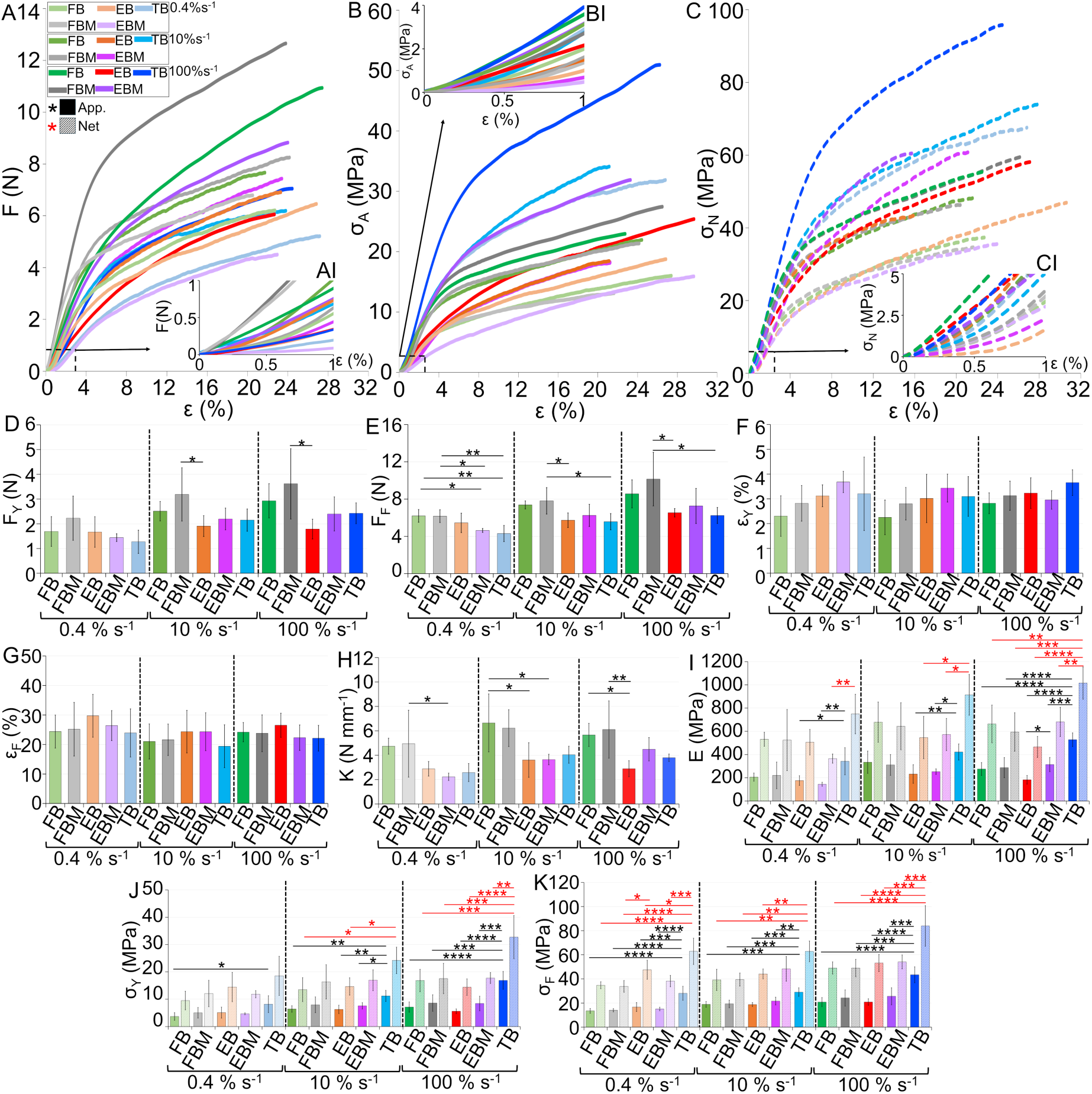
Mechanical characterization of bundles at different strain-rates (0.4, 10 and 100 % s^-1^; apparent properties = solid lines/bars, net properties = dashed lines/bars): A) typical force-strain, B) apparent stress-strain and C) net stress-strain curves (AI, BI, CI zoom in at the toe region); D) yield force (F_Y_); E) failure force (F_F_); F) yield strain (ε_Y_); G) failure strain (ε_F_); H) stiffness (K); I) elastic modulus (E); J) yield stress (ο−_Y_); K) failure stress (ο−_F_). The statistically significant differences between the different bundles categories, at different strain-rates, were assessed with a one-way ANOVA followed by a Tukey post hoc (ns p>0.05, *p≤0.05, **p≤0.01, ***p≤0.001, ****p≤0.0001; apparent properties = black asterisks; net properties = red asterisks).

In terms of stress and elastic modulus, FB/FBM showed reduced values of apparent properties due to their slightly higher cross-section and the presence of layers of random nanofibers compared to the TB ones (Figure 3I-3K, Table S3). Notably, the TB apparent yield/failure stress and elastic modulus showed the same values as human Achilles tendon fascicles reported in literature, being also statistically significantly higher compared to the other scaffolds categories/sides.[57] These properties were found suitable to promote tendon-specific ECM and tenogenic markers.[60]

Concerning EB and EBM, both scaffolds showed intermediate mechanics with respect to FB, FBM and TB. The conical non-mineralized fibrocartilage-inspired junction, placed in between their gauge length and their FB (FBM for EBM) and TB sides, demonstrated to modulate the stiffness of FB/FBM and the high elastic modulus of the TB side as the natural enthesis does (Figure 3H-3I, Table S3). This effect could be attributed to the high shrinkage and waviness of the nanofibers in correspondence with the junction area (Figure 2CII, EII, FII). In fact, the slightly higher strains (and displacements) required to realign the nanofibers in correspondence to the junction (Figure 3F-3G, Table S3). In general, the contribution of the nano-mineralization was evident at higher strain-rates where EBM resulted stiffer and with higher moduli compared to EB (Figure 3H-3I). Both EB and EBM were able to well mimic the natural function of the enthesis, to act as stress concentration reducers with respect to their FB/FBM and TB sides, while maintaining mechanical properties fully in line with the natural counterpart.[57,61] Furthermore, the net mechanical properties of all scaffolds (i.e. computed considering their volume fraction, so only the real resisting material) were around 2-3 times higher compared to the apparent ones, confirming the high mechanical competence of all EB, EBM and all their sides (FB, FBM and TB). This makes them highly mechanically biomimetic with the natural tissue (Figure 3 and Table S3 and Table S4).[62,63] These aspects allowed both EB, EBM and their components to store significant amounts of work before yield and failure (Table S3, S5 and S7). It is worth mentioning that no EB and EBM broke at their junctions during all tensile tests, confirming the mechanical stability of this design.

To further confirm even more the mechanical biomimicry of the gradient of mechanical properties expressed by EB, EBM and their different components, all scaffolds showed the presence of inflection points (both apparent and net stress and strains) fully in line with the ones of the natural tendon tissue reported in literature (Figure S4, S6, S8).[64] This point detects the initial transition between the strain stiffening and the strain softening of the linear region of the stress-strain curve, detecting the first ruptures/yields of collagen fibrils and nanofibrous materials[65]. These values are pivotal to trigger tendon cells in the regeneration of natural enthesis and tendon tissue.[66]

### 2.3. In Vitro hMSCs Proliferation, Differentiation and ECM Production

To investigate the capability of EB and EBM to self-support hMSCs proliferation, differentiation and production of enthesis-specific ECM, hMSCs spheroids were cultured on scaffolds up to 28 days in basic culture media. Spheroids were chosen since they provide a three-dimensional (3D) environment that promotes natural cell-cell interactions, similar to those found in vivo. This can enhance cellular functions, such as increased viability and secretion of extracellular matrix components and mimic physiological conditions more accurately than two-dimensional (2D) cultures.[67,68]

#### 2.3.1. Alignment of actin fibers and nuclei shape of cultured hMSCs and formed ECM

hMSCs cultured on both EB and EMB scaffolds, showed region-specific morphologies when proliferated on their different regions (Figure 4). After 7 days of culture, SEM investigation revealed that hMSCs were able to attach and colonize both types of scaffolds (Figure 4A-4F I, II). The hMSCs attached along the TB and J regions (Figure 4AI, 4AII, 4BI, 4BII, 4DI, 4DII, 4EI, 4EII), showed a prevalent axial orientation with respect to the bundle’s axis starting from day 7, following the predominantly aligned direction of nanofibers (Figure S3 and Table S9). Conversely, enlarged and spread morphologies of hMSCs were observed at the FB and FBM regions (Figure 4CI, 4CII, 4FI, 4FII), in accordance with the random orientation of nanofibers on the surface of scaffolds in those regions. At 14 and 28 days, along the length of EB and EBM, nanofibers became progressively covered by an increasing number of axially aligned cells and ECM that, at 28 days, reached a quite uniform layer along the length of both EB and EBM (Figure 4A-4F III-VI).

**Figure 4.**
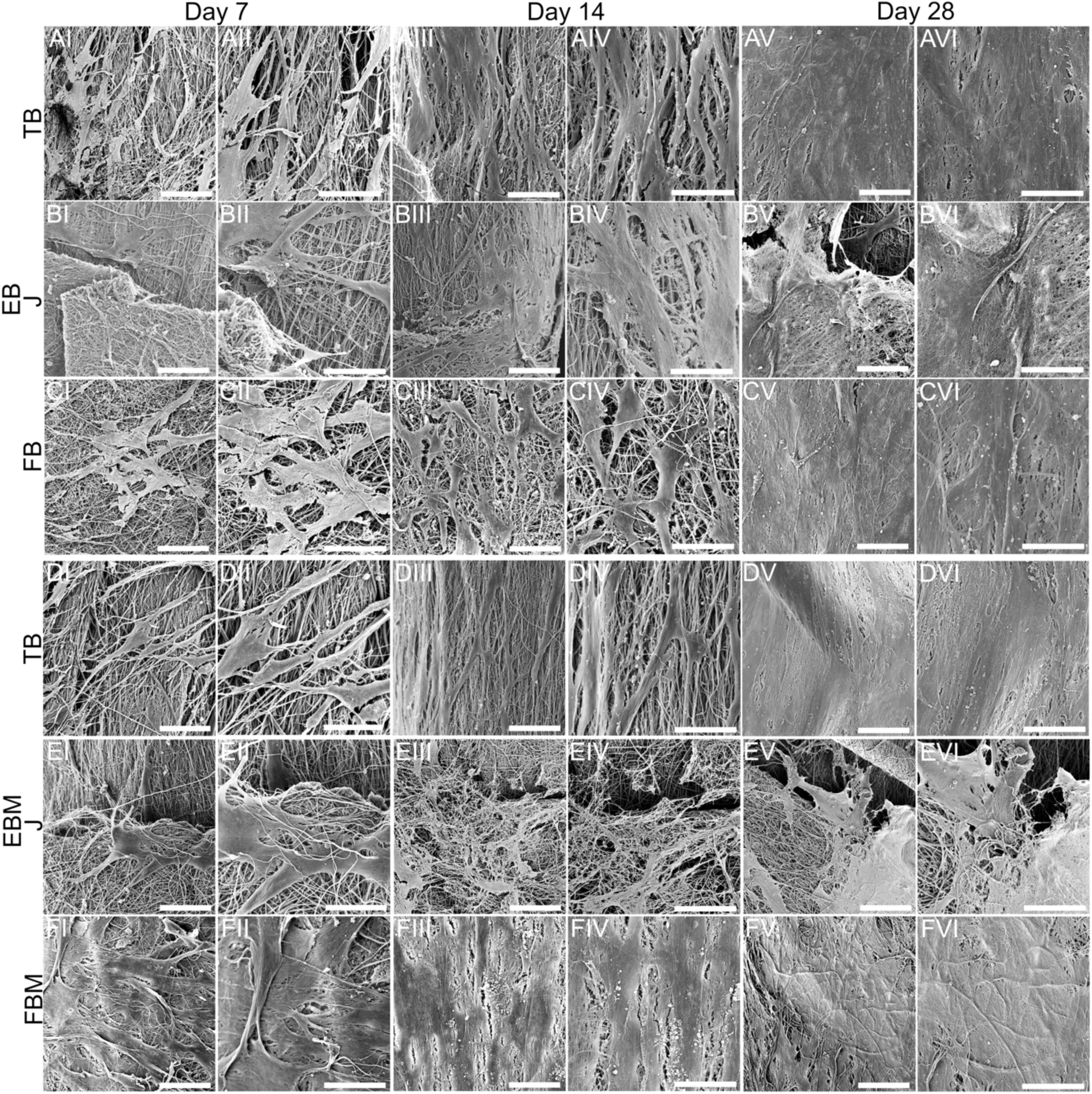
SEM investigation of scaffolds after the cell culture at the different time points. I, II) 7 days, III-IV) 14 days and V, VI) 28 days (images I, III, V: magnification = 500x, scale bar = 50 µm; images II, IV, VI: magnification = 1000x, scale bar = 10 µm). Images A-C show hMSCs grown on EB in their different regions: A) TB, B) J and C) FB. Images D-F show hMSCs grown on EBM in their different regions: A) TB, B) J and C) FB.

In the FB/FBM regions, hMSCs showed a clear tendency to modify their alignment moving towards an axial orientation (Figure 4BIII-4BVI, 4CIII-4CVI). This was confirmed by the orientation analysis of hMSCs actin filaments of cells obtained via confocal images (see Figure 5, Figure S3 and Table S9).

**Figure 5.**
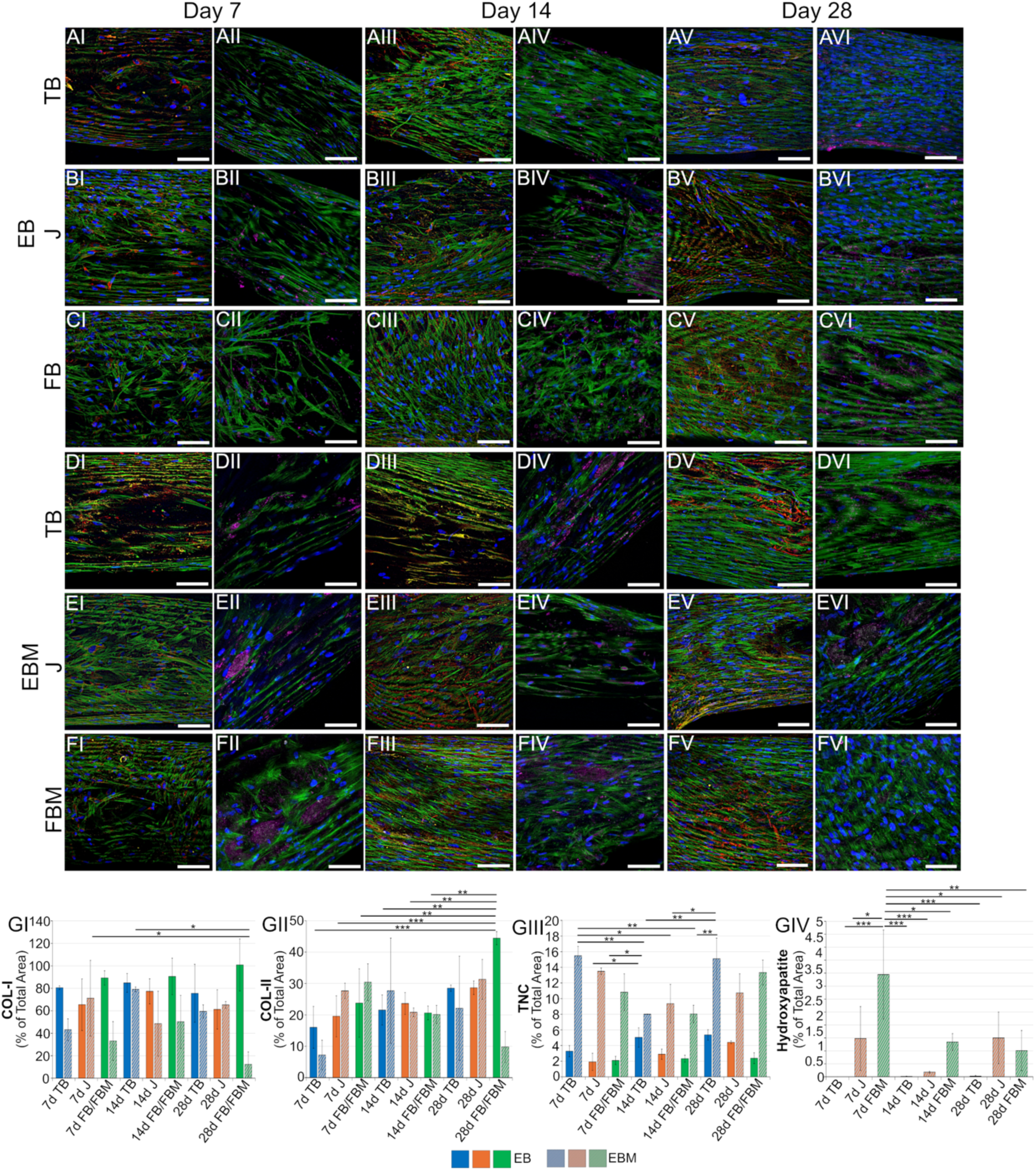
Maximum intensity projection of confocal fluorescent images of hMSCs on scaffolds at the different time points. For images I, III, V colors represent: blue = nuclei, green = F-actin, red = collagen type I (COL-I), yellow = tenascin-c (TNC); for images II, IV, VI: blue = nuclei, green = F-actin, purple = collagen type II (COL-II), grey = hydroxyapatite (scale bar = 100 µm) (see Figure S1 and S2 for the separate channels). Images A-C show EB scaffolds at 7, 14 and 28 days of culture with A) TB, B) J and C) FB sides respectively, while D-F show EBM scaffolds (same times points) with D) TB, E) J and F) FBM sides respectively. G) Quantification of matrix component (expressed in percentage of total cell area) in the different regions of EB and EBM at the different time points: GI) COL-I, GII) COL-II, GIII) TNC and GIV) hydroxyapatite via OsteoImage. The significance of differences of ECM quantifications on EB and EBM at the different time points of culture (7, 14 and 28 days) and regions, assessed with a one-way ANOVA followed by a Tukey post hoc (ns p>0.05, *p≤0.05, **p≤0.01, ***p≤0.001, ****p≤0.0001).

Starting from day 7 at the FB and FBM regions of EB and EBM, cells showed a preferential orientation of actin filaments in the range 0° - 18° of 38 ± 2% (FB) and of 35 ± 5% (FBM). After 28 days of culture, actin filaments orientation increased in the range 0°-18° reaching values of 49 ± 7% (FB) and of 59 ± 11% (FBM). The actin axial orientation of hMSC was probably caused by the inner layers of axially aligned nanofibers. Those layers, for the shrinkage of their nanofibers, occurred after crosslinking and produced an axial pre-strain of the FB/FBM regions (Figure 2CIII, 2EIII and 2FIII). Similarly in the J and TB regions, hMSCs increased their axial elongation appearing progressively thinner than the ones in the FB/FBM regions covering them with ECM (Figure 4A-4FV, 4VI, Figure 5, Figure S3 and Table S9). The same trend was confirmed also looking at the cells’ nuclei that showed an axial orientation tendency while visually maintaining a higher circularity in the FB and FBM regions, compared with the more elongated and thinner in the TB/J ones, at the different time points (Figure 5, Figure S1 and Figure S2). This resembled the different cellular populations in the natural tissue.[69]

To investigate the enthesis-specific ECM produced by hMSCs, immunostaining was performed targeting for some of the typical components of the enthesis tissue: collagen type I (COL-I) and type II (COL-II), tenascin c (TNC) and the mineral content (Figure 5, **Figure S1**, **Figure S2, Table S10** and **Table S11**).[7]

Confocal images confirmed the progressive proliferation of cells, previously seen with SEM (Figure 4), and the increasing deposition of ECM in the different regions of scaffolds. Specifically, the quantification analysis revealed that a significant amount of COL-I, the major component of both tendon, enthesis and bone tissue,[7,8,13] was produced by cells on all the regions of EB and EBM scaffolds, with a higher percentage of area on the non-mineralized ones. For EB samples, starting from day 7, more than the 80% of the cell area was covered by COL-I reaching after 28 days most of their surface (Figure 5, Figure 5GI, Figure S1AIII – S1CIII, S1AVIII – S1CVIII, S1AXIII – S1CXIII and Figure S2AIII - S2CIII, S2AIX - S2CIX, S2AXV S2CXV). A reduced amount of COL-I was produced on the mineralized samples EBM (40 - 60% of their area), with a notable drop after 28 days at the FBM region. It is worth mentioning that TB and FB were the regions with the higher percentage of area covered by COL-I, while being less present on the J region as in the natural tissue counterpart. Also COL-II, the typical protein present in both mineralized and non-mineralized fibrocartilage,[8,13] was deposited on scaffolds, reaching maximum values around the 40% of total area, and with a tendency to be homogeneously present on all the different regions of EB and EBM, with a particular focus on the J, FB and FBM ones. This again was similar to the natural enthesis, where COL-II is more expressed at the non-mineralized and mineralized fibrocartilage region. Interestingly, while EB showed a progressive increment in the coverage by COL-II of its different regions. For this protein, after 28 days a relevant drop in its presence was detected on the FBM region (Figure 5, Figure 5GII, Figure S1AIV – S1CIV, S1AIX – S1CIX, S1AXIV – S1CXIV and Figure S2AIV S2CIV, S2AX - S2CX, S2AXVI - S2CXVI, Table S10 and S11). This behavior of progressive decrease in COL-I and COL-II production by hMSCs on FBM regions of EBM scaffolds seems to follow the progressive loss of mineral content by their FBM region (Figure 5, Figure 5GIV and Figure S2AVI - S2CVI, S2AXII - S2CXII, S2AXVIII - S2CXVIII, Table S10 - S11). In fact, the mineral content at FBM passed from mean values of 3.5 % at day 7 to cover a surface area of less than 1% at day 28. A slight presence of mineral was detected on the J region of EBM (variable and with mean values around 1%), probably caused by a partial filtration of the mineral solution at this region during the mineralization phase (Figure 5GIV, Table S10 - S11). In terms of percentage of surface area covered by TNC, a typical protein released by cells in the early stages of tendon and enthesis genesis and healing,[70] a higher percentage seemed to be present on EBM regions (around 8 - 16%) compared with the EB ones (around 6 - 2 %) with a significant higher presence in the TB regions of both scaffolds (Figure 5GIII, Table S10 - S11). Despite region-specific deposition of COL-I, COL-II and TNC on EB and EBM, all these proteins were continuously distributed along the length of scaffolds, ensuring the continuity of the ECM along the entire length of the scaffolds’ surface.

#### 2.3.2. Enthesis Inspired Scaffolds Drive hMSCs Differentiation and ECM Production

The evolution of the phenotypic state of the cells, grown on EB (i.e. non mineralized FB side) and EBM (i.e. mineralized FB side) scaffolds, as well as the protein expression of ECM, were monitored after 7, 14 and 28 days of culture in basic media (**Figure 6**). This investigation was carried out to evaluate the effect of nano-mineralization to drive the stem cell fate and ECM production. The choice of media was guided by the need to see how the region-specific architecture and the PLLA/COL-I blend were able to self-guide the stem cell fate. Both the gene and protein expression were studied on the entire population of cells within the scaffolds. To have an overview of the expression of both tenogenic, chondrogenic and osteogenic markers typical of enthesis,[11] specific genes were investigated: *Col-I* as prevalent ECM component for all the four enthesis regions; scleraxis (*Scx*) and *Tnc* as tendon markers; SRY-Box Transcription Factor 9 (*Sox9*) and *Col-II* as chondrogenic markers; osteopontin (*Opn*), osterix (*Osx*) and runt-related transcription factor 2 (*Runx2*) as mineralized fibrocartilage and bone markers. From the protein expression level instead, COL-I, COL-II, TNC and tenomodulin (TNMD) were analyzed.

**Figure 6.**
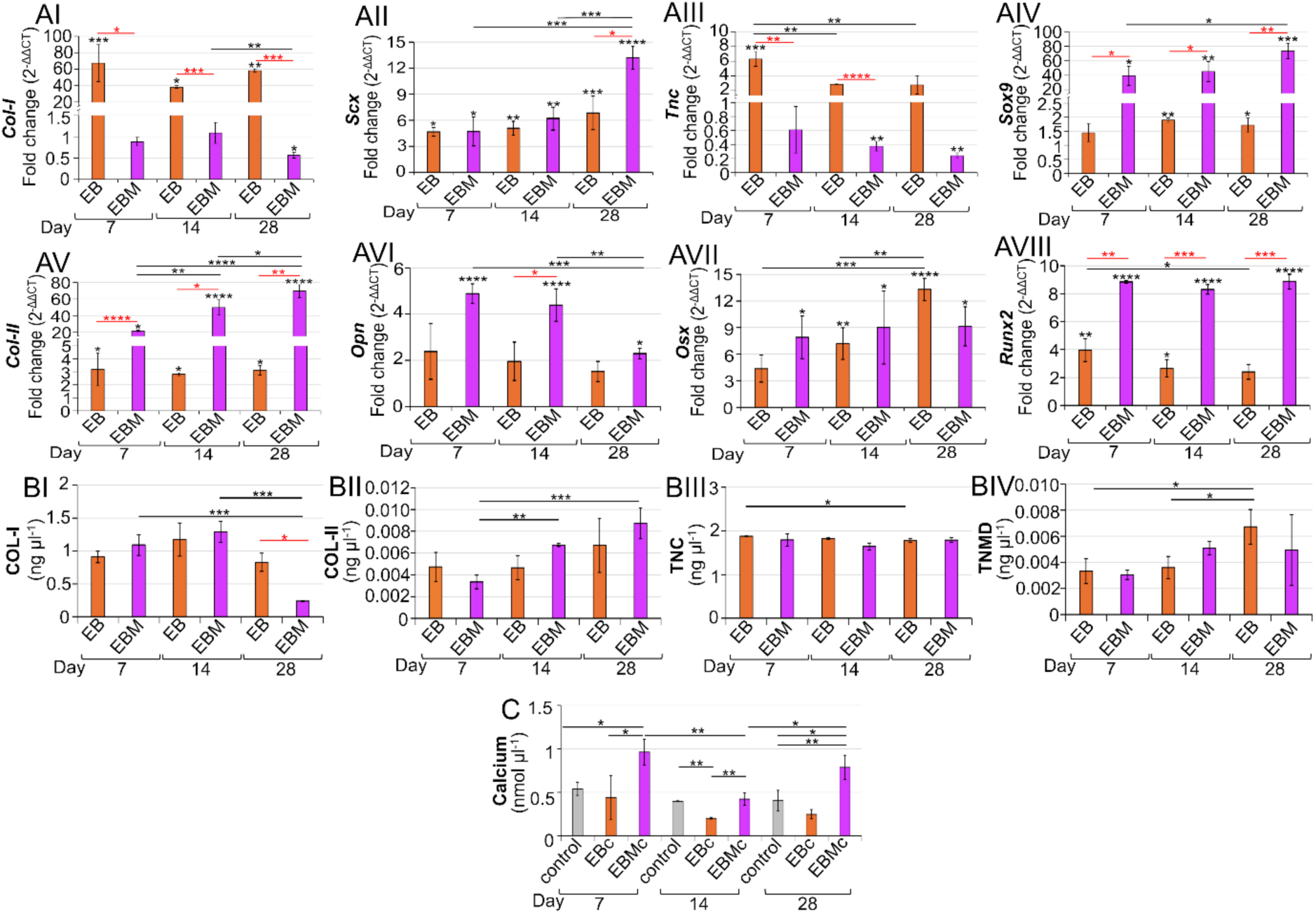
Gene and protein expression of hMSCs on scaffolds at the different time points (7, 14 and 28 days) and calcium assay. A) Gene expression: AI) collagen type I (*Col-I*), AII) scleraxis (*Scx*), AIII) tenascin c (*Tnc*), AIV) SRY-Box Transcription Factor 9 (*Sox9*), AV) collagen type II (*Col-I*), AVI) osteopontin (*Opn*), AVII) osterix, AVIII) runt-related transcription factor 2 (*Runx2*). B) ELISA test results: BI) collagen type I (COL-I), BII) collagen type II (COL-II), BIII) tenascin (TNC) and BIV) tenomodulin (TNMD). C) Comparison between the calcium assay carried out between the control medium, EB and EBM at different time points of culture with hMSCs (7, 14 and 28 days). The significance of differences of each gene, protein and each of the single family for the calcium assay (control medium, EB and EBM) at the different time points of culture (7, 14 and 28 days) assessed with a one-way ANOVA followed by a Tukey post hoc (ns p>0.05, *p≤0.05, **p≤0.01, ***p≤0.001, ****p≤0.0001, black asterisks; for RT-qPCR over the bars the significance of differences compared to day 1 of EB and EBM scaffolds are reported). The significance of differences between the same gene and protein expressed by hMSCs on EB and EBM at the different time points of culture (7, 14 and 28 days) assessed with an unpaired parametric t-test with Welch’s correction (ns p>0.05, *p≤0.05, **p≤0.01, ***p≤0.001, ****p≤0.0001, red asterisks).A one-way ANOVA followed by a Tukey post hoc between the different families of the calcium assay each time point was also performed.

In terms of *Col-I*, hMSCs cultured on EB scaffolds expressed significantly higher gene levels (around 60 fold change) compared with the EBM ones, where it resulted down-regulated (around 0.5 - 1 fold change) at all time points (Figure 6AI, Table S12 - S14). Similar levels of protein expressed up to day 14 (day 14: EB_COL-I_ = 1.17 ± 0.25 ng µl^-1^, EBM_COL-I_ = 1.29 ± 0.16 ng µl^-1^), while at day 28 the protein levels were reduced for both the scaffolds families but with a statistically significant reduction for the EBM ones (day 28: EB_COL-I_ = 0.83 ± 0.14 ng µl^-1^, EBM_COL-I_ = 0.24 ± 0.01 ng µl^-1^) (Figure 6BI, Table S15 - S17). Focusing on tendon-specific markers, the expression of *Scx*, gene related to the tendon development and involved in the regulation of *Col-I* and *Tnmd*,[71] and *Tnc* were investigated. In both EB and EBM *Scx* showed a progressive increment with respect of day one for both hMSCs cultures with a significant different increment at day 28 for the EBM compared with the EB at the same time point (mean fold change day 28: EB*_scx_* = 6.8; EBM*_scx_* = 13) (Figure 6AII, Table S12 - S14). The increment of *Scx* was in accordance with the progressive increment of COL-I production indicating also a possible differentiation of hMSCs into tenocytes.[72] *Tnc*, being an early marker for tenogenesis, was significantly up-regulated in the early stage of culture for EB, passing from 6.3 of day 7 to around 2.8 fold changes at day 14 - 28 while EBM was down-regulated at each time point (from 0.6 fold change of day 7 to 0.2 fold change at day 28) (Figure 6AIII, Table S12 - S14). Interestingly, the levels of TNC protein were similar and stable for both EB and EBM at all the time points (with a mean value of around 1.78 ng µl^-1^) (Figure 6BIII, Table S15 - S17). The expression of TNMD was also investigated showing a progressive increment at the later time points, being this protein related to the development and maturation of adult tendons.[73] TNMD expression was slightly higher on EB (day 7: EB_TNMD_ = 0.003 ng µl^-1^, day 28: EB_TNMD_ = 0.007 ng µl^-1^) compared to EBM (day 7: EBM_TNMD_ = 0.003 ng µl^-1^, day 28: EBM_TNMD_ = 0.005 ng µl^-1^) (Figure 6BIV, Table S15 - S17). Being TNMD involved in the regulation of the composition and organization of the extracellular matrix in tendons, its increment could positively influence the synthesis of collagen, improving the strength of ECM.

EBM scaffolds showed a significant increase of the chondrogenic transcription factor *Sox9* and *Col-II* in the EBM scaffolds (up to EBM*_sox9_* = 74 fold change, EBM*_col-II_* = 70 fold change at day 28) compared to the EB ones (up to EB*_sox9_* = 1.7 fold change; EB*_col-II_* = 3.1fold change at day 28) (Figure 6AIV and 6AV and Tables S12 - S14). Specifically, *Sox9* is an important transcription factor that guides the differentiation of hMSCs in chondrocytes while COL-II is the prevalent collagen at the non-mineralized and mineralized fibrocartilage region.[7] Despite the significant higher gene expression of *Col-II* in EBM, the protein levels were incremental in both scaffolds’ families after 28 days of culture (EB_COL-II_ = 0.007 ± 0.002 ng µl^-1^, day 28: EBM_COL-II_ = 0.009 ± 0.001 ng µl^-1^).

The expression of *Opn* was also investigated being a protein present at the mineralized fibrocartilage of the enthesis and plays an important role in the regulation of mineralization.[74] *Opn* is known to be a protein that mediates the bone regeneration via mechanical stress caused by collagen fibrils sliding,[75] a fundamental scenario when associated with electrospun scaffolds. In light of this, considering also the presence of the nano-mineralization, hMSCs on EBM scaffolds showed higher and significant values compared to the EB ones at 7 and 14 days (day 7: EB*_Opn_* = 2.4 and EBM*_Opn_* = 5.9 fold change; day 14: EB*_Opn_* = 2.0 and EBM*_Opn_* = 4.4 fold change) reducing its expression at 28 days of culture (EB*_Opn_* = 1.5 and EBM*_Opn_* = 2.3 fold change) (Figure 6AVI, Tables S12 - S14).

The *Osx* gene was studied as osteoblast-specific transcription factor involved in the activation of multiple genes during the differentiation of hMSCs in mature osteoblasts and osteocytes.[76] Here, EB showed a progressive increment of gene expression reaching after 28 days of culture higher values compared to hMSCs cultured in EBM (EB*_Osx_* = 13 and EBM*_Osx_* = 9.2 fold change) (Figure 6AVII, Tables S12 - S14). As last osteogenic marker, the expression of *Runx2* was investigated. *Runx2* is a transcription factor that regulates the differentiation of chondrocytes and stem cells in osteoblasts.[77] The significant and constant expression of *Runx2* by hMSCs cultured on EBM compared to the ones grown on EB (mean values of EBM*_runx2_* around 9 fold change per each time point; EB*_Runx2_* from 3.4 down to 2.4 fold change at day 28) (Figure 6AVIII and Tables S12 - S14) suggests the tendency of EBM to drive hMSCs in a progressive ossification of scaffolds.

This tendency was further confirmed by the calcium assay carried out between the EB and EBM cultures at the different time points (Figure 6C and **Tables S18** and **S19**). In fact, for the EBM scaffolds a significant amount of calcium was released starting from day 7 (day 7: EB_calcium_ = 0.44 ± 0.25 nmol μl^-1^, EBM_calcium_ = 0.96 ± 0.15 nmol μl^-1^). This higher amount was probably caused by the release of the nano-mineralization from the FBM side that, as also visible from the OsteoImage (Figure 5), was also detected at the J level. To confirm this mineral production, an EDS analysis was also performed. While ECM production was clearly visible by SEM images (Figure 4), the small amount of mineral content was more difficult be detected (**Figure S4** and **Table S20**).

After a significant reduction at day 14, at day 28 EBM showed a significant increment in the amount of calcium released while EB maintained about the same calcium values in the medium. The reason for the growth might be due to the calcium produced by the differentiated hMSCs in osteogenic lineage as visible from gene expression analysis (Figure 6AVI-6AVIII).

### 2.4. Multiscale Synchrotron X-ray In-situ Tensile Tests and Digital Volume Correlation

Considering the RT-qPCR and hMSCs cultures results, the best candidates as enthesis fascicle-inspired scaffolds resulted in the EB ones. To deeply investigate their full-field strain distribution from the nano-up to the micro-scale, cutting-edge multiscale synchrotron X-ray *in situ* stepwise tensile tests followed by a DVC analysis were carried out, by using a custom-made *in situ* mechanical tensile tester (**Figure S5**). A first group of *in situ* tests, zooming-in at the TB, J and FB regions of EB was performed via SnCT (voxel size = 300 nm; ID16B beamline, ESRF, Grenoble, FR). To have an overview of the micromechanics of the whole EB, SµCT was also used (voxel size = 650 µm; ID19 beamline, ESRF, Grenoble, FR). The strain steps were different for each sample and beamline, justified by the fact that the loading was interrupted at different points attempting to catch significant changes in the macroscopic mechanical response (force-strain curve) supported by visual changes in the imaged sample during acquisition. Four loading steps were performed during the SµCT in-situ test, while the smaller field of view of the SnCT limited the visual inspection of the imaged samples during acquisition, resulting in performing a single loading step before failure (Figure 7AIV, 7BIV, 7CIV and 7DVII). A DVC analysis was successfully carried out on all samples at the different beamlines (**Figure 7**, **Figure 8** and **Figure 9**), detecting the evolution of the local strain distributions of nanofibers, caused by their reorganization along with the application of the external apparent strain steps. For all DVC measurements, the uncertainties at zero strain (DVC between successive reference scans without loading the samples, see Experimental Section/Methods) resulted around one order of magnitude lower compared with the computed strains during the *in situ* tests (Table S21).

**Figure 7.**
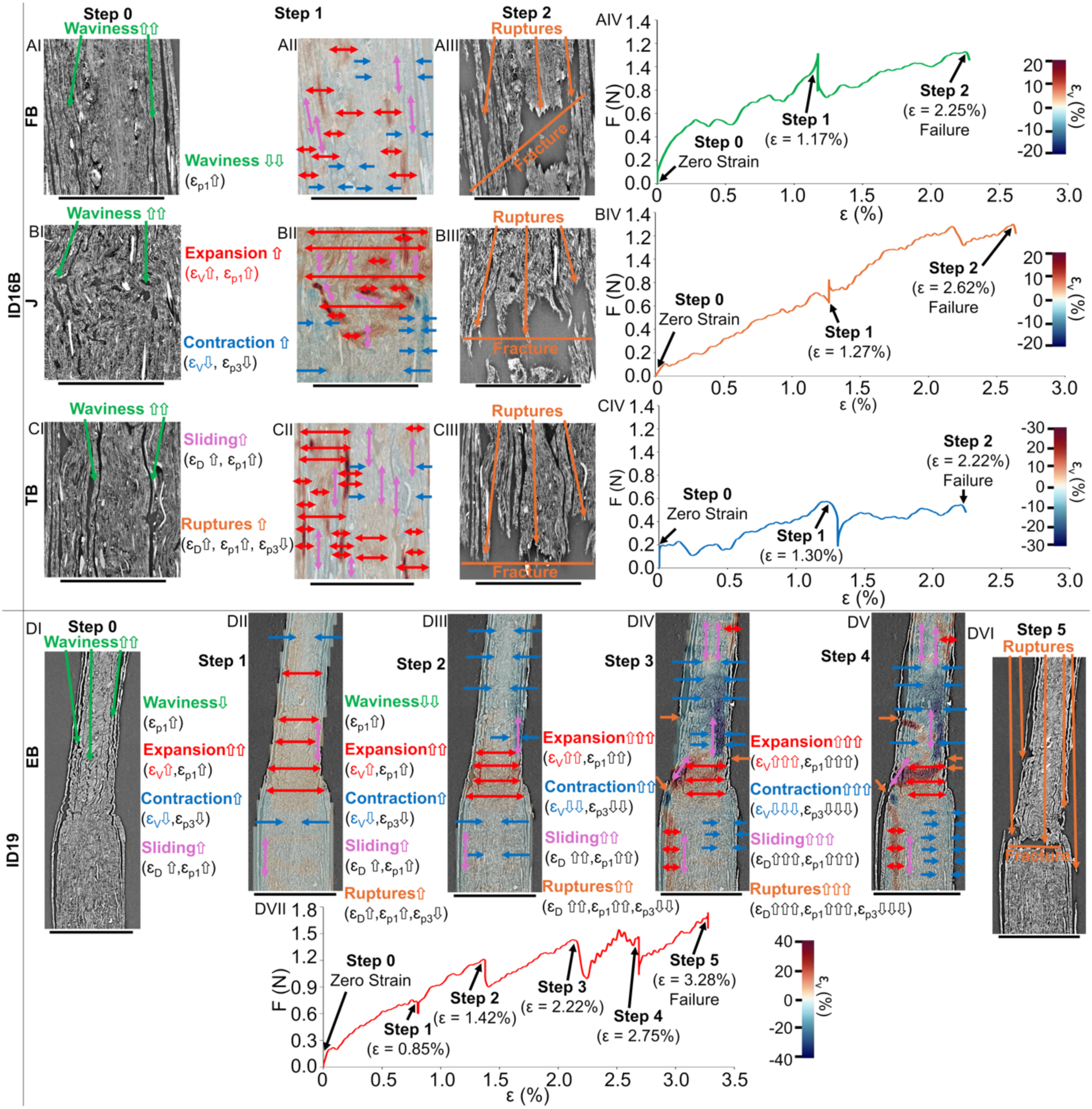
Macroscopic force-strain curves and region-specific DVC strain evolution fields of scaffolds during the multiscale synchrotron X-ray in situ tensile tests at ID16B (SnCT) and ID19 (SµCT) (arrows: red = volumetric expansion, blue = volumetric contraction, pink = sliding of nanofibers/layers). Evolution of the volumetric strain (ε_v_) fields as a function of the other strain components in internal axial reconstructed slices of the three regions of EB at ID16B: A) FB; B) J; C) TB. I) step 0: zero-strain; II) step 1: strain step; III) step 2: failure (voxel size = 300 nm; scale bar = 300 µm); IV) *in situ* force-strain curves. D) Evolution of the volumetric strain (ε_V_) as a function of the other strain components in internal axial reconstructed slices of the whole EB at ID19: I) step 0: zero-strain; II) step 1; III) step 2; IV) step 3; V) step 4; VI) step 5 (voxel size = 650 nm; scale bar = 600 µm); VII) *in situ* force-strain curve.

**Figure 8.**
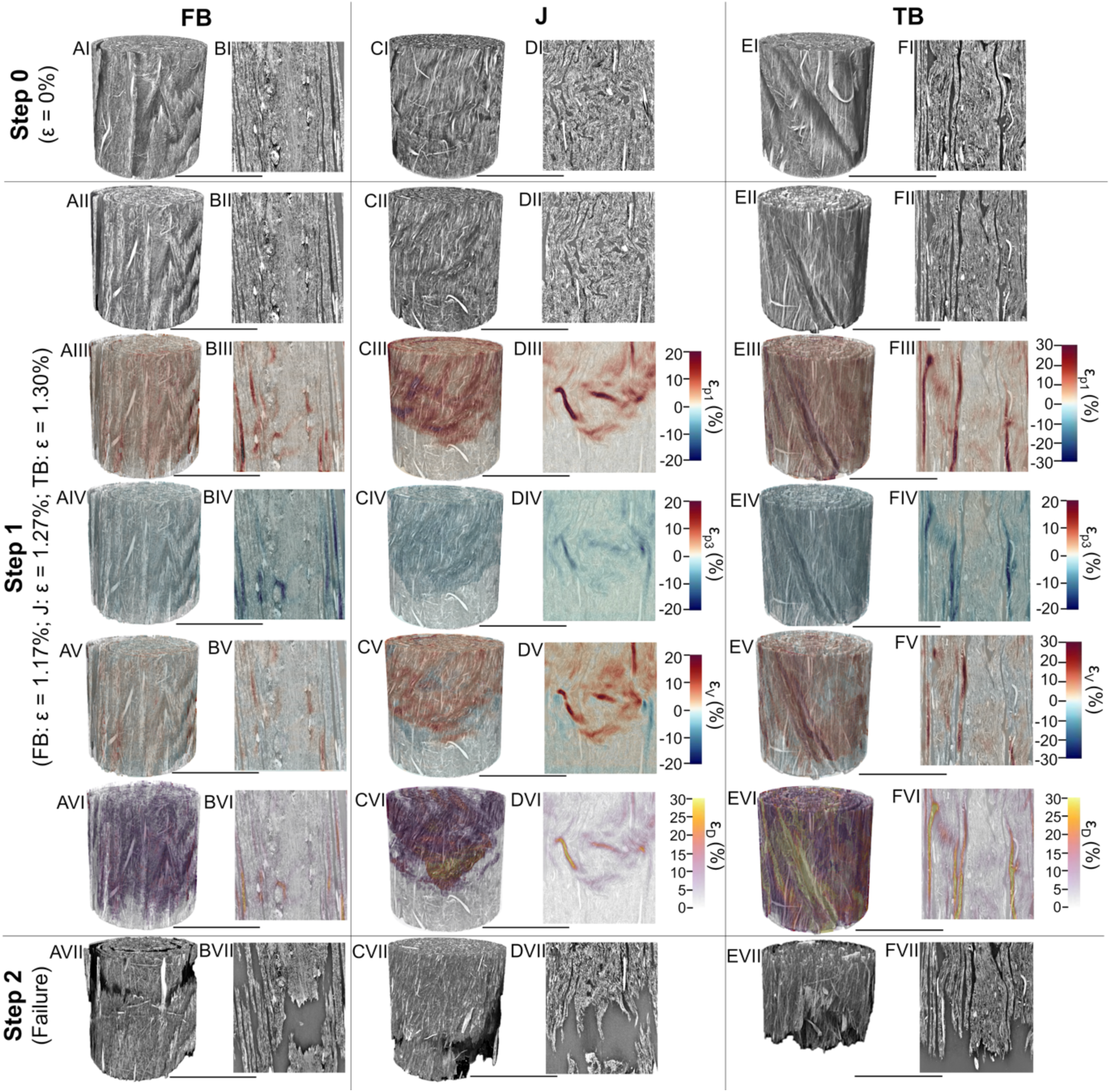
DVC strain components during the *in situ* stepwise tensile tests of the different parts of EB acquired at ID16B (voxel size = 300 nm, scale bar = 300 µm). AI, CI, EI) volume rendering of FB, J and TB; BI, DI, and FI) axial reconstructed slice in the volume of the corresponding scaffold at zero strain. Volume rendering and internal axial slice of scaffolds at the apparent strain step 1: II) reconstructed grayscale images; III) principal strain one (ε_p1_; maximum principal strain); IV) principal strain three (ε_p3_; minimum principal strain); V) volumetric strain (ε_V_); VI) deviatoric strain (ε_D_). VII) Volume rendering and 2D internal slice of scaffolds after failure.

**Figure 9.**
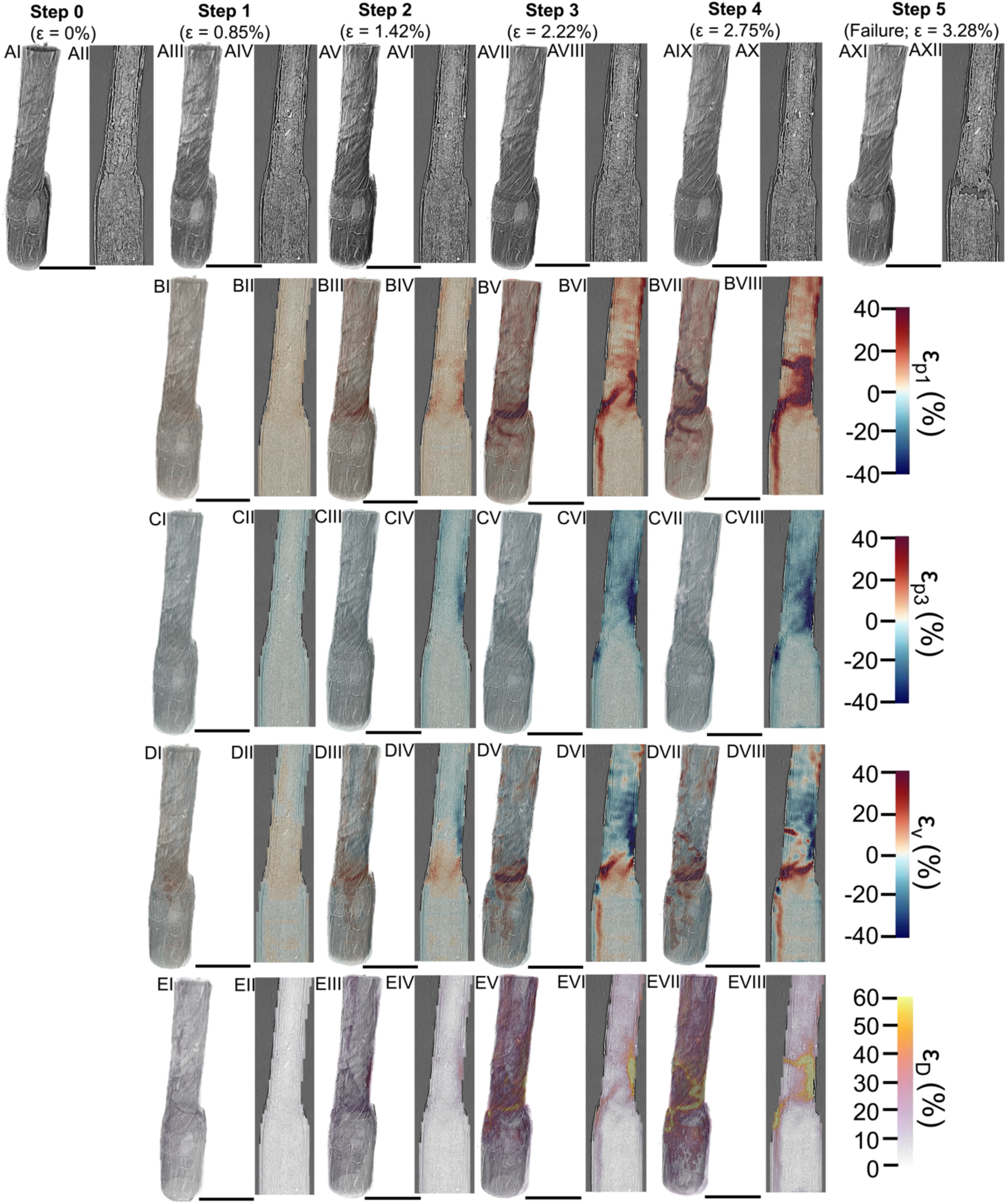
Evolution of the DVC strain components during the *in situ* stepwise tensile test of a whole EB acquired at ID19 (voxel size = 650 nm, scale bar = scale bar = 600 µm). I, II) step 0 (zero-strain); III-IV) step 1; V-VI) step 2; VII-VIII) step 3; IX-X) step 4; XI-XII) step 5 (failure). Per each strain step: A) volume rendering and axial reconstructed slices; B) principal strain one (ε_p1_; maximum principal strain); C) principal strain three (ε_p3_; minimum principal strain); D) volumetric strain (ε_V_); E) deviatoric strain (ε_D_).

Zooming into the nanoscale with the SnCT *in situ* tests, at the step 0 (i.e. zero strain state, intact sample) the nanofibers/layers of all scaffolds showed a wavy shape induced by the shrinkage occurred after their removal from the drum collector and the subsequent crosslinking (Figure 7AI, BI, CI). At strain step 1 (ε_FB_step1_ = 1.17 %, F_FB_step1_ = 1.10 N; ε_TB_step1_ = 1.30 %, F_TB_step1_ = 0.47 N; ε_J_step1_ = 1.27 %, F_J_step1_ = 0.64 N), the nanofibers/layers of scaffolds were stretched mostly recovering their waviness (Figure 7AII-CII and Figure 8AII-FII).

Focusing on the FB sample (Figure 7A, Figure 8AII-AVI and 8BII-BVI and Table S21), the DVC analysis revealed that the maximum principal strains (ε_p1_) (i.e. axial tensile component of strain), resulted in a mean value of ε_p1_mean_ = 4.79 ± 6.77 %. These ε_p1_ strains caused parallel rearrangements/yields of groups of nanofibers, producing a mean value of minimum principal strains (ε_p3_) (i.e. compressive component of strain) of ε_p3_mean_ = - 3.76 ± 4.85 %. Being the aligned nanofibers stiffer and stronger than the random ones, the parallel stretching of these different layers of nanofibers inside FB caused the concurrent presence of volumetric expansion and contraction regions. This effect was highlighted by the relatively low volumetric strains (ε_V_) at FB (with a mean value of ε_V_mean_ = 0.68 ± 6.21 %). In these scaffolds, the reduced volumetric expansion was probably caused by the packing/damping effect produced on the layers of aligned nanofibers by the layers of random ones and by the higher numbers of molecular connections produced by the crosslinking (FB contains a higher number of nanofibers). This effect was also confirmed by the limited amount of sliding between the nanofibers and layers, highlighted by the deviatoric strain (ε_D_) (i.e. sliding component of strain) with a mean value of ε_D_mean_ = 6.21 ± 7.34 % (Figure 7AII, Figure 8AVI, 8BVI and Table S21). Passing at the TB side (Figure 7C, Figure 8E, 8F and Table S21), the mean values of ε_p1_mean_ = 6.71 ± 7.51 % (Figure 7CII, Figure 8EII-8FIII) resulted higher compared to FB. This can be explained by the fact that TB is fully made of axially aligned nanofibers with a higher possibility of sliding along each other, as also shown by the higher mean values of ε_D_mean_ = 7.91 ± 7.16 % (Figure 7CII, Figure 8EVI, 8FVI). Together with the axial stretching (ε_p1_), groups of nanofibers rearranged themselves (ε_p3_) (probably yielding in some cases), producing a mean value of ε_p3_mean_ = −5.33 ± 3.77 % (Figure 7CII, Figure 8EIV, 8FV). Notably, both in FB (ε_p3_max_ = 2.47 %) and more visible in TB (ε_p3_max_ = 3.29 %) locally ε_p3_ reached positive peak values of strain (Figure 8AIV, 8BIV, 8EIV, 8FIV). These peaks were caused by the volumetric expansion of the aligned nanofibers, produced after the recovery of their waviness, as detected also in the natural tendon tissue.[17,19] This behavior was further confirmed by the TB ε_V_mean_ = 1.74 ± 9.67 %, in correspondence to its volumetric expansion (Figure 7CII, Figure 8EV, 8FV).

Moving at EB and focusing on its junction (J), here the volumetric expansion of the conical region was magnified by reaching the highest mean values of volumetric strain ε_V_mean_ = 2.52 ± 3.25 %. Notably a highly heterogeneous strain distribution was measured with positive volumetric local strains at the TB side and negative volumetric local strains at the FB side with concentration of high expansion values at the conical region (Figure 7BII, Figure 8CV, 8DV). This effect was obtained thanks to the parallel presence of three factors: i) the difference in cross-section between FB and TB sides (D_FB_ ∼ 700 µm; D_TB_ ∼ 400 µm); ii) the abrupt passage between a region with layers of random/aligned nanofibers (FB) and a region of fully aligned nanofibers (TB) that caused also iii) the higher level of shrinkage of EB among the various regions (i.e. sk_EB_ ∼ 19%) (Figure 2, Figure 7BI, 7BII, Figure 8C, 8D and Video S1-S8). In particular, the effect of shrinkage concentrated a relevant waviness of nanofibers at the J side (Figure 7BI) that, under the effect of the axial stretching applied to reach step 1, maximized the presence of ε_p1_ at the conical region (and TB side of EB), reaching values of ε_p1_mean_ = 3.82 ± 4.42 % (Figure 8CIII, 8DIII). The development of relevant ε_p1_ was facilitated by the high sliding of nanofibers at the J side (ε_D_mean_ = 5.94 ± 4.46 %) localizing in particular in its central region (Figure 8CVI, 8DVI), maximizing the volumetric expansion of EB (Figure 7BII). This relevant production of ε_p1_/ε_D_ caused the yielding of groups of nanofibers (Figure 7BII and Figure 8CIV, 8DIV) with values of ε_p3_mean_ = −2.59 ± 2.27 %.

After step 1, samples were loaded up to failure that occurred at step 2 (ε_FB_step2_ = 2.25 %, F_FB_step2_ = 1.13 N; ε_TB_step2_ = 2.22 %, F_TB_step2_ = 0.54 N; ε_J_step1_ = 2.62 %, F_J_step2_ = 0.64 N) (Figure 7AIII, 7BIII, 7CIII and Figure 8AVII-8FVII).

Notably, the failure of all samples occurred before the yielding point and around the inflection point (i.e. 0.4 % s^-1^: IFF_FB_ = 1.4 ± 0.5 N, IFε_FB_ = 1.9 ± 0.9 %; IFF_TB_ = 1.0 ± 0.3 N, IFε_TB_ = 2.5 ± 1.3 %; IFF_EB_ = 0.8 ± 0.3 N, IFε_EB_ = 1.4 ± 0.4 %) (Table S4). This was probably caused by the mutual effect of a number of factors: i) the reduced dimensions of samples (i.e. gauge length of 4.5 mm) that partially caused their embrittlement; ii) the low strain rate of the *in situ* tester (i.e. 0.045 % s^-1^); iii) the radiation dose effect of the synchrotron X-rays that could potentially have caused some damages to the PLLA and COL-I molecules and the related progressive dehydration of the samples.[18,76] It is worth noting, that for both the tensile tests (Figure 3) and the *ex situ* preliminary tensile tests (Figure S5) a considerably larger macroscopic yielding of the samples was observed, with no scaffold failing at the junction. The effect of the radiation induced damage was further investigated on preliminary samples through a series of successive scans (see Experimental Section/Methods, DVC uncertainty (Table S21) and beam impact (**Figure S6** and **S7**). Interestingly, the macroscopic fracture behavior of the different scaffolds emerged by the local failure patterns of the different orientations of nanofibers, showing the typical trend of ductile and brittle materials. For FB, probably the fracture was initiated by the random layers of nanofibers (being this configuration notoriously more ductile than the aligned ones) producing a failure with a 45° orientation, typical of ductile materials (Figure 7AIII and Figure 8AVII, 8BVII). TB instead, being composed of axially aligned nanofibers, produced a fracture mode with a perpendicular orientation with respect to the loading orientation that is typical of brittle materials (Figure 7CIII and Figure 8EVII, 8FVII). A similar fracture mode occurred at J, demonstrated as a fracture started at the interface between J and FB but driven by groups of aligned nanofibers and clearly following the conical shape of the junction (Figure 7BIII and Figure 8CVII, 8DVII).

The SµCT *in situ* tensile test (Figure 7D and Figure 9) allowed the investigation of the whole strain distribution field on the whole EB sample. The waviness of nanofibers/layers was evident at step 0 (zero-strain step), with a higher level focused on the J and TB sides, and less evident at the FB, due to the relevant amount of packed nanofibers present in this region (Figure 7DI and Figure 9AI, 9AII). In this case, thanks to the increased field of view of the SµCT scans, it was possible to carry out four consecutive strain steps (step 1: ε__step1_ = 0.85 %, F__step1_ = 0.77 N; step 2: ε__step2_ = 1.42 %, F__step2_ = 0.91 N; step 3: ε__step3_ = 2.22 %, F__step3_ = 1.23 N; step 4: ε__step4_ = 2.75 %, F__step4_ = 1.32 N) before failure (step 5: ε__step5_ = 3.28 %, F__step5_ = 1.74 N).

In terms of ε_p1_, strains progressively increased in the three different regions of EB (Figure 9B and Table S20). The FB region showed the lowest mean strain values from step1 (ε_p1_mean_ = 1.49 ± 2.65 %) up to step 4 (ε_p1_mean_ = 5.59 ± 7.76 %) with some local concentration corresponding with a side of the scaffold. TB showed increased values of ε_p1_ from step 1 (ε_p1_mean_ = 2.15 ± 1.71 %) to step 4 (ε_p1_mean_ = 11.5 ± 11.8 %) with local concentration corresponding with the top side of J and close to the clamp. The J region revealed the highest values of ε_p1,_ due to the high shrunk nanofibers to be stretched, with local strain concentration distributed along its length starting from step 1 (ε_p1_mean_ = 3.06 ± 3.38 %) up to step 4 (ε_p1_mean_ = 4.4 ± 16.9 %). The waviness of nanofibers at the J region was mostly recovered starting from the step 3 (Figure 7D and Figure 9B). Concerning ε_p3,_ FB reported the lowest values among the three regions, with local minimum values in correspondence with the bottom side of J (step 1: ε_p3_mean_ = −1.25 ± 1.81 %; step 4: ε_p3_mean_ = −4.51 ± 5.75 %) (Figure 9C and Table S21). This was probably caused by: i) the reduced rearrangements of nanofibers for the high packing of this zone; ii) the volumetric contraction for the yields/fractures of some layers of nanofibers (see Figure 7DIV, 7DV and Figure 9CVI-9CVIII); iii) the local compressive regions produced by J at the interface with FB. Consequently, some positive local concentrations of ε_p3_ were detected, due to the attempt of local expansions of the aligned nanofibers that was dampened by the presence of the random layers of nanofibers.

TB resulted in the region with the highest mean values of ε_p3_ that were progressively magnified (step 1: ε_p3_mean_ = −2.40 ± 2.13 %; step 4: ε_p3_mean_ = −16.4 ± 9.55 %,) (Figure 9C and Table S20). This can be attributed to the increasing number of yielded/broken nanofibers and the progressive volumetric restriction by the Poisson effect particularly focalized in TB, close to the top side of J (Figure 9C). Moreover, similarly to the FB region some positive local concentration of ε_p3_ were detected supporting the local expansion of the aligned nanofibers during the stretching of EB (Table S20).

The J region highlighted the presence of ε_p3_ that supported its volumetric expansion (step 1: ε_p3_mean_ = −1.61 ± 2.26%; step 4: ε_p3_mean_ = −9.73 ± 11.3 %). In fact, due to the high initial shrinkage of this region, the first two steps showed lowers levels of ε_p3_mean_ compared to the TB region. However, starting from step 3 the mean value of ε_p3_ started to visibly increase in magnitude at the top and bottom regions of J, due to the progressive expansion of its central part (Figure 9CVI-9CVIII).

The structural evolution of EB during the tensile test produced a subsequent increment of sliding (ε_D_), mostly concentrated at the TB and J regions, and less evident at the FB side (Figure 9E and Table S21). The high friction, pack and the presentence of crosslinking, limited the increase of ε_D_ at the FB region being mostly manifested close to the J (step 1: ε_D_mean_ = 2.00 ± 2.76 %; step 4: ε_D_mean_ = 7.39 ± 8.34 %).

Despite the crosslinking, the low friction between the aligned nanofibers at the TB region caused the appearance of high values of ε_D_, in particular in the last two strain steps and close to the top of the J side (step 1: ε_D_mean_ = 3.33 ± 2.22 %; step 4: ε_D_mean_ = 21.0 ± 11.5 %).

Similar values of ε_D_ were measured at the J side (step 1: ε_D_mean_ = 3.41 ± 3.38 %; step 4: ε_D_mean_ = 17.6 ± 15.7 %), demonstrating a progressive localization of the deviatoric strain in a band at the bottom part of the junction.

Concerning the volumetric strain (Figure 7D, Figure 9E and Table S21), the FB side of EB reported mean values of ε_V_ negligible along the different strain steps, with positive and negative peaks caused by local expansions/contraction of its internal layers and the upper expansion of the J region (step 1: ε_V_mean_ = 0.20 ± 2.55 %; step 4: ε_V_mean_ = 0.46 ± 7.80 %).

The TB side of EB, despite some local expansions in close proximity of the junction and the clamp, showed a constant volumetric contraction (step 1: ε_V_mean_ = −0.31 ± 2.55 %step 4: ε_V_mean_ = −8.57 ± 17.7 %).

The junction instead, confirmed the volumetric expansion, already highlighted with the SnCT *in situ* experiment (step 1: ε_V_mean_ = 1.68 ± 3.51 %; step 4: ε_V_mean_ = 4.4 ± 16.9 %). Together with the evolution of the deviatoric strain at this region, the progressive volumetric expansion yielded the appearance of a dilative shear band with localized high level of strains at the bottom part of the junction.

As expected from the strain fields evolution, at the last step the failure occurred at the bottom side of J (Figure 7DVI and Figure 9AXI, 9XII). The fracture mode confirmed the rupture behavior previously seen on J at ID16B (Figure 7DVI and Figure 9AXI, 9AXII), driven by the local failure patterns of the nanofibers and their reorganization. As discussed above for the SnCT, on top of the reduced sample length, lower loading rate and radiation dose effect, this macroscopic failure can be partially attributed to the underlying structure of the scaffold. In fact, TB, J and FB are regions with different porosities and nanofibers’ organization. The progressive volumetric expansion of J caused an increase of the internal porosity resulting in a partial detachment between its aligned nanofiber layers. Moreover, the difference in cross-section between FB and TB sides (D_FB_ ∼ 700 µm; D_TB_ ∼ 400 µm) and the abrupt passage between a region with layers of random/aligned nanofibers (FB) and a region of fully aligned nanofibers (TB) caused a significant concentration of strains at their interface, making this zone ideal for the fracture.

The differences between the DVC strain levels calculated in the two beamlines, at relatively similar strain steps (step 1 for SnCT and step 2 for SµCT), can be attributed to the higher scanning resolution of SnCT that was essential to probe the complex underlying structure of nanofibers and allowed the measurement of their respective local strain distributions at the nano-level. Meanwhile, the relatively lower scanning resolution of the SµCT, despite not sufficient to resolve the smaller nanofibers, allowed the measurement of the strain distribution along the whole EB examining the full-field mechanical behavior of the wrapped layers of nanofiber mats constituting the scaffold.

It is also worth mentioning that the mean values of strain, obtained from the DVC analysis of the SµCT *in situ* test, were in the same range of the apparent strains (before yield) obtained during the mechanical tensile tests (at 0.4 % s^-1^) and close to the IFε values (Table S4).

## 3. Conclusion

We designed innovative electrospun enthesis fascicle-inspired scaffolds, made of a blend of high molecular weight PLLA and collagen type I, with region-specific nanofibers orientations able to faithfully mimic the morphology and the mechanical properties of the different regions of the enthesis tissue. The structure of scaffolds was investigated via SEM and multiscale synchrotron X-ray tomography allowing to faithfully validate the scaffolds morphological biomimicry. The hMSCs cultures both on the as spun (EB) and on nano-mineralized at the mineralized fibrocartilage region (EBM), highlighted by using SEM and confocal microscopy the cell proliferation and the production of a relevant amount of ECM on scaffolds. The RT-RT-qPCR and Elisa analysis confirmed a more balanced deposition of enthesis-specific markers on the scaffolds without nano-mineralization (EB) while the nano-mineralized ones (EBM) resulted more prone to a progressive ossification. In parallel, the full-field strain distribution of the best candidate scaffolds (EB) was successfully investigated both for scaffolds and mostly for electrospun materials, with a multiscale nano- to micro-synchrotron X-ray tomography *in situ* stepwise tensile tests coupled with Digital Volume Correlation. This cutting-edge investigation allowed, for the first time, visualizing and resolving the evolution of the full-field strain distribution of nanofibers inside electrospun materials, showing how they influenced their overall nano- and micro-mechanics. Finally, the multiscale DVC investigation, was able to document the distinctive auxetic behavior of the enthesis non-mineralized-inspired junction of these innovative scaffolds, confirming their unique morphological and mechanical biomimicry with the natural enthesis fascicles, paving the way for a new generation of biomimetic scaffolds for the regeneration of the enthesis tissue.

## 4. Experimental Section/Methods

### Materials

Medical grade poly(L lactic) acid (PLLA) (Purasorb PL18, Mw = 1.7-1.9 x10^5^ g mol^-1^, Corbion, Amsterdam, NL) and acid soluble collagen type I (Coll), extracted from bovine skin (Kensey Nash Corporation DSM Biomedical, Exton, USA) were used. 1,1,1,3,3,3-Hexafluoro-2-propanol (HFIP) (Sigma-Aldrich, Saint Louis, USA) was used as solvent. For the crosslinking protocol N-(3-Dimethylaminopropyl)-N’-ethylcarbodiimide hydrochloride (EDC), N-Hydroxysuccinimide (NHS) and ethanol (EtOH) (Sigma-Aldrich, Saint Louis, USA) were used as received. The following polymeric blend solution was used: PLLA/Coll-75/25 (w/w) prepared from a 15% (w/v) solution of PLLA and Coll dissolved in HFIP.

### Scaffolds Fabrication

An industrial electrospinning machine (Fluidnatek® LE-100, Bionicia, Valencia, Spain), equipped with a high-speed rotating drum collector (length = 300 mm, diameter = 200 mm) was used. To electrospun the nanofibers, an applied voltage of 24 kV (needles side) and a slightly negative voltage of −0.5 kV (collector side) were applied. To easily detach the nanofibers mats, the drum was covered with a sheet of polyethylene (PE) coated paper (Turconi S.p.A, Italy). The polymeric solution was delivered with three metallic needles (internal diameter = 0.51 mm, Hamilton, Romania), through PTFE tubes (Bola, Germany), using a feed rate of 0.5 mL h^−1^ controlled by three syringe pumps installed in the electrospinning system. The needles-collector distance was of 200 mm; the sliding spinneret with the needles had an excursion of 190 mm, with a sliding speed of 3600 mm min^−1^. The electrospinning session was carried out at a temperature of 25 °C a relative humidity of 25%. To produce the scaffolds the electrospinning process was carried out in two steps. At first, a mat of random nanofibers was obtained by electrospinning the PLLA/Coll solution for 3h with the drum rotating at 50 rpm (peripheral speed = 1 m s^−1^). The mat was cut in stripes along the axis of the drum and alternately removing them from it. This process allowed reproducing the random arrangement of the bone/mineralized fibrocartilage ECM.[10] Then, the PLLA/Coll solution was electrospun for additional 3h at 2000 rpm (peripheral speed = 21 m s^−1^). This allowed covering the random mats and the gaps, previously obtained, with a homogeneous mat of aligned nanofibers (along the circumference of the drum), while the gaps were fully covered by aligned nanofibers. The final mat was firstly horizontally cut in between of each stripe, previously defined, and then with circumferential cuts of 30 mm far from each other. The circumferential stripes so obtained were rolled up on the drum obtaining enthesis-inspired bundles (EB) with region specific diameters and nanofibers orientations: a portion with random + aligned nanofibers, resembling the mineralized fibrocartilage side (FB), and one of fully axially aligned nanofibers, reproducing the tendon side (TB) with a small conical junction, connecting FB and TB, resembling the non-mineralized fibrocartilage (J).[10] Moreover, the difference of diameters between FB (approximately 700 μm) and TB (approximately 400 μm) allowed, in between them, the production of a conical junction of aligned nanofibers of around 500 μm long (diameter around 600 µm).[10] Finally, to closely mimic the natural mineralized fibrocartilage-bone tissue side, the FB region of a family of bundles were mineralized with nano-hydroxyapatite crystals obtaining enthesis-inspired mineralized bundles (EBM) with a nano-mineralized fibrocartilage region (FBM). To comprehensively investigate the mechanical properties of bundles (see below), understanding the effects of the junctions in tuning the scaffolds mechanics, all the different regions (both mineralized and not) were also separately fabricated and tested.

### Collagen Crosslinking

To fix the collagen in the nanofibers, EB were immersed for 24 h at room temperature (RT) in a crosslinking solution of EDC and NHS 0.02 M in 95% ethanol, using a consolidated procedure.[78] Then specimens were immersed in phosphate buffer saline (PBS, 0.1 M, pH = 7.4) for 30 min, washed in distilled water for 2 hours (by changing water every 15 min) and dried under fume hood at RT for 24 h.

### Mineralization

To mineralize the crosslinked EBM, their random+aligned regions were immersed in a solution, based on a previously optimized adapted 5-time simulated body fluid (ionic concentration in solution: Na^+^ = 733.5 mM, Mg^2+^ = 7.5 mM, Ca^2+^ = 12.5 mM, Cl^-^ = 720 mM, HPO_4_^2-^ = 5 mM, HCO_3_^-^ = 21 mM),[79] and left for 1 h at 37 °C in an incubator. EBM were washed for 15 min in distilled water and left to dry under a fume hood for 24 h. To visualize the mineralized areas of EBM scaffold, 60 mM of alizarine red solution (pH 4.1 - 4.3) was placed on the FBM region of scaffold and incubated for 20 min at RT. The scaffold was finally washed for 15 min in distilled water and stored in PBS at 4°C until being imaged with a stereomicroscope (Nikon SMZ25, Leuven, BEL).

### Scanning Electron Microscopy

The morphology of EB and EBM as well as the samples after cell cultures were analyzed via SEM (IT200, JEOL Ltd, Tokyo, JAP) using a voltage of 10 KeV. Before being imaged, the cellularized samples were at first fixated in 2.5% Glutaraldehyde in 0.1M PBS (Sigma-Aldrich) and then washed three times for 15 min in 0.1 M PBS. After fixation, samples were placed for 1h in 1% Osmium tetroxide (OsO4) (Van Loenen Instruments, NL) in 0.1 M of PBS and then washed again three times for 15 min in 0.1 M of PBS. Samples were then dehydrated with a series of 30 min incubation steps on solutions with increasing concentration of EtOH (v/v) at 70%, 90%, 100%, 100%, washed two times for 30 min in 1,1,1,3,3,3,-Hexamethyldisilazane (DMHS) (Sigma-Aldrich) and left to air dry overnight. Before being imaged, each sample was sputter coated with gold (SC7620, Quorum, Laughton, UK). To investigate the mineral content on the FB and FBM regions of EB and EBM samples (before and after the different time points of hMSCs cultures), energy dispersive x-ray spectroscopy (EDS) mode of SEM was used with a voltage of 20 KeV. As control the analysis was carried out also on the J/TB regions. The mass percentage of the different elements were computed as mean ± standard deviation (SD) between three images of each region (i.e. FB or FBM and J/TB) of the same sample with a magnification of 1000x (for the investigation after mineralization 5000x images were acquired).

The open-source software ImageJ was used to measure the diameters of nanofibers and hydroxyapatite nanocrystals. Specifically for each region of EB and EBM, 200 diameters of the different features were measured and showed both as mean ± SD as well as diameter distribution (magnification = 5,000x).

The nanofibers’ orientation on the surface of scaffolds was investigated with the Directionality plugin of Fiji.[80] This approach allowed the quantification of the number of nanofibers within a given angle range from the axis, using a Local Gradient Orientation method, following a previously validated procedure.[81] The analysis was performed along the different regions of EB and EBM. The analysis was performed on five images (magnification = 5,000x) along the scaffold’s axis and the results reported as mean ± SD between five images per each region. Moreover, also the diameter distribution of the FBM nanofibers and their mineral crystals were calculated with the same method previously described.

### Mechanical tensile tests

The mechanical tensile characterization of electrospun bundles (i.e. TB, FB, MFB, EB, EBM; n = 5 per category) were tested with a mechanical testing machine (Electroforce 3200, TA Instruments, New Castle, USA) with a 45 N load cell (TA Instruments, New Castle, USA). The machine worked at RT and under displacement control using a monotonic ramp to failure with three different strain rates: 0.4 % s^−1^ (consistent with a quasi-static task and useful to setup the *in situ* experiments); 10 % s^−1^ and 100 % s^−1^ (to simulate a physiological motor task and a worst case scenario of failure).[54–56] Before the crosslinking, each sample had a total length (l) of 40 mm of which 20 mm as gauge length (l_0_) and 10 mm per each side for clamping them. After the crosslinking and mineralization, all samples shrunk depending by the nanofibers’ arrangement in the different regions. The total length after shrinkage (l_sk_) was measured per each specimen and the percentage of shrinkage (sk) was calculated per each sample family as follows:

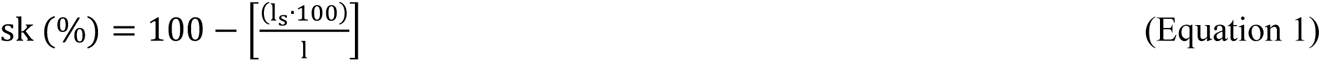

Then l_0_ was maintained at 20 mm (leaving for the gripping side around 5 mm per each side). To evaluate the junction mechanics, for EB and EBM the junction was placed in the middle of the gauge length (with 10 mm of TB + J and 10 mm of FB or FBM). The weight of the samples (w) was calculated using a precision balance (MG214Ai-M, VWR International, Radnor, USA) and expressed as mean ± SD of three measurements. To hydrate the samples before the test, they were immersed 2 min in PBS (Sigma-Aldrich, the Netherlands).

The force-displacement curves were converted to stress-strain graphs using two different approaches. In the first one, the apparent stress (α_A_) was calculated by dividing the force by the cross-sectional area of the specimen measured before the test (as mean ± SD between ten measurements along the gauge length of the scaffold) whereas in the second one, the net stress (α_N_), was used to determine the mechanical properties of the specimen excluding the contribution of its porosity.

The net stress was calculated by dividing the apparent stress by the volume fraction (v) of the specimens. The volume fraction was calculated by using the equation:

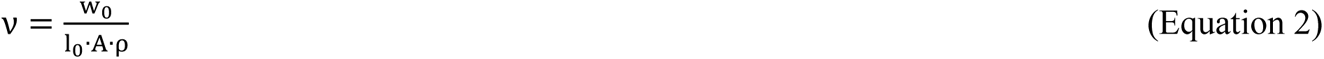

Where (w_0_) is the weight of the gauge length of the specimen, (l_0_) is gauge length of the specimen, (A) is the cross-sectional area of the specimen, ρ is the density that was calculated in two different ways depending on the mineral content. Specifically, w_0_ was calculated as follows:

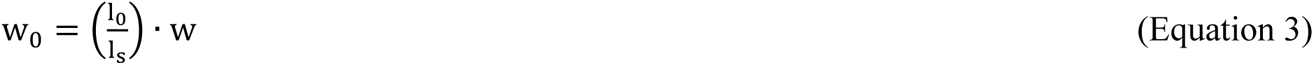

Similarly, for the calculation of the weight of the gauge length of mineralized samples the procedure was the same but adding to w the weight of the mineral content (w_M_).

For FB, TB, EB, ρ_B_ is the density of the bundle’s blend constituent materials that, considering the density of PLLA (ρ_PLLA_ = 1.25 g cm^-3^) and of the Coll (ρ_Coll_ = 1.34 g cm^-3^), resulted in:

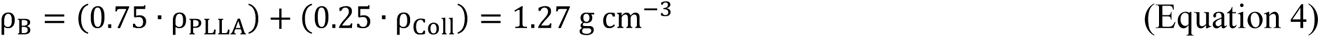

For FBM and EBM instead, the density (ρBM) was calculated as follows:

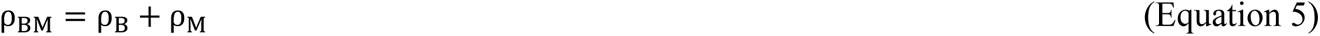

Where ρ_M_ is the density of the mineral component that was exterminated per each mineralized specimen as:

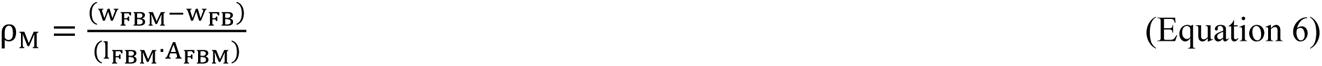

Where (w_FBM_) is the weight of the mineralized fibrocartilage side after mineralization, (w_FB_) is the weight of the mineralized fibrocartilage side before mineralization, (l_FBM_) and (A_FBM_) are the length and area of the mineralized fibrocartilage side after mineralization.

The following indicators were considered: yield Force (F_Y_), yield stress (σ_Y_), yield strain (ε_Y_), elastic modulus (E), stiffness (K), failure force (F_F_), failure stress (σ_F_), failure strain (ε_F_), unit work to yield (W_Y_), unit work to failure (W_F_). The W_Y_ and W_F_ were calculated using the trapezoid method.

The volume fraction also allowed the calculation of the percentage porosity (P) of bundles as follows:

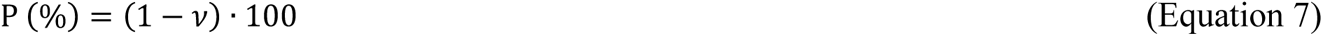

The inflection point value of scaffolds was evaluated using a previously validated method.[48,65] This is the point, in the linear region of the stress-strain curve, where materials transit from strain-stiffening to strain-softening. In brief, a MATLAB script was adopted, based on the csaps function, to fit a cubic smoothing spline to the mechanical data. A smoothing parameter in the csaps routine was set to 0.7, ensuring a smooth first and second derivative curves. The inflection point was finally determined by locating the zero point of the second derivative.[64] Mean and SD values of inflection point strain (IPε), apparent (IPσ_A_) and net (IPσ_N_) stress for all the samples categories were calculated.

### Cell Culture

Human mesenchymal stromal cells (hMSCs) were isolated and expanded as previously described[82] from bone marrow aspirates obtained from a single donor after informed consent, based on a corresponding document from the Medical Ethical Committee Toetsingscommissie; METC) of the Medisch Spectrum Twente hospital, Enschede, The Netherlands (study protocol ‘Functioneel weefselherstel met behulp vanuit beenmerg verkregen stamcellen’; K06-002). The cells were cultured in tissue culture flasks at a density of 1500 cells per cm^−2^ and expanded in basic medium, consisting of α-Minimum Essential Medium without nucleotides and with Glutamax (α-MEM, Thermo Fisher Scientific), supplemented with 10% v/v fetal bovine serum (FBS; Sigma-Aldrich), 0.2 × 10^−3^ M L-ascorbic acid 2-phosphate magnesium salt (Sigma-Aldrich), and 100 U/ml penicillin/streptomycin (Thermo Fisher Scientific). Cells were cultured in standard conditions of 37 °C in a humified atmosphere with 5% CO_2_. Medium was replaced every 2-3 days and cells were passaged when reaching 80% confluence.

### Spheroids Formation and Culture

Cell spheroids were formed in U-bottom microwells fabricated by gas-assisted (micro)thermoforming from a polycarbonate (PC) film with a thickness of 50 µm (it4ip), as previously described.[83] Prior to cell culture, the microwells, containing 289 microcavities arranged in a honeycomb-like pattern,[84] were sterilized in a graded series of isopropanol (VWR) (100%, 70%, 50%, 25%, 10% v/v in water) and then washed twice with Dulbecco’s phosphate buffered saline (PBS; Sigma-Aldrich). The sterilized arrays were placed in 24-well plates and secured at the bottom of each well using sterile O-rings (ERIKS). To prevent cell attachment, the microwells were incubated with 350 µl of 1% w/v Pluronic F108 solution (Sigma-Aldrich) overnight at 37 °C. Prior to cell seeding, the Pluronic solution was removed and the microwell arrays containing the membranes were washed three times with sterile PBS. hMSCs at passages 3 or 4 were seeded into the microwell at a density of around 500,000 cells per microwell in basic medium. After 48 hours of incubation at 37 °C and under 5% CO_2_, during which the cells aggregated into spheroids, the culture medium was gently refreshed to remove non-spheroid cells and replaced. Spheroids were cultured for 7 days and medium was replaced every 2-3 days.

### Cell Culture on Electrospun Scaffold

Scaffolds were secured at the bottom of a 24 well-plate using pre-autoclaved O-rings and sterilized together by immersion in 70% ethanol for 1 hour. Afterwards, samples were rinsed three times with PBS. Spheroids were collected from the microwells after gently pipetting up and down several times. The suspension was collected in a 15 ml falcon tube and centrifuged at 300 rpm for 4 min. The supernatant was then aspirated and the spheroids were resuspended in 300 µL of basic medium. Finally, 50 µL of the spheroid suspension was then gently dispensed onto the scaffold. Cells were cultured for 7, 14 and 28 days in basic medium. Medium was refreshed every second day.

### RNA Isolation and Gene Expression Analysis

Total RNA was isolated using the RNeasy Mini Kit (Qiagen) according to the manufacturer’s protocol and RNA purity and concentration were measured using a BioDrop µLITE (BioDrop). Complementary DNA (cDNA) was then synthesized from 250 ng total RNA in a total 20 µL reaction volume, using iScript cDNA synthesis kit (Bio-Rad), following manufacturer’s instructions. Gene expression analysis was performed by real-time quantitative PCR (RT-qPCR) using the iQ SYBR Green Supermix (Bio-Rad) in a CFX96 Real-Time PCR Detection Kit (Bio-Rad) amplifying 20 ng of cDNA in a total volume of 20 µL. Samples were subjected to a thermal cycle of 50 °C for 2 min, 95 °C for 2 min, and then 95 °C for 15 s and 60 °C for 30 s for a total of 40 cycles. Quantification of transcription levels was performed through the ΔΔCt method using glyceraldehyde 3-phosphate dehydrogenase (GAPDH) as housekeeping gene. Results are presented as relative gene expression levels normalized to Day 1 for both EB and EBM scaffolds. The sequences of primers for each gene are listed in **Table 1**.

**Table 1.**
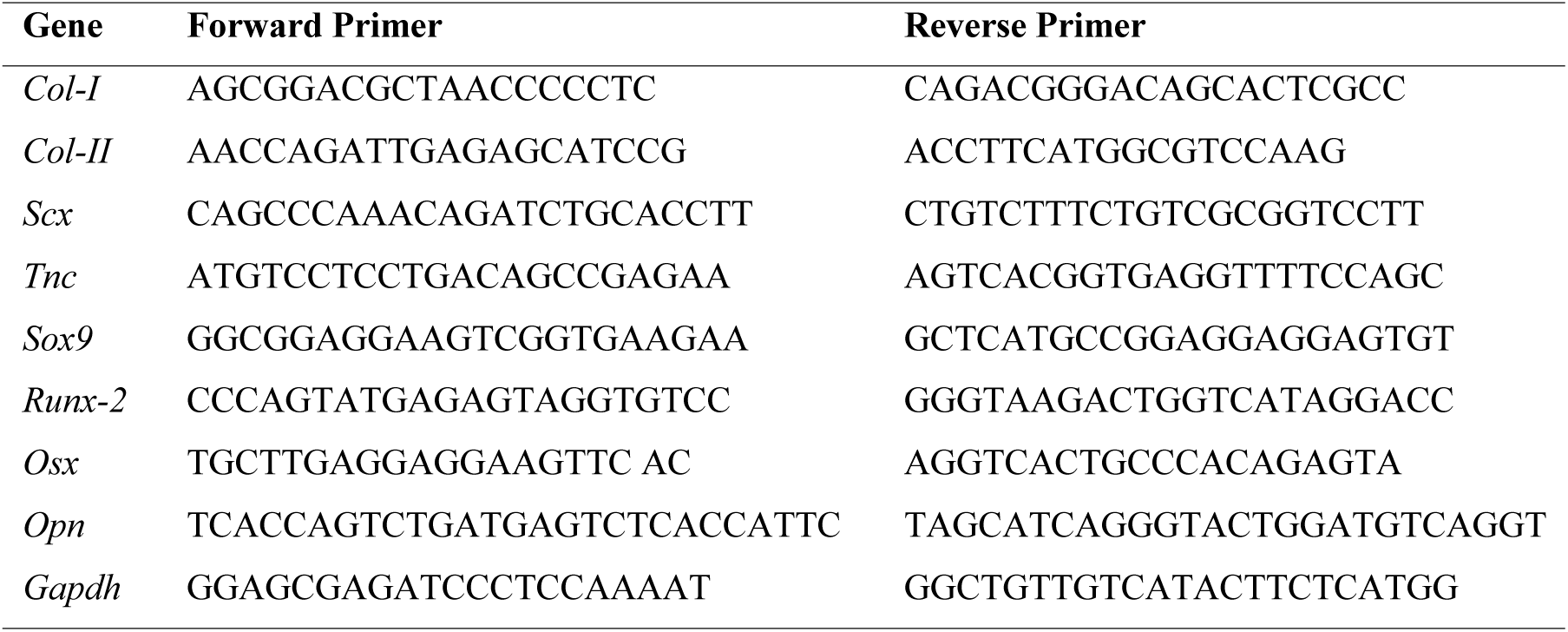
List of primer sequences used for each gene.

### Immunofluorescence Staining and Confocal Fluorescence Microscopy

Cultured scaffolds were washed with PBS and fixed with 4 % (v/v) formaldehyde solution (ACS reagent, Sigma-Aldrich) for 30 min. Fixed samples were treated with 0.1 % (w/v) Sudan Black B dye (in 70 % ethanol; Sigma-Aldrich, the Netherlands) for 45 min and then incubated with 0.1 % Triton™ X-100 (in DPBS; Sigma-Aldrich, the Netherlands) for 15 min at RT. After three washing steps, non-specific binding sites were blocked in CAS-Block (Thermo Fisher Scientific) for 1 h at RT. The blocking solution was then removed and the samples were incubated with the primary antibody rabbit anti-collagen I (dilution 1:100, ab34710, Abcam) or rabbit anti-collagen II (dilution 1:100, ab34712, Abcam) and mouse anti-tenascin C (dilution 1:100, MA5-16086, Thermo Fisher scientific) diluted in CAS-Block solution overnight at 4 °C. After primary antibody incubation, samples were rinsed three times with PBS and incubated for 1 h with secondary antibodies Alexa Fluor 568 anti-rabbit (dilution 1:1000) and Alexa Fluor 488 anti-mouse (dilution 1:1000) in PBS. Mineralized scaffolds were stained with OsteoImage (OsteoImage Mineralization Assay, Lonza), according to the manufacturer’s protocol, to visualize mineral deposits. Finally, F-actin was labeled with phalloidin conjugated with Alexa Fluor 488 or Alexa Fluor 647 (dilution 1:100; Thermo Fisher Scientific) and cell nuclei were counterstained with 4′,6-diamidino-2-phenylindole (DAPI, dilution 1:100; Sigma-Aldrich) for 30 min at RT. The stained samples were mounted under cover slips with Dako Fluorescent Mounting Medium (Agilent) and imaged by confocal laser-scanning fluorescence microscopy using a TCS SP8 STED (stimulated emission depletion) microscope (Leica Microsystems). The images were acquired as *z*-stacks with slices every 10 µm at a 25× magnification using a water immersion objective.

The quantification (n=3 images per each region; n=3 images for EB and EBM scaffold; n=3 images per each time point) of ECM production was performed using CellProfiler (https://cellprofiler.org/<u>; version 4.2.0</u>) by customized pipelines.[85] Briefly, in each pipeline, the nucleus morphology was identified as a primary object using the Otsu adaptive thresholding method on the DAPI channel, and cell morphology was determined using the propagation algorithm in combination with the Otsu adaptive thresholding on the phalloidin channel. Cells touching the edges of the images were excluded from the dataset. After background correction, COL-I, COL-II, TNC and OsteoImage staining were quantified as the pixel area stained by the input marker relative the total segmented cell area.

The orientation of actin filaments of hMSCs was carried out by using the Directionality plugin of ImageJ (Local Gradient Orientation method) on confocal images, per each time point of culture and per each region of EB and EBM scaffolds (i.e. TB, J, FB and FBM) of the actin staining, along the axe of scaffolds. In brief each image was at first converted in 8 bit and the gray levels were homogenized to ensure the same grey levels for all the images. Then the Directionality plugin was applied and the results were shown as mean ± SD assuming as axial orientation 0° and 90° as the transversal one.

#### ELISA

COL-1, COL-II, TNC and TNMD production by hMSCs production was analyzed after 7, 14 and 28 days of culture on both scaffolds using ELISA. The test was performed on 3 samples per condition. Briefly, the culture medium was centrifuge at 2000x g at 4 °C for 20 min to remove cell debris and stored at −80 °C until all groups were collected. Total protein concentration was determined with BCA assay (Thermo Fisher). COL-I was assessed using Human Collagen Type I ELISA Kit (abcam, ab28250). COL-II was assessed using Human COL2alpha1 ELISA Kit (Amsbio, HO777). TNC was assessed using Human Tenascin-C ELISA Kit (abcam, 213831) and TNMD using Human Tenomodulin ELISA Kit (MBS9713860, BioSource).

### Calcium Assay

Calcium release was quantified using the Calcium Colorimetric Assay Kit (Sigma Aldrich), following manufacturer instruction.

### Synchrotron X-ray 3D imaging

Defined the most promising candidate between the two categories of scaffolds (i.e. EB and EBM), synchrotron experiments were performed only on the non-mineralized ones (i.e. EB). To capture the 3D multi-scale morphology and the full-field local mechanics of the scaffolds, the complementarity of two ESRF beamlines was employed: ID16B (SnCT at 100 and 300 nm of voxel size) and ID19 (SµCT at 650 nm of voxel size). The nano-scale imaging instrument available at the ID16B beamline was used with a conic and quasi-monochromatic beam (ΔE/E = 10^-2^) at an energy of 29.2 keV and a flux of 8 × 10^11^ ph s^-1^.[86] The technique of holo-tomography was applied by acquiring phase contrast 3D imaging for four slightly different propagation distances.[87] For each tomographic scan along a 360° rotation, 1839 projections were recorded with an exposure time of 10ms each on a PCO edge 5.5 sCMOS camera (2560 x 2160 pixels) equipped with a 17 µm LSO scintillator. The pixels of the detector were binned (1280 × 1080 pixels) to have four times the flux per pixel and thus reducing the total acquisition time down to ∼19 sec per scan. Reconstruction of the ID16B data was achieved with the open-source software PyNX through a two-step approach.[88] Firstly, an iterative phase retrieval calculation was applied using as an initial guess a Paganin-like approach with a delta-to-beta ratio δ/β = 150. Subsequently, a filtered back-projection reconstruction was performed using the ESRF software Nabu (https://gitlab.esrf.fr/tomotools/nabu), yielding volumes of 128 × 128 × 108 µm^3^ with an isotropic voxel size of 100 nm for the morphological investigation, and volumes of 384 × 384 × 324 µm^3^ with an isotropic voxel size of 300 nm for the *in situ* mechanical tests.

A parallel pink beam provided by the first harmonic of a short-period undulator (λ = 13 mm) with an energy of 25.9 keV (2.6% B.W.) was used at the ID19 beamline[89] providing an estimated flux of 13×10^12^ ph/0.1% B.W/s. The sample was positioned 145 m away from the source, and the beam was conditioned by a series of two in-vacuum slits and permanent filters along the flight tube (0.8 mm diamond, 2 mm Be) mostly for heat load moderation. The indirect detector consisted of high-resolution optical assembly[90] with 8 µm GaGG:Ce single crystal scintillator, 10x objective lens (Olympus, UPlan, NA=0.4), folding mirror and a PCO.edge 5.5 sCMOS camera positioned 90 degrees with respect to X-ray beam. A consecutive vertical stack of 3 tomographic scans, covering the FB, J and TB sides of an EB sample was acquired with an overlap of 270 µm per scan. For each scan, 4000 projections were acquired along a 360° rotation with an exposure time of 40 ms and total acquisition time of 3 min per one tomogram. The reconstruction was performed again with the ESRF software Nabu using the single distance phase retrieval with a delta-to-beta ratio δ/β = 200. The reconstructed volumes were stitched after applying spectral ring filtering correction[91], producing a final volume of 2560×2560×5600 voxels, with an isotropic voxel size of 650 nm.

The 3D-renderings of the scans of FB, TB, J and EB were carried out by using the Dragonfly software (Comet Technologies Canada Inc., Montreal, CAN). Moreover, the orientations of nanofibers inside the different regions of scaffolds were measured by using the Directionality plugin of ImageJ on internal crops (590 slices each of 1094 × 1072 pixels) of scans at 100 nm of voxel size for all the different regions of EB (i.e. FB, TB and J) with the same parameters previously defined. At each bin of the directionality histogram was attributed the mean ± SD among the whole image stack for the dedicated range of angles.

### Synchrotron X-ray in situ mechanical tests

*In situ* tensile tests were carried out at the ESRF using a dedicated electromechanical testing machine. The load frame (called Nanox2) for in situ X-ray synchrotron experiments was developed at the Centre des Matériaux (Paris, France) within the course of the Long Term Proposal MA4925 at ESRF. The frame uses a Faulhaber brushless DC motor to move a crosshead with a travel range of 6 mm and a tunable displacement speed. The cable system regroups in one connector the motor control and the load cell input and output lines and has been designed to be compatible with slip rings mounted on various tomography stations so that *in situ* loading experiments can be carried out seamlessly. The stress rig was modified to comply with the tomography stations of both ID16B and ID19 beamline using a modified grip system, which sets the sample at 53 mm of the base plate. A magnetic connector was installed (see Fig. S6) so that the motor was powered off between loading steps and the cable disconnected to acquire tomographic scans. The lower jaw was mobile while the upper jaw transmitted the load to the frame via a glassy carbon tube (transparent for X-ray imaging). To clamp the scaffolds, dedicated 3D printed grips were developed. To prevent stress concentration, bundles were glued between pieces of carboard which were held in compression via a set screw in the grip. The gauge length of the samples was about 4.5 mm.

The testing protocol consisted in a stepwise progressive axial loading. Firstly, a reference scan was acquired at the pristine state (unloaded sample). An axial displacement was then applied, with a loading rate of 2 µm s^-1^ (strain rate = 0.045 % s^-1^). To stabilize the sample relaxation and micro-movements, after each loading step, a relaxation period of 8 min was applied. After this period, a tomographic acquisition was performed. This sequence was repeated until the last scan, which was taken right after the failure of the specimen. Note that the number of scans and the displacement steps were different for each sample, justified by the fact that the loading was interrupted at different points attempting to catch significant changes in the macroscopic response.

Three sets of samples (TB, FB and EB) were tested *in situ* at ID16B to capture the nano-scale mechanics of the three distinct regions (TB, FB, and J). For the EB sample, similarly with the *ex situ* tests, the junction was placed in the middle of the gauge length. Taking advantage of the larger field of view at ID19 (sacrificing spatial resolution) an EB sample was tested *in situ*, directly imaging the three regions of the entire scaffold. The combination of the imaging instruments in both beamlines allowed for a multiscale full-field mechanical investigation of EB scaffolds and their corresponding regions, from the structural level at 650 nm down to the nanofiber level at 300 nm.

A series of *ex situ* tests was also performed to determine the loading steps and decouple the mechanical response from the beam impact.

### Digital Volume Correlation (DVC)

The full-field 3D deformation fields were measured with the open-source software SPAM.[92] A total DVC analysis was carried out by mapping the first scan (i.e., undeformed sample) with each of the remaining scans, as opposed to an incremental analysis which maps two consecutive load steps. As a first step of the DVC analysis, an alignment of each pair of images was done through a rigid-body registration (capturing overall displacements and rotations). A local DVC approach followed, where a set of nodes was defined in the reference image (i.e. undeformed sample) around which independent cubic sub-volumes (i.e. correlation windows) were extracted and sought in the deformed image based on a gradient-based iterative procedure. The size of the correlation windows and the number of measurement nodes depend on the texture of the imaged samples and define the spatial resolution of the measured fields. To overcome the problem of the large and heterogeneous deformation fields particularly encountered in a total DVC analysis, as well as, to achieve a high spatially resolved deformation field, a pyramidal coarse-fine approach was employed. It consisted of running a series of computations with gradually decreasing the correlation windows size and the node spacing. For all pyramidal levels, an overlap of 50% was set between neighboring correlation windows. For the *in situ* tests at ID16B, the pyramidal levels of the correlation window sizes were [160, 80, 48, 24]vx^3^ or [48, 24, 7.2, 4.2]µm^3^, while for the ones at ID19 they were [200, 120, 80, 50]vx^3^ or [130, 78, 52, 32.5]µm^3^.

Strains were obtained from the displacement vectors measured at the nodes of the finest pyramidal level, by computing the transformation gradient tensor F (local derivative of displacements) on Q8 shape functions, linking 2 × 2 × 2 measurement nodes. The displacement field was firstly smoothed by applying a 3D median filter of a 1 voxel radius. A polar decomposition of F yielded the right stretch tensor U and the rotation tensor R for each Q8 element. The finite large-strain framework was used to calculate the following strain invariants:

1. the volumetric strain (ε_V_):

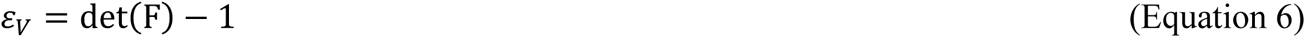

ii) the principal maximum (ε_p1_) and minimum (ε_p3_) strains based on the diagonalization of the right Cauchy-Green strain tensor:

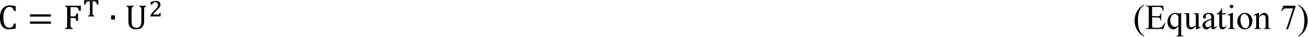

iii) the deviatoric strain (ε_D_), based on a multiplicative decomposition of the stretch tensor U into an isotropic and deviatoric part:

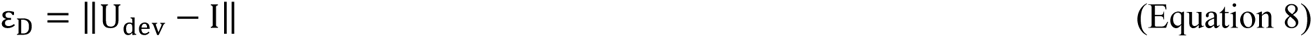

Where I is the identity matrix with:

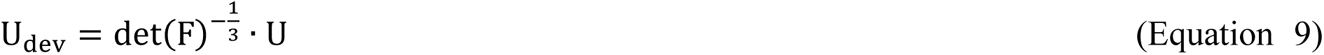

### DVC uncertainty and beam impact

An investigation of the radiation induced damage, coupled with the DVC measurements uncertainty was performed for both ID16B and ID19 beamlines. Based on the fact that the region of the EB scaffold that exhibited the higher strain magnitude was the J one, the coupled investigation was focused on this region. The tested samples were firstly loaded at a small strain level (i.e. 0.5 %) avoiding to exhibit the initial waviness of the nanofibers, induced by the removal of bundles from the drum collector and the subsequent crosslinking. At the ID16B beamline, four successive holo-tomography scans were acquired at a J type sample. During the SnCT *in situ* tensile tests, a single holo-tomography scan (i.e. one loading step) was performed between the intact (i.e. zero-strain) and the post-failure state of the material. This means that the accumulated dose on this radiation test sample is triple the one induced during the *in situ* tensile tests. Since the levels of the measured DVC strains during the SµCT *in situ* test were lower compared to the SnCT one, two successive scans were performed at an EB sample at ID19 focusing on the J region. Through the ε_V_ and ε_p3_ strain components, although an overall progressive contraction of the material was measured (caused by its progressive drying and damaging of the polymeric chains), no local strain localization or macroscopic failure of the sample was observed for any beamline. Moreover, the accumulative strain (by the end of the fourth successive scan for ID16B and the second scan for ID19) reached values of one order of magnitude lower compared to the ones measured during the *in situ* tests, serving as a direct quantification of the DVC measurement of uncertainties for both beamlines. On top of providing a measure for the DVC uncertainty, the successive scans showed that there was a contribution of the X-rays to the observed mechanical behavior (causing an overall homogeneous contraction of the material), but this contribution was not sufficient by itself to cause the macroscopic failure of the sample.

### Statistical Analysis

The significance of differences of the apparent and net tensile mechanical properties of the different scaffolds’ categories, at the same strain-rate (i.e. FB, FBM, EB, EBM and TB; n=5 for each), was assessed with a one-way ANOVA followed by a Tukey post hoc (ns p>0.05, *p≤0.05, **p≤0.01, ***p≤0.001, ****p≤0.0001) (Figure 3 and Table S5 and S6). The significance of differences of the apparent and net tensile mechanical properties of the same scaffolds, at the different strain-rates (n=5 for each), was assessed with a one-way ANOVA followed by a Tukey post hoc (ns p>0.05, *p≤0.05, **p≤0.01, ***p≤0.001, ****p≤0.0001) (Table S7 and S8).

The significance of differences of the ECM quantifications and OsteoImage of EB and EBM between the different time points and regions (7, 14 and 28 days; TB, J, FB), assessed with a one-way ANOVA followed by a Tukey post hoc (ns p>0.05, *p≤0.05, **p≤0.01, ***p≤0.001, ****p≤0.0001) (Figure 5G, Table S11).

The significance of differences of each gene expressed by hMSCs on EB and EBM at the different time points of culture (7, 14 and 28 days) assessed with a one-way ANOVA followed by a Tukey post hoc (ns p>0.05, *p≤0.05, **p≤0.01, ***p≤0.001, ****p≤0.0001) (Figure 6A and Table S13). The significance of differences between the same gene expressed by hMSCs on EB and EBM at the different time points of culture (7, 14 and 28 days) assessed with an unpaired parametric t-test with Welch’s correction (ns p>0.05, *p≤0.05, **p≤0.01, ***p≤0.001, ****p≤0.0001) (Figure 6A and Table S14).

The significance of differences of each ECM protein expressed by hMSCs on EB and EBM at the different time points of culture (7, 14 and 28 days), detected via Elisa test, were assessed with a one-way ANOVA followed by a Tukey post hoc (ns p>0.05, *p≤0.05, **p≤0.01, ***p≤0.001, ****p≤0.0001) (Figure 6B and Table S16).

The significance of differences between EB and EBM of each ECM protein expressed by hMSCs at the different time points of culture (7, 14 and 28 days), detected via Elisa test, were assessed with an unpaired parametric t-test with Welch’s correction (ns p>0.05, *p≤0.05, **p≤0.01, ***p≤0.001, ****p≤0.0001) (Figure 6B and Table S17).

The significance of differences between the calcium assays of control medium, EB cellularized and cellularized with and without cells at different time points (7, 14 and 28 days) assessed with a one-way ANOVA followed by a Tukey post hoc (ns p>0.05, *p≤0.05, **p≤0.01, ***p≤0.001, ****p≤0.0001) (Figure 6C and Table S19).

## Supporting Information

Supporting Information at the end of the manuscript.

## Acknowledgments

Francesca Giacomini and Olga Stamati contributed equally to this work.

The European Commission and the Horizon Europe Marie Postdoctoral Fellowship project 3NTHESES (n. 101061826) are greatly acknowledged for funding the study. Kensey Nash Corporation d/b/a DSM Biomedical and Corbion are acknowledged for furnishing the COL-I and PLLA respectively. The European Synchrotron Radiation Facility (ESRF) is greatly acknowledged for furnishing the in-house research beamtime IHLS3489 at ID16B and IHLS3505 at ID19. The EPSRC (EP/W003333/1) project is acknowledged for have partially funded Bratislav Lukic fellowship. The financial support of the Gravitation Program of the Netherlands Organization for Scientific Research (NWO) (project ‘Materials-Driven Regeneration’; grant no. 024.003.013) is greatly acknowledged. Alessandra Di Lorenzo is acknowledged for the help in samples preparation and morphological/mechanical characterization.

## Conflict of Interest

The authors declare no conflict of interest.

## Data Availability Statement

Data will be made available on request.

## ToC

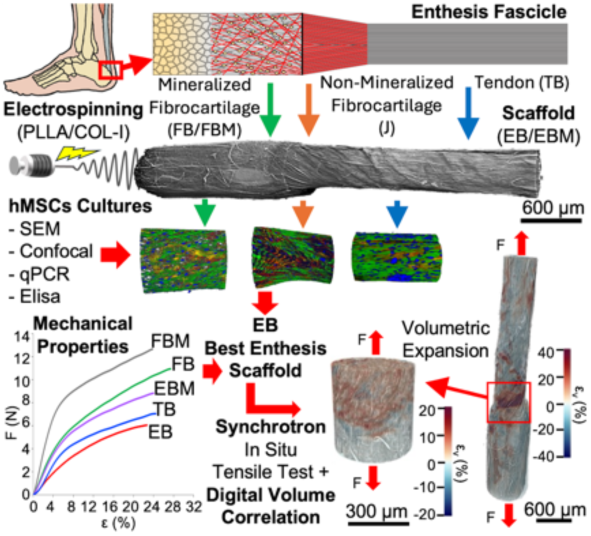

Enthesis injuries are a worldwide healthcare problem. Biomimetic electrospun enthesis fascicle-inspired scaffolds, with and without nano-mineralization are developed. hMSCs express the most balanced enthesis markers on the non-mineralized scaffolds. Multiscale synchrotron *in situ* tensile tests with digital volume correlation reveal the full-field strain distribution of nanofibers at the nanoscale, detecting a volumetric expansion at the junction region like natural enthesis.

## Supporting Information

**Figure S1.**
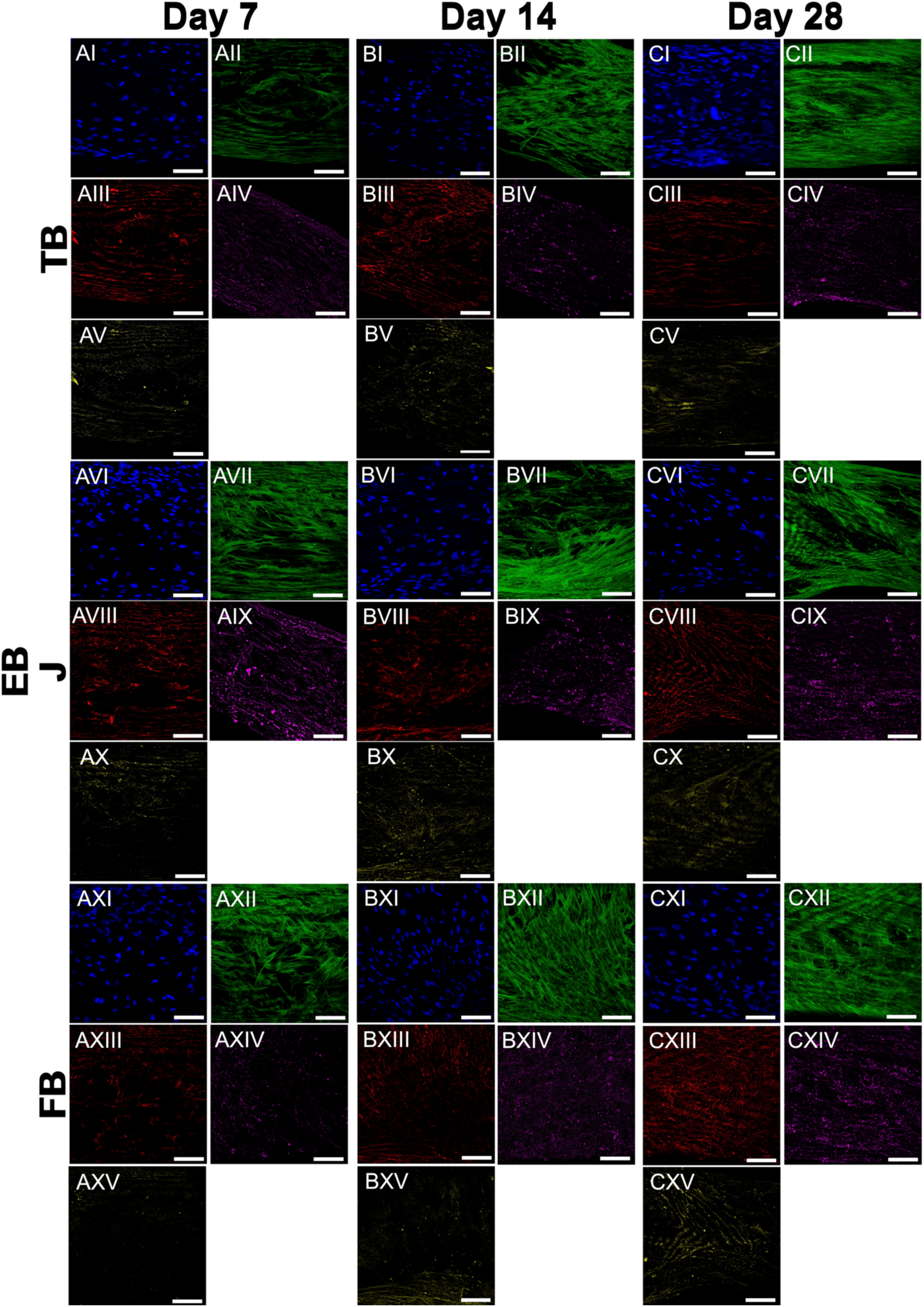
Confocal images of the separate channels of EB at the different time points. A) 7 days, B) 14 days and C) 28 days of culture: I, VI, IX) hMSCs nuclei (blue); II, VII, XII) hMSCs actin (green); III, VIII, XIII) COL-I (red); IV, IX, XIV) COL-II (purple); V, X, XV) TNC (yellow) (scale bar = 100 µm).

**Figure S2.**
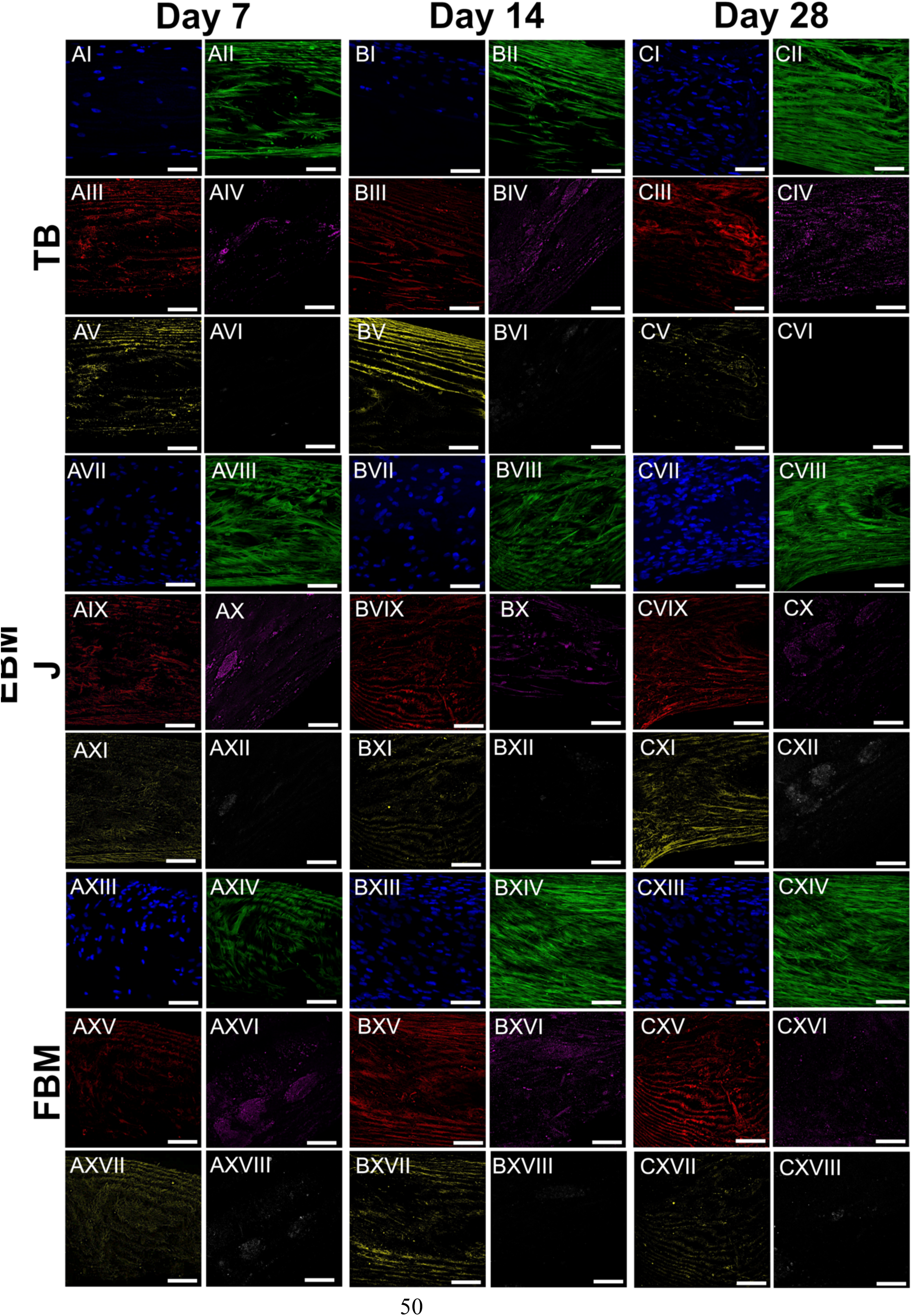
Confocal images of the separate channels of EBM at the different time points. A) 7 days, B) 14 days and C) 28 days of culture: I, VII, XII) hMSCs nuclei (blue); II, VIII, XIV) hMSCs actin (green); III, IX, XV) COL-I (red); IV, X, XVI) COL-II (purple); V, XI, XVII) TNC (yellow); VI, XII, XVIII) hydroxyapatite (gray) (scale bar = 100 µm).

**Figure S3.**
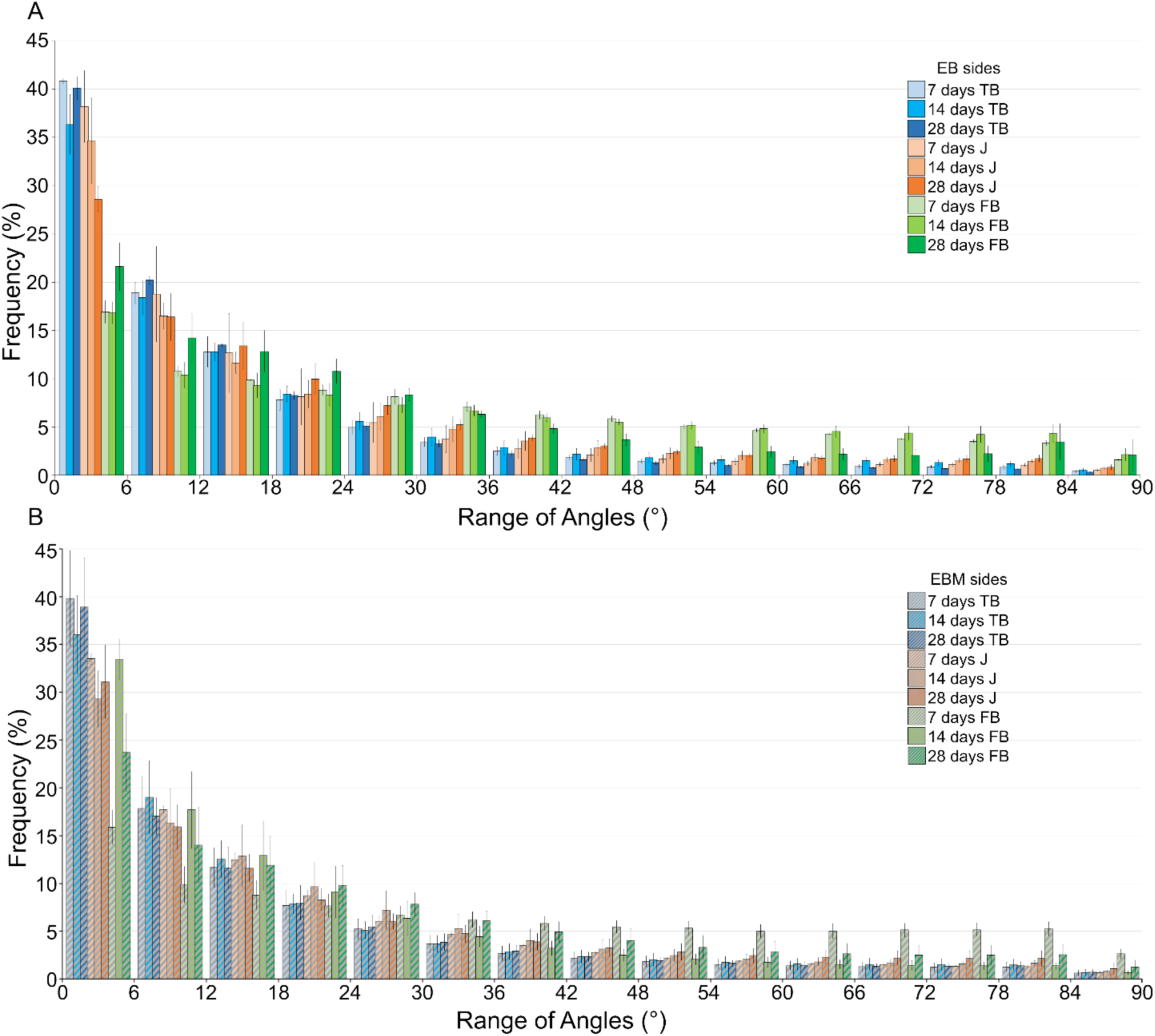
Evolution of actin filaments orientation of hMSCs grown on the different regions of EB and EBM during the time points of culture. In the histograms 0° represents the axial orientation while 90° the transversal one with respect to the axis of EB or EBM. A) Orientation of actin filaments in the different regions of EB scaffolds; B) orientation of actin in the different regions of EBM scaffolds.

**Figure S4.**
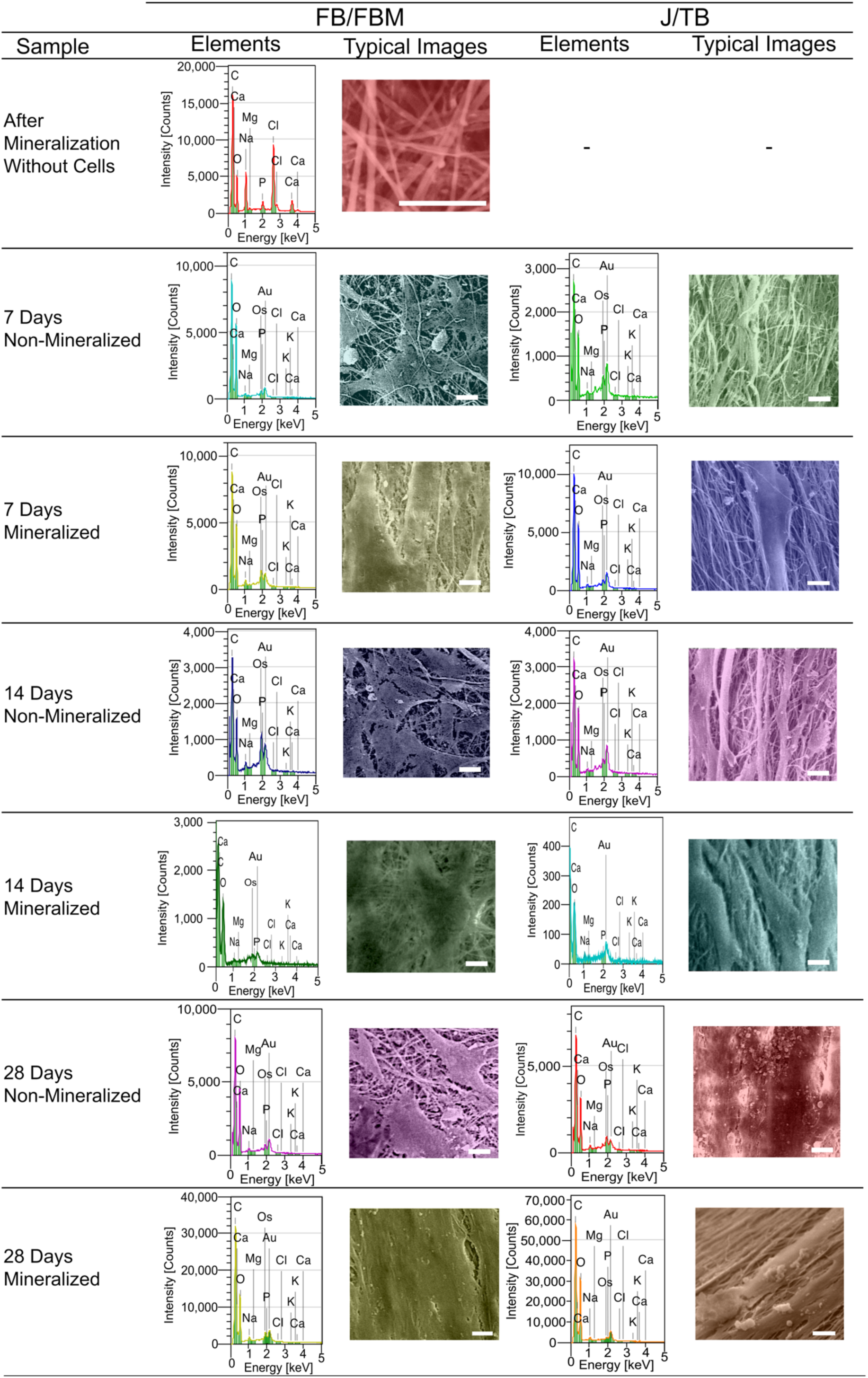
EDS investigations on the mineralized and non-mineralized bundles at the different time points of the hMSCs cultures (scalebar = 10 µm).

**Figure S5.**
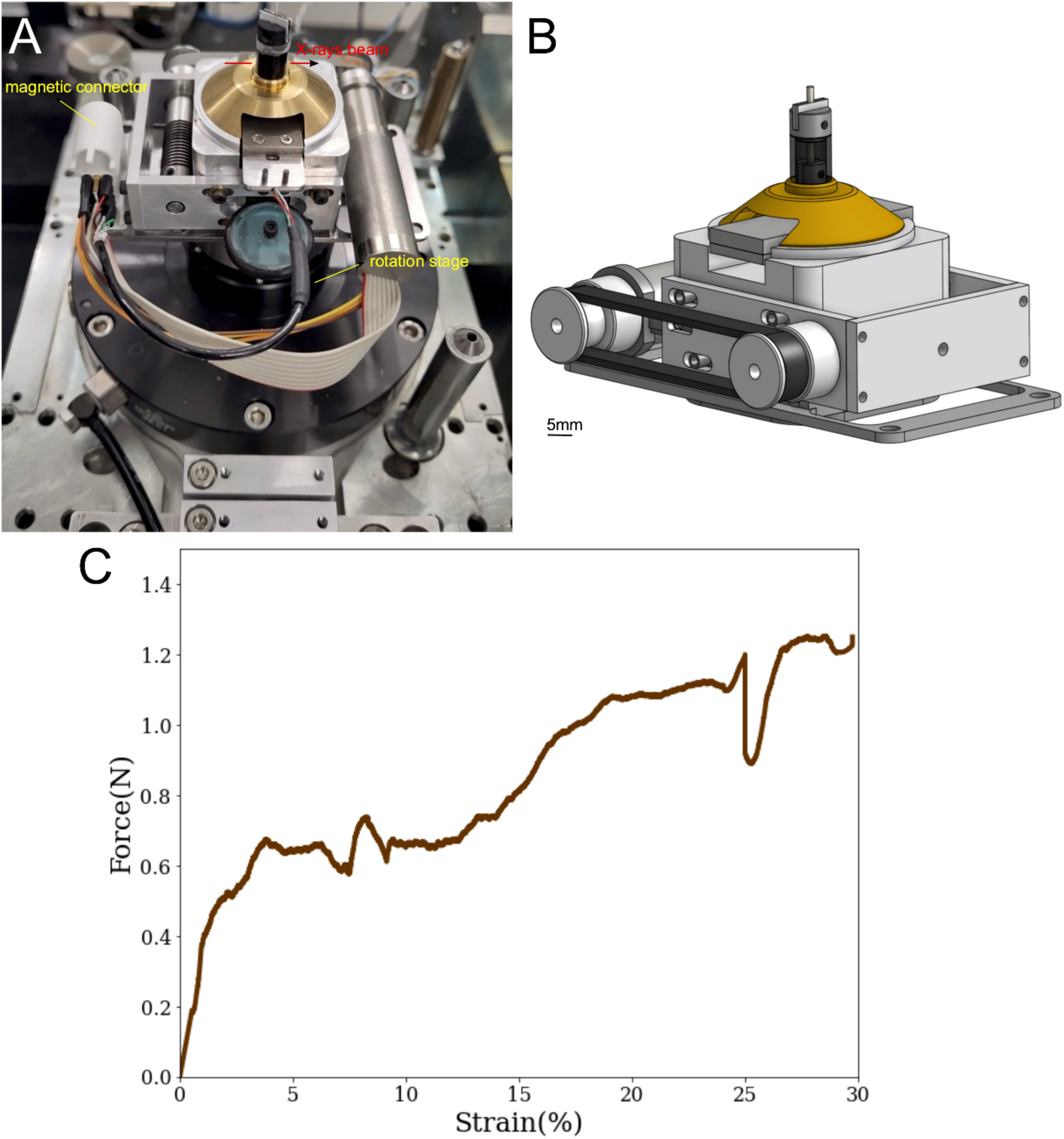
*In situ* tensile testing machine. A) machine mounted on the rotation stage at ID16B beamline; B) 3D rendering of the *in situ* tester; C) typical *ex situ* curve of EB.

**Figure S6.**
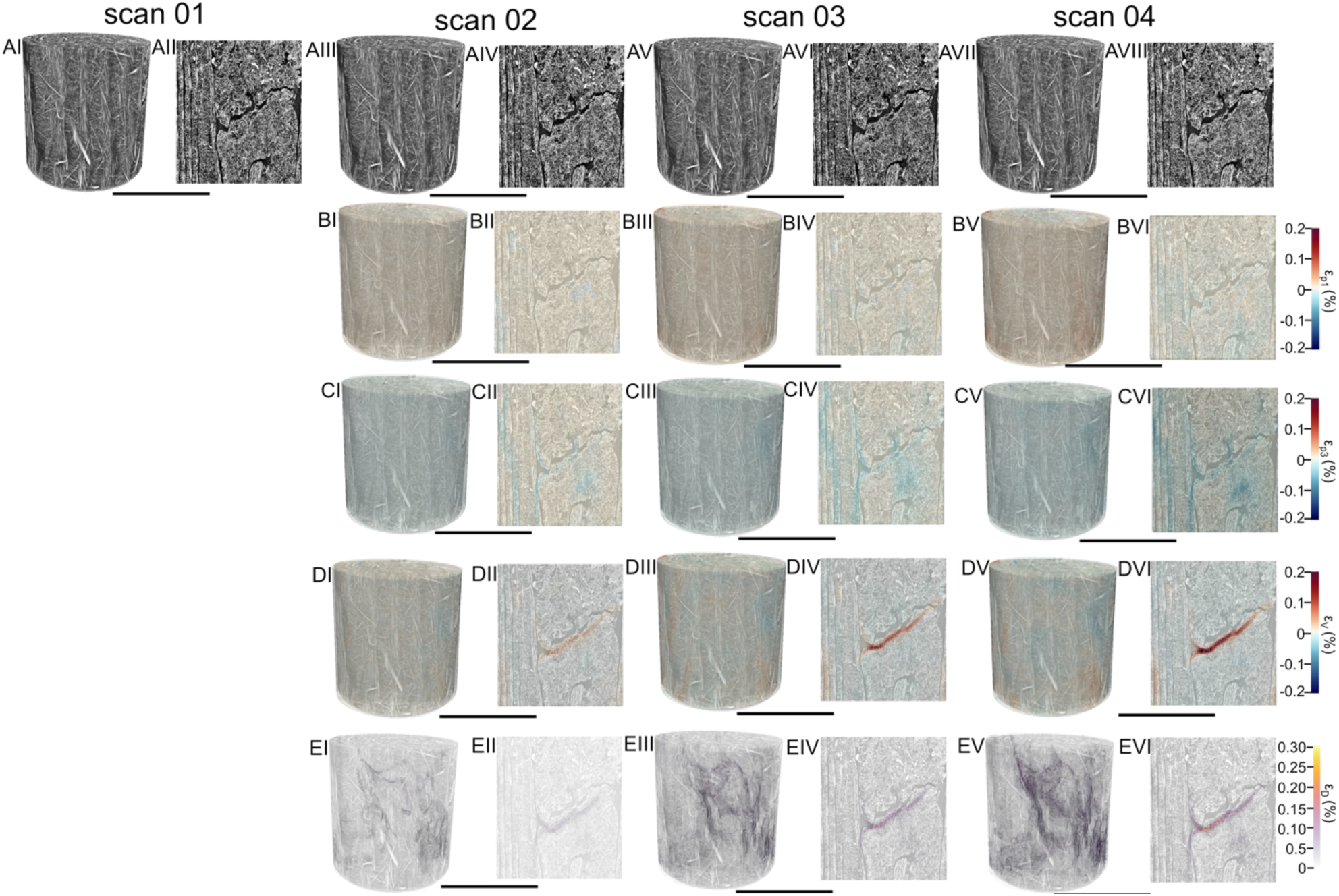
Evaluation of DVC uncertainty and radiation dose effect of SnCT at ID16B. Repeated scans have been performed for the sample without loading and DVC calculation have been performed for the different reconstructed volumes. I, II) Scan 1, III, IV) scan 2, V, VI) scan 3, V, VI) scan 4. A) Greyscale volume rendering and internal slices; B) ε_p1_; C) ε_p3_; D) ε_V_; E) ε_D_ (voxel size = 300 nm, scale bar = 300 µm).

**Figure S7.**
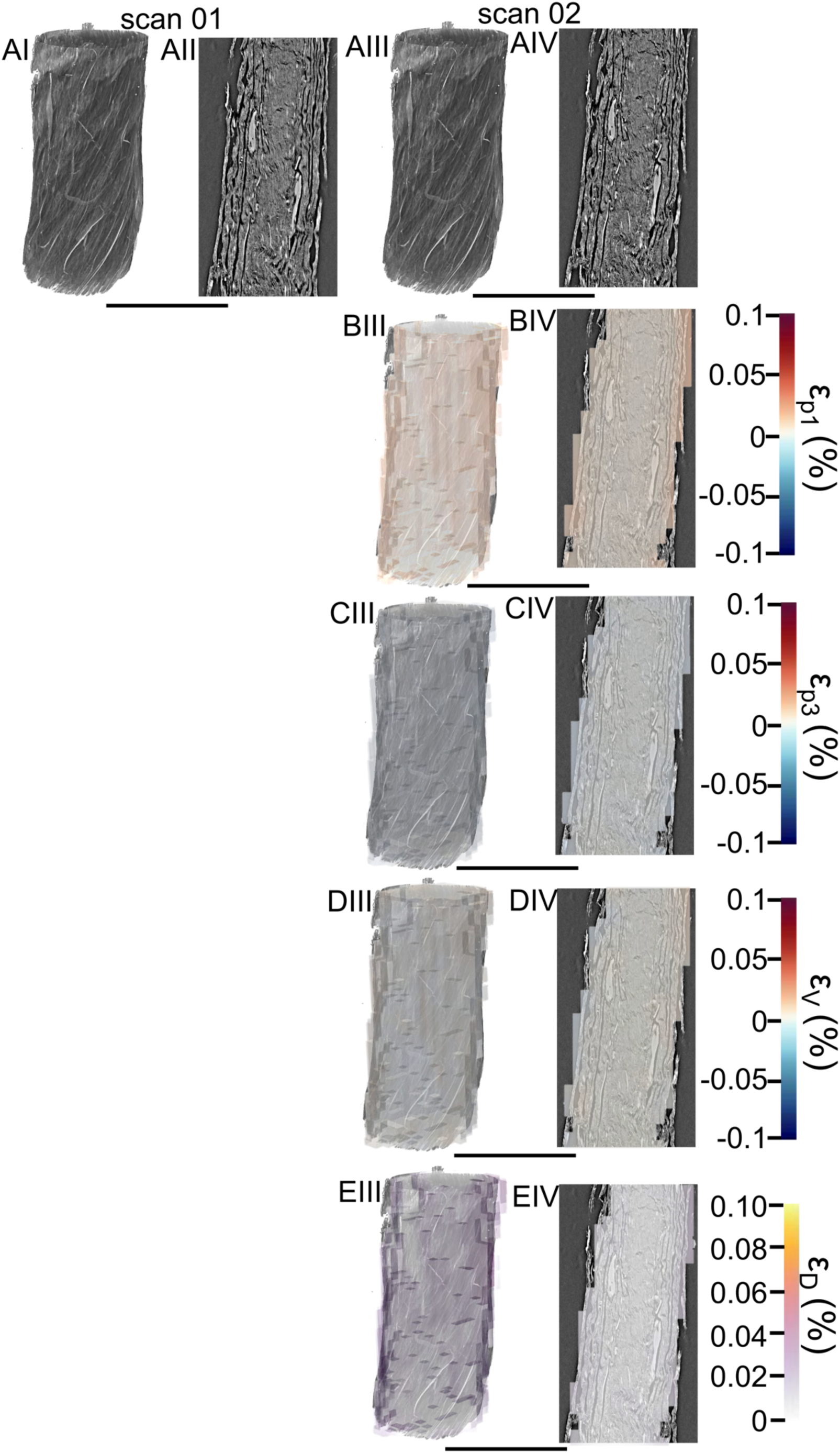
Evaluation of DVC uncertainty and radiation dose effect of SµCT at ID19. Two repeated scans have been performed for the sample without loading and DVC calculation have been performed between the two reconstructed volumes. I, II) Scan 1, III, IV) scan 2. A) Greyscale volume rendering and internal slices; B) ε_p1_; C) ε_p3_; D) ε_V_; E) ε_D_ (voxel size = 650 nm, scale bar = 600 µm).

**Table S1.** Mean morphological parameters of scaffolds for mechanical tests.

| Sample | l<br>[mm] | l<sub>sk</sub><br>[mm] | l<sub>0</sub><br>[mm] | sk<br>[%] | l<sub>0</sub>/l<sub>sk</sub> | D<br>[$\mu\text{m}$] | d<br>[$\mu\text{m}$] | A<br>[mm<sup>2</sup>] | $\rho_M$<br>[mg mm<sup>-3</sup>] | $\rho_{BM}$<br>[mg mm<sup>-3</sup>] | v<br>[-] | P<br>[%] |
| --- | --- | --- | --- | --- | --- | --- | --- | --- | --- | --- | --- | --- |
| EB | 40 | 32±0.5 | 20 | 19±1 | 0.6 | 635±10 | - | 0.32±0.01 | - | - | 0.39±0.04 | 61±4 |
| FB | 40 | 34±0.3 | 20 | 16±1 | 0.6 | 733±20 | 0.39±0.19 | 0.42±0.03 | - | - | 0.44±0.06 | 56±6 |
| EBM | 40 | 32±1.1 | 20 | 19±3 | 0.6 | 612±10 | - | 0.29±0.01 | 0.03±0.008 | 1.3±0.01 | 0.45±0.05 | 55±5 |
| FBM | 40 | 34±0.6 | 20 | 16±2 | 0.6 | 733±30 | 0.41±0.16 | 0.42±0.02 | 0.02±0.019 | 1.29±0.02 | 0.47±0.04 | 53±4 |
| TB | 40 | 34±0.2 | 20 | 14±1 | 0.6 | 456±30 | 0.30±0.16 | 0.16±0.03 | - | - | 0.47±0.004 | 51±3 |

**Table S2.** Mean morphological parameters of scaffolds for mechanical tests.

| Sample | w<br>[mg] | w<sub>FBM</sub><br>[mg] | w<sub>M</sub><br>[mg] | w<sub>0</sub><br>[mg] |
| --- | --- | --- | --- | --- |
| EB | 5.1±0.3 | - | - | 3.2±0.2 |
| FB | 7.9±0.5 | - | - | 4.7±0.3 |
| EBM | 5.3±0.5 | 5.5±0.5 | 0.2±0.05 | 3.4±0.2 |
| FBM | 8.2±0.3 | 8.6±0.5 | 0.4±0.29 | 5.1±0.3 |
| TB | 3.4±0.4 | - | - | 2.0±0.2 |

**Table S3.**
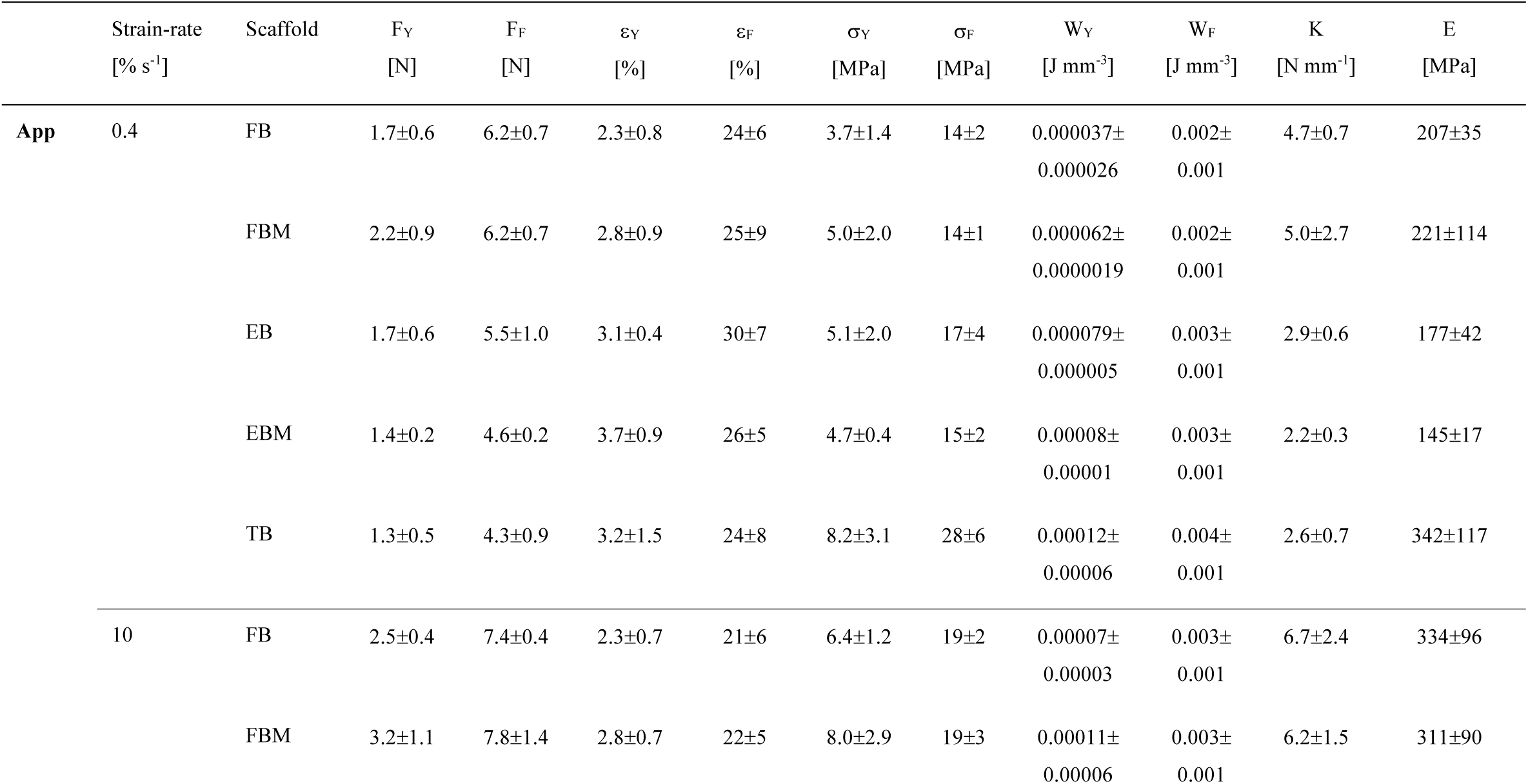
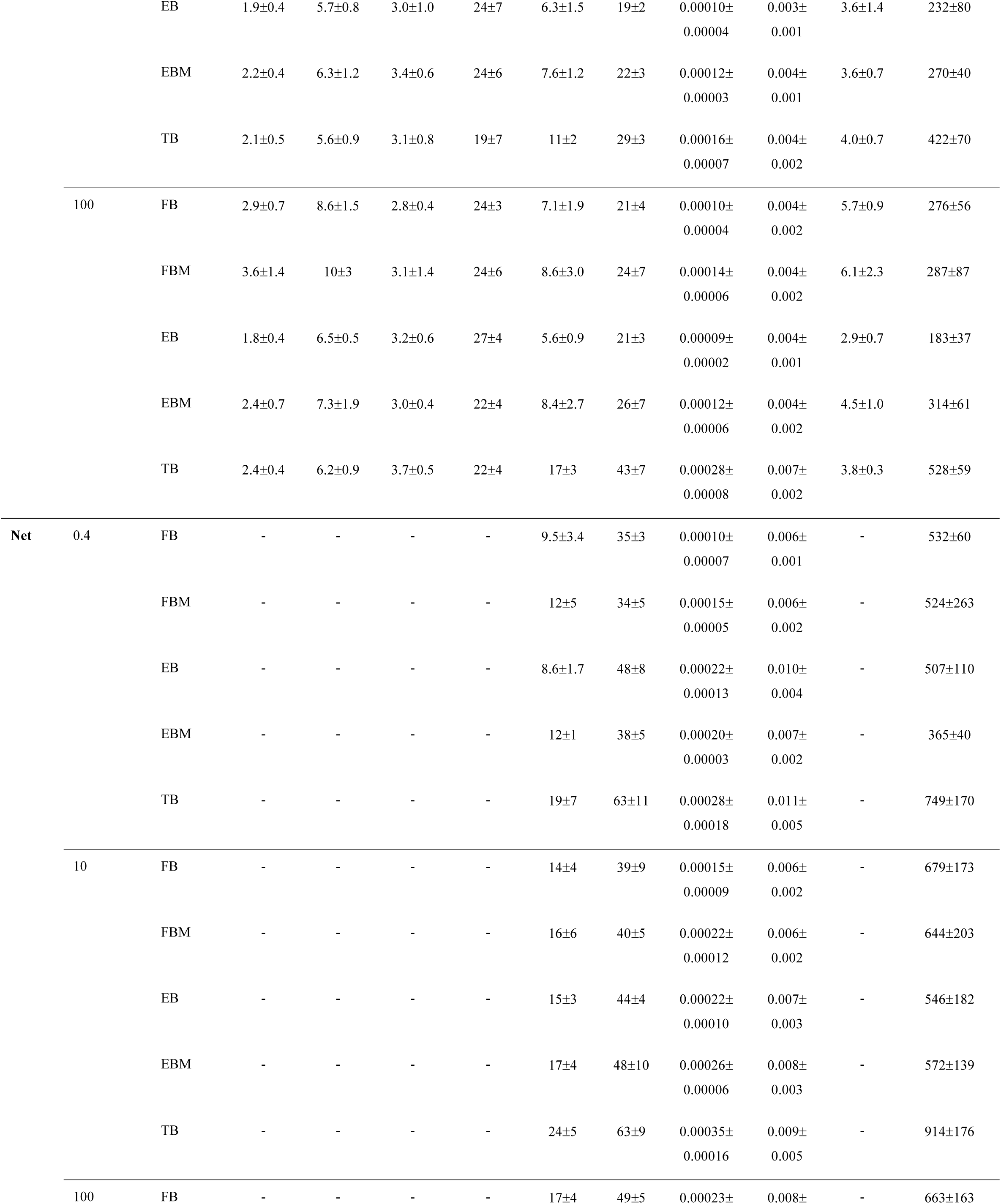
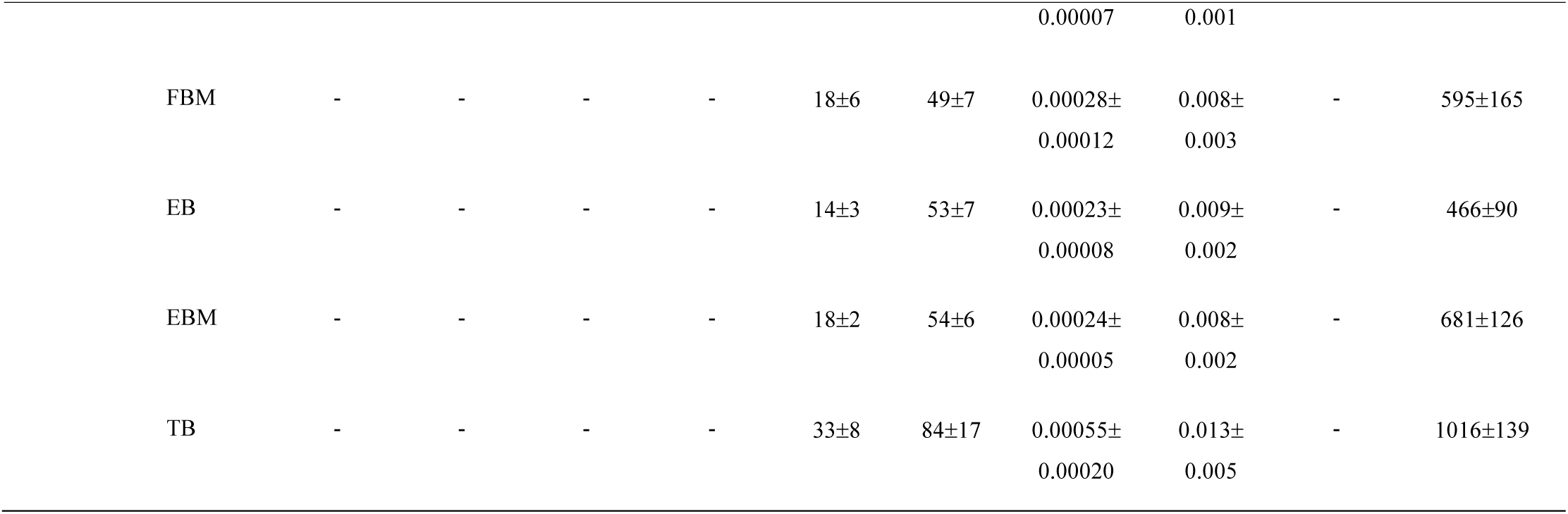
Apparent and net mechanical properties of scaffolds at the different strain-rates investigated.

**Table S4.** Apparent and net inflection point properties.

| Strain-rate<br>[% s<sup>-1</sup>] | Scaffold | IFF<br>[N] | IFε<br>[%] | IF$\sigma_{App.}$<br>[MPa] | IF$\sigma_{Net}$<br>[MPa] |
| --- | --- | --- | --- | --- | --- |
| 0.4 | FB | 1.4±0.5 | 1.9±0.9 | 2.8±1.2 | 7.3±3.2 |
|  | FBM | 1.4±0.6 | 1.7±0.6 | 2.8±1.4 | 6.7±3.5 |
|  | EB | 0.8±0.3 | 1.4±0.4 | 2.1±0.9 | 6.5±2.1 |
|  | EBM | 0.9±0.4 | 2.2±1.1 | 2.6±1.0 | 6.6±3.0 |
|  | TB | 1.0±0.3 | 2.5±1.3 | 5.8±2.2 | 13±6 |
| 10 | FB | 1.4±0.2 | 1.1±0.4 | 3.0±0.7 | 6.2±2.2 |
|  | FBM | 1.7±0.8 | 1.4±0.5 | 3.6±2.0 | 7.4±4.5 |
|  | EB | 1.0±0.2 | 1.4±0.3 | 2.7±0.6 | 6.4±1.3 |
|  | EBM | 1.4±0.2 | 2.2±0.7 | 4.6±0.7 | 11±4 |
|  | TB | 1.5±0.5 | 2.1±1.0 | 7.2±3.0 | 16±6 |
| 100 | FB | 3.0±0.3 | 2.1±0.4 | 5.3±0.6 | 13±2 |
|  | FBM | 3.2±1.0 | 1.9±0.2 | 5.5±1.9 | 11±4 |
|  | EB | 1.5±0.3 | 1.9±0.5 | 3.4±0.4 | 8.7±0.7 |
|  | EBM | 2.3±0.4 | 2.1±0.2 | 5.8±1.1 | 13±3.0 |
|  | TB | 2.1±0.4 | 2.5±0.4 | 11±1 | 21±5 |

**Table S5.**
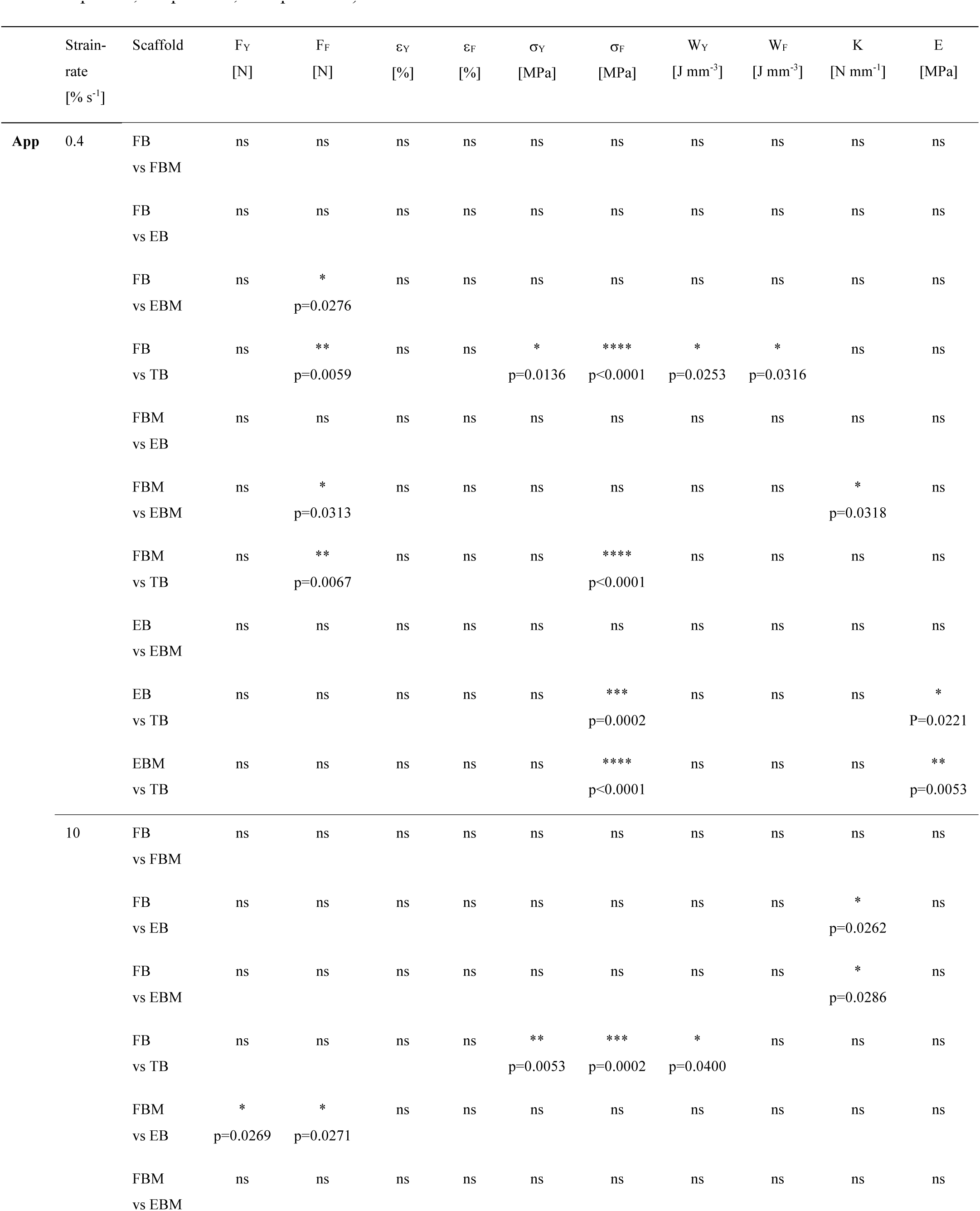
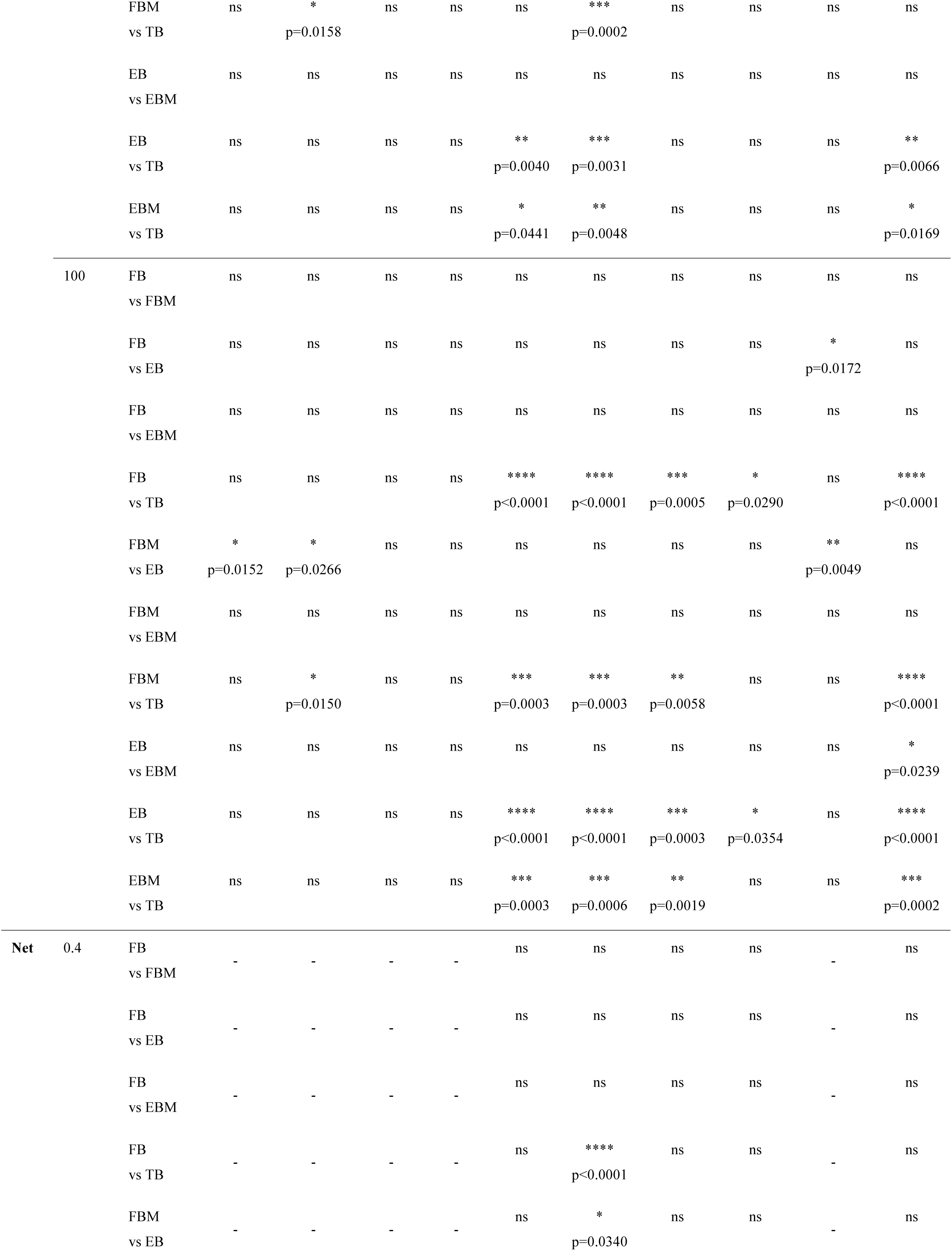
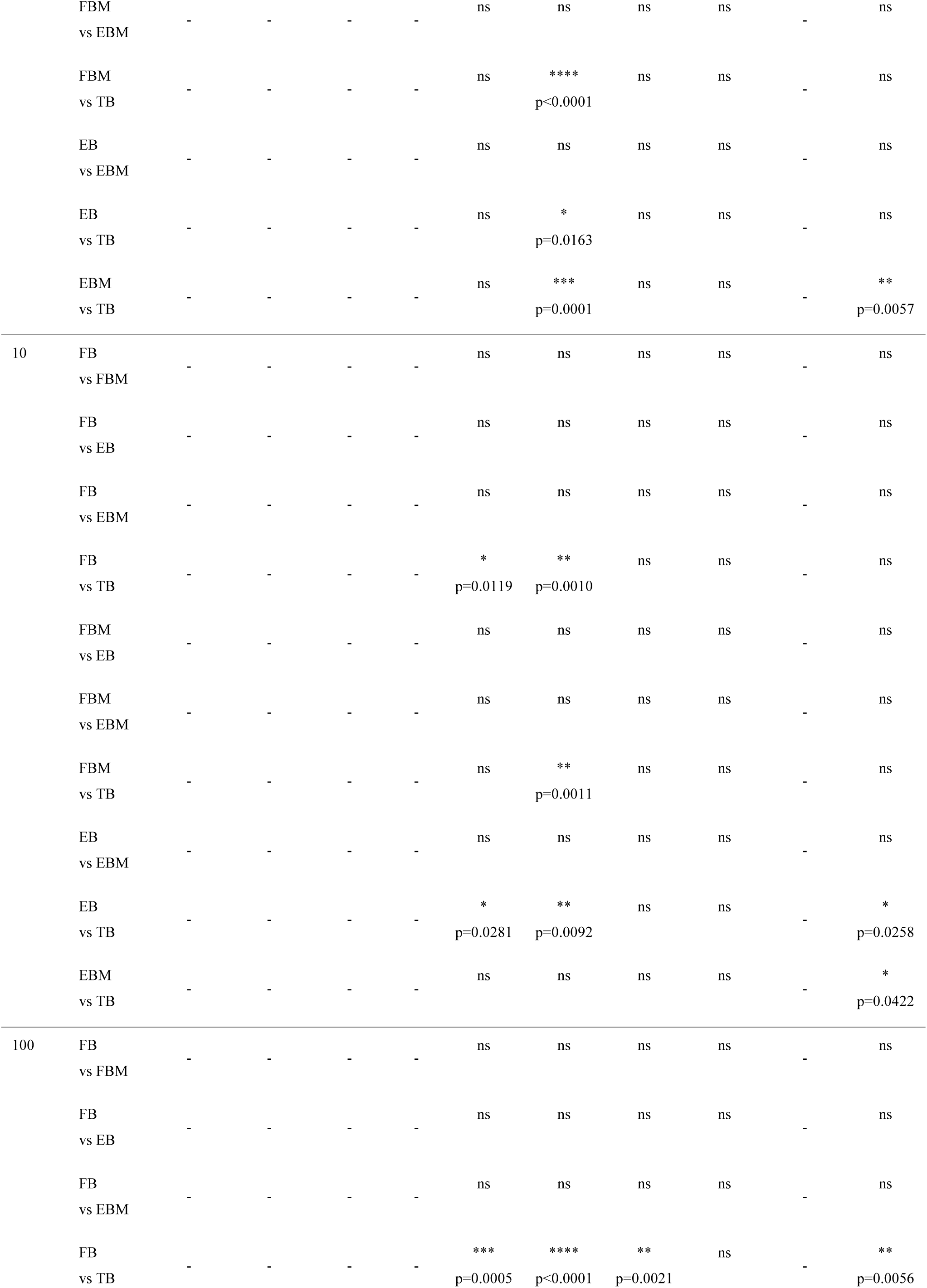
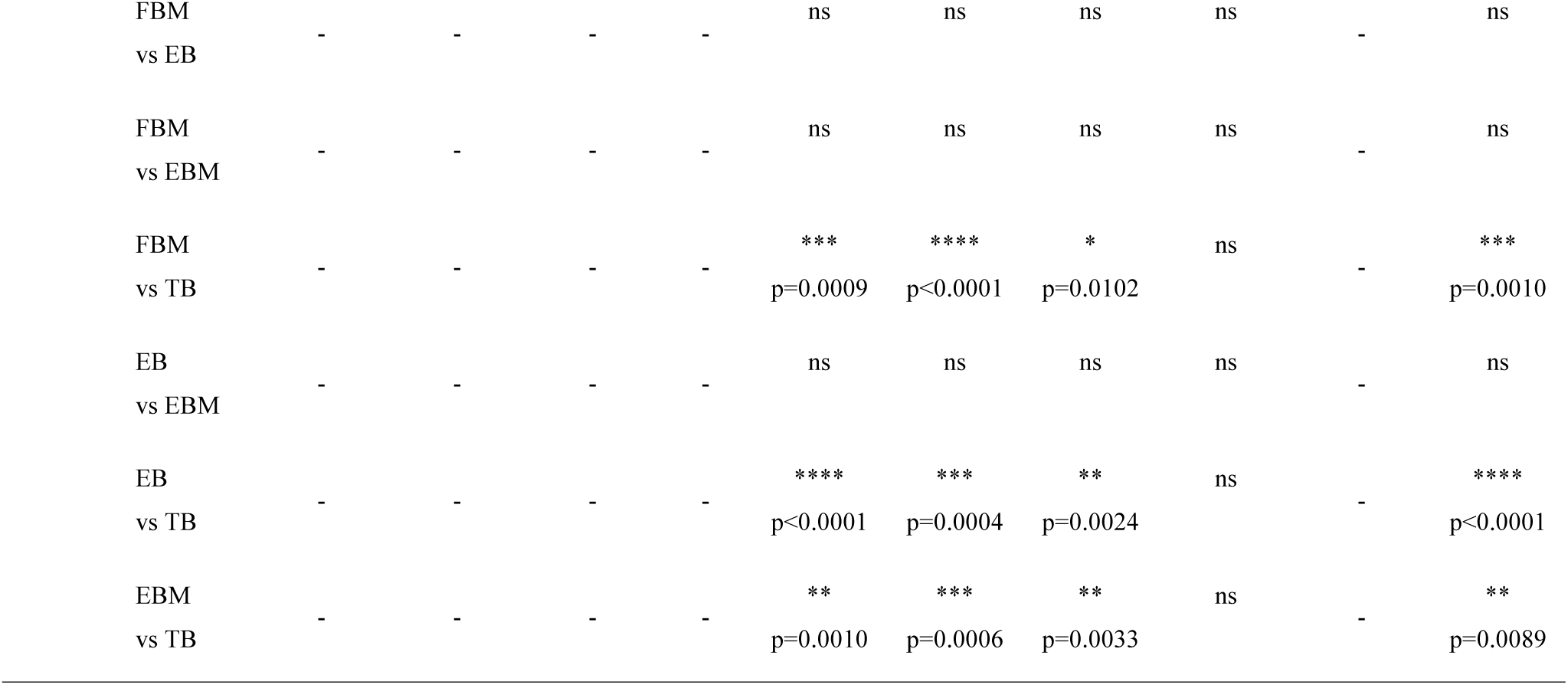
The significance of differences between the different categories of bundles in terms of apparent and net tensile mechanical properties assessed with a one-way ANOVA followed by a Tukey post hoc (ns p>0.05, *p≤0.05, **p≤0.01, ***p≤0.001, ****p≤0.0001).

**Table S6.**
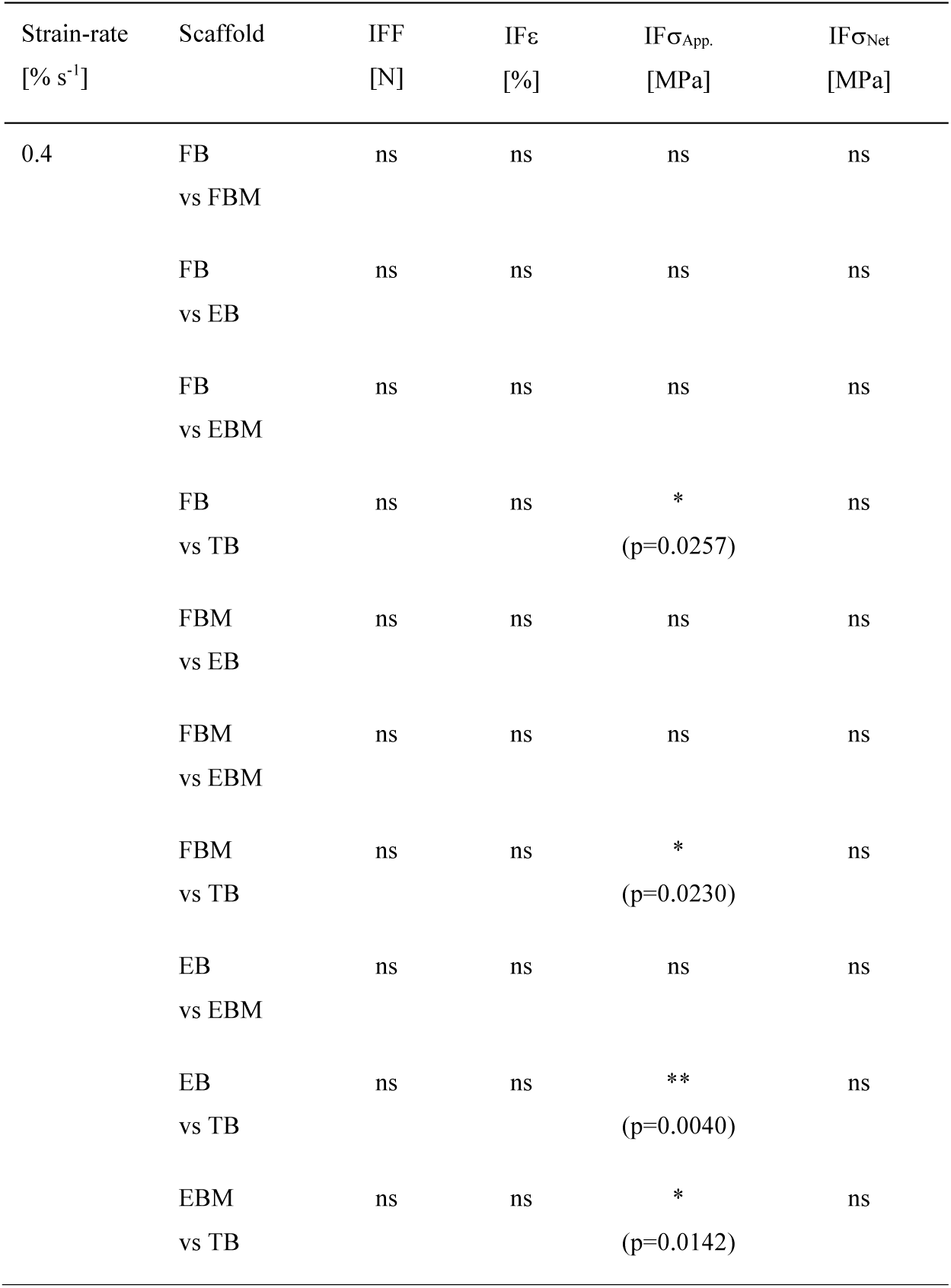
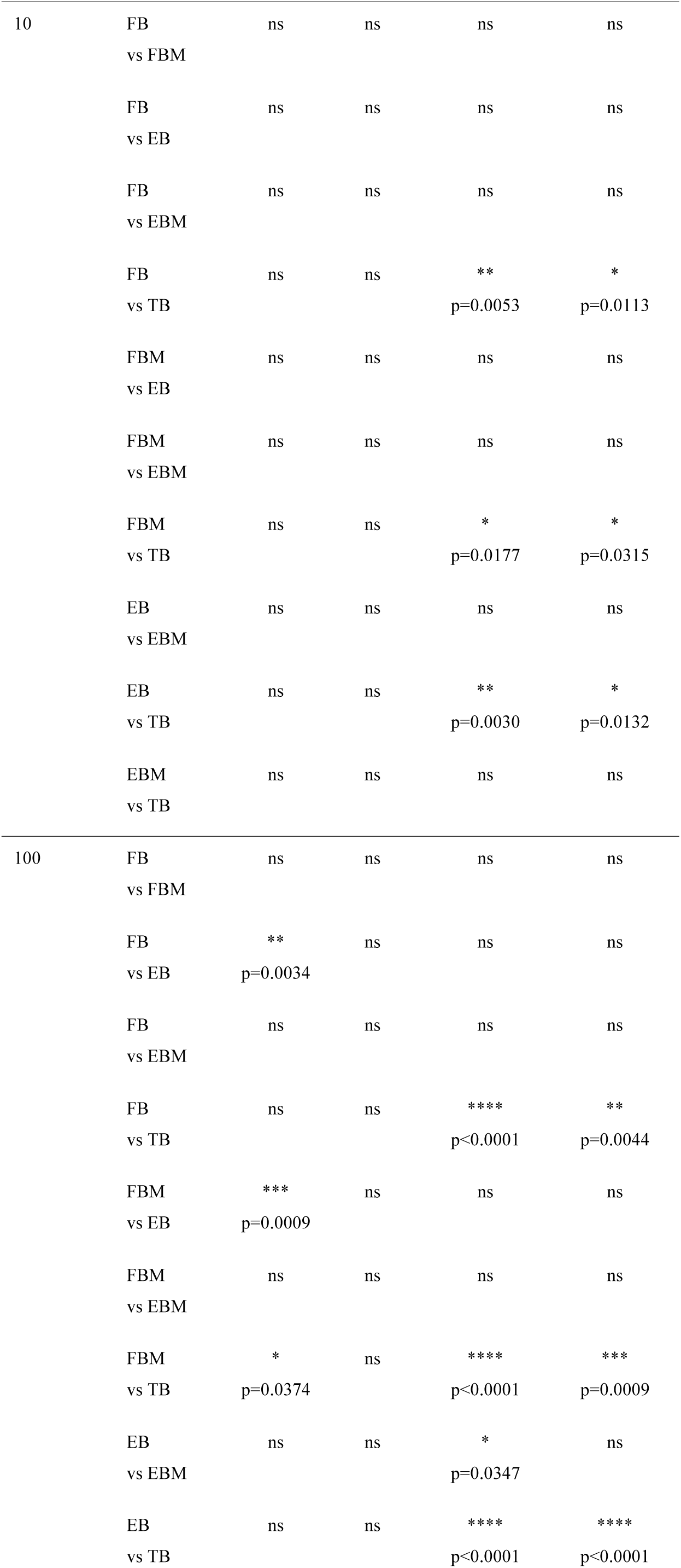
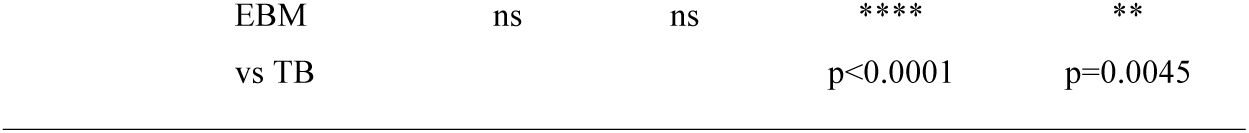
The significance of differences between the different categories of bundles in terms of apparent and net inflection point properties assessed with a one-way ANOVA followed by a Tukey post hoc (ns p>0.05, *p≤0.05, **p≤0.01, ***p≤0.001, ****p≤0.0001).

| Strain-rate<br>[% s<sup>-1</sup>] | Scaffold | IFF<br>[N] | IFε<br>[%] | IF$\sigma_{App.}$<br>[MPa] | IF$\sigma_{Net}$<br>[MPa] |
| --- | --- | --- | --- | --- | --- |
| 0.4 | FB<br>vs FBM | ns | ns | ns | ns |
|  | FB<br>vs EB | ns | ns | ns | ns |
|  | FB<br>vs EBM | ns | ns | ns | ns |
|  | FB<br>vs TB | ns | ns | *<br>(p=0.0257) | ns |
|  | FBM<br>vs EB | ns | ns | ns | ns |
|  | FBM<br>vs EBM | ns | ns | ns | ns |
|  | FBM<br>vs TB | ns | ns | *<br>(p=0.0230) | ns |
|  | EB<br>vs EBM | ns | ns | ns | ns |
|  | EB<br>vs TB | ns | ns | **<br>(p=0.0040) | ns |
|  | EBM<br>vs TB | ns | ns | *<br>(p=0.0142) | ns |
| 10 | FB<br>vs FBM | ns | ns | ns | ns |
|  | FB<br>vs EB | ns | ns | ns | ns |
|  | FB<br>vs EBM | ns | ns | ns | ns |
|  | FB<br>vs TB | ns | ns | **<br>p=0.0053 | *<br>p=0.0113 |
|  | FBM<br>vs EB | ns | ns | ns | ns |
|  | FBM<br>vs EBM | ns | ns | ns | ns |
|  | FBM<br>vs TB | ns | ns | *<br>p=0.0177 | *<br>p=0.0315 |
|  | EB<br>vs EBM | ns | ns | ns | ns |
|  | EB<br>vs TB | ns | ns | **<br>p=0.0030 | *<br>p=0.0132 |
|  | EBM<br>vs TB | ns | ns | ns | ns |
| 100 | FB<br>vs FBM | ns | ns | ns | ns |
|  | FB<br>vs EB | **<br>p=0.0034 | ns | ns | ns |
|  | FB<br>vs EBM | ns | ns | ns | ns |
|  | FB<br>vs TB | ns | ns | ****<br>p<0.0001 | **<br>p=0.0044 |
|  | FBM<br>vs EB | ***<br>p=0.0009 | ns | ns | ns |
|  | FBM<br>vs EBM | ns | ns | ns | ns |
|  | FBM<br>vs TB | *<br>p=0.0374 | ns | ****<br>p<0.0001 | ***<br>p=0.0009 |
|  | EB<br>vs EBM | ns | ns | *<br>p=0.0347 | ns |
|  | EB<br>vs TB | ns | ns | ****<br>p<0.0001 | ****<br>p<0.0001 |

|  |  |  |  |  |
| --- | --- | --- | --- | --- |
| EBM<br>vs TB | ns | ns | ****<br>p<0.0001 | **<br>p=0.0045 |

**Table S7.** The significance of differences of apparent and net mechanical properties of bundles at different strain-rates assessed with a one-way ANOVA followed by a Tukey post hoc (ns p>0.05, *p≤0.05, **p≤0.01, ***p≤0.001, ****p≤0.0001).

|  | Scaffold | F<sub>Y</sub><br>[N] | F<sub>F</sub><br>[N] | ε<sub>Y</sub><br>[%] | ε<sub>F</sub><br>[%] | σ<sub>Y</sub><br>[MPa] | σ<sub>F</sub><br>[MPa] | W<sub>Y</sub><br>[J mm<sup>-3</sup>] | W<sub>F</sub><br>[J mm<sup>-3</sup>] | K<br>[N mm<sup>-1</sup>] | E<br>[MPa] |
| --- | --- | --- | --- | --- | --- | --- | --- | --- | --- | --- | --- |
| **App** | FB | ns | ns | ns | ns | * | * | ns | ns | ns | * |
|  | 0.4 vs 10 |  |  |  |  | p=0.0363 | p=0.0256 |  |  |  | p=0.0278 |
|  | FB | * | ** | ns | ns | ** | ** | * | ns | ns | ns |
|  | 0.4 vs 100 | p=0.0135 | p=0.0067 |  |  | p=0.0096 | p=0.0040 | p=0.0262 |  |  |  |
|  | FB | ns | ns | ns | ns | ns | ns | ns | ns | ns | ns |
|  | 10 vs 100 |  |  |  |  |  |  |  |  |  |  |
|  | FBM | ns | ns | ns | ns | ns | ns | ns | ns | ns | ns |
|  | 0.4 vs 10 |  |  |  |  |  |  |  |  |  |  |
|  | FBM | ns | * | ns | ns | ns | ** | ns | ns | ns | ns |
|  | 0.4 vs 100 |  | p=0.0156 |  |  |  | p=0.0077 |  |  |  |  |
|  | FBM | ns | ns | ns | ns | ns | ns | ns | ns | ns | ns |
|  | 10 vs 100 |  |  |  |  |  |  |  |  |  |  |
|  | EB | ns | ns | ns | ns | ns | ns | ns | ns | ns | ns |
|  | 0.4 vs 10 |  |  |  |  |  |  |  |  |  |  |
|  | EB | ns | ns | ns | ns | ns | ns | ns | ns | ns | ns |
|  | 0.4 vs 100 |  |  |  |  |  |  |  |  |  |  |
|  | EB | ns | ns | ns | ns | ns | ns | ns | ns | ns | ns |
|  | 10 vs 100 |  |  |  |  |  |  |  |  |  |  |
|  | EBM | ns | ns | ns | ns | * | ns | ns | ns | * | ** |
|  | 0.4 vs 10 |  |  |  |  | p=0.0446 |  |  |  | p=0.0105 | p=0.0023 |
|  | EBM | * | * | ns | ns | * | ** | ns | ns | *** | **** |
|  | 0.4 vs 100 | p=0.0212 | p=0.0190 |  |  | p=0.0105 | p=0.0087 |  |  | p=0.0003 | p<0.0001 |
|  | EBM | ns | ns | ns | ns | ns | ns | ns | ns | ns | ns |
|  | 10 vs 100 |  |  |  |  |  |  |  |  |  |  |
|  | TB | * | ns | ns | ns | ns | ns | ns | ns | ** | ns |
|  | 0.4 vs 10 | p=0.0234 |  |  |  |  |  |  |  | p=0.0070 |  |
|  | TB | ** | * | ns | ns | ** | ** | ** | ns | * | * |
|  | 0.4 vs 100 | p=0.0041 | p=0.0105 |  |  | p=0.0011 | p=0.0022 | p=0.0038 |  | p=0.0217 | p=0.0129 |
|  | TB | ns | ns | ns | ns | * | ** | * | ns | ns | ns |
|  | 10 vs 100 |  |  |  |  | p=0.0209 | p=0.0035 | p=0.0382 |  |  |  |
| **Net** | FB | - | - | - | - | ns | ns | ns | ns | - | ns |
|  | 0.4 vs 10 |  |  |  |  |  |  |  |  |  |  |
|  | FB | - | - | - | - | * | ** | * | ns | - | ns |
|  | 0.4 vs 100 |  |  |  |  | p=0.0326 | p=0.0080 | p=0.0414 |  |  |  |
|  | FB | - | - | - | - | ns | ns | ns | ns | - | ns |
|  | 10 vs 100 |  |  |  |  |  |  |  |  |  |  |
|  | FBM | - | - | - | - | ns | ns | ns | ns | - | ns |
|  | 0.4 vs 10 |  |  |  |  |  |  |  |  |  |  |
|  | FBM | - | - | - | - | ns | ** | ns | ns | - | ns |
|  | 0.4 vs 100 |  |  |  |  |  | p=0.0043 |  |  |  |  |
|  | FBM | - | - | - | - | ns | ns | ns | ns | - | ns |
|  | 10 vs 100 |  |  |  |  |  |  |  |  |  |  |
|  | EB | - | - | - | - | ns | ns | ns | ns | - | ns |
|  | 0.4 vs 10 |  |  |  |  |  |  |  |  |  |  |
|  | EB | - | - | - | - | ns | ns | ns | ns | - | ns |
|  | 0.4 vs 100 |  |  |  |  |  |  |  |  |  |  |
|  | EB | - | - | - | - | ns | ns | ns | ns | - | ns |
|  | 10 vs 100 |  |  |  |  |  |  |  |  |  |  |
|  | EBM | - | - | - | - | * | ns | ns | ns | - | * |
|  | 0.4 vs 10 |  |  |  |  | p=0.0234 |  |  |  |  | p=0.0300 |
|  | EBM | - | - | - | - | ** | *** | ns | ns | - | ** |
|  | 0.4 vs 100 |  |  |  |  | p=0.0098 | p=0.0007 |  |  |  | p=0.0019 |
|  | EBM | - | - | - | - | ns | * | ns | ns | - | ns |
|  | 10 vs 100 |  |  |  |  |  | p=0.0190 |  |  |  |  |
|  | TB | - | - | - | - | ns | ns | ns | ns | - | ns |
|  | 0.4 vs 10 |  |  |  |  |  |  |  |  |  |  |
|  | TB | - | - | - | - | * | ns | ns | ns | - | ns |
|  | 0.4 vs 100 |  |  |  |  | p=0.0158 |  |  |  |  |  |
|  | TB | - | - | - | - | ns | ns | ns | ns | - | ns |
|  | 10 vs 100 |  |  |  |  |  |  |  |  |  |  |

**Table S8.**
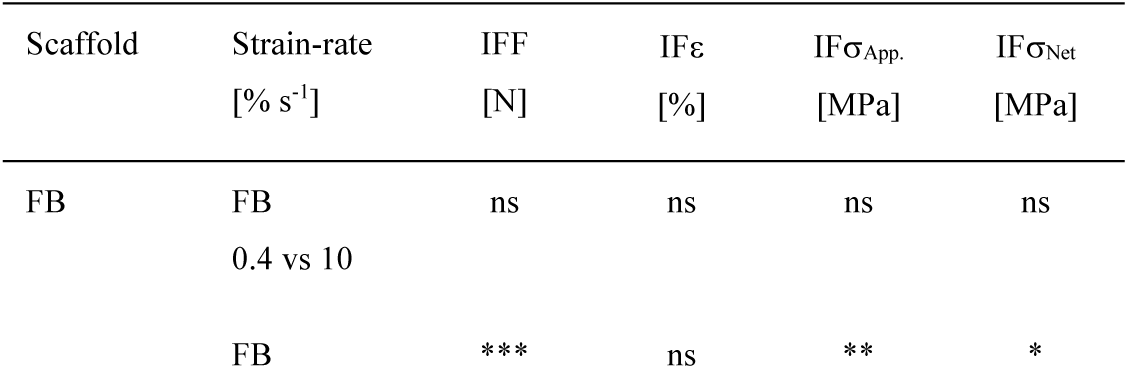
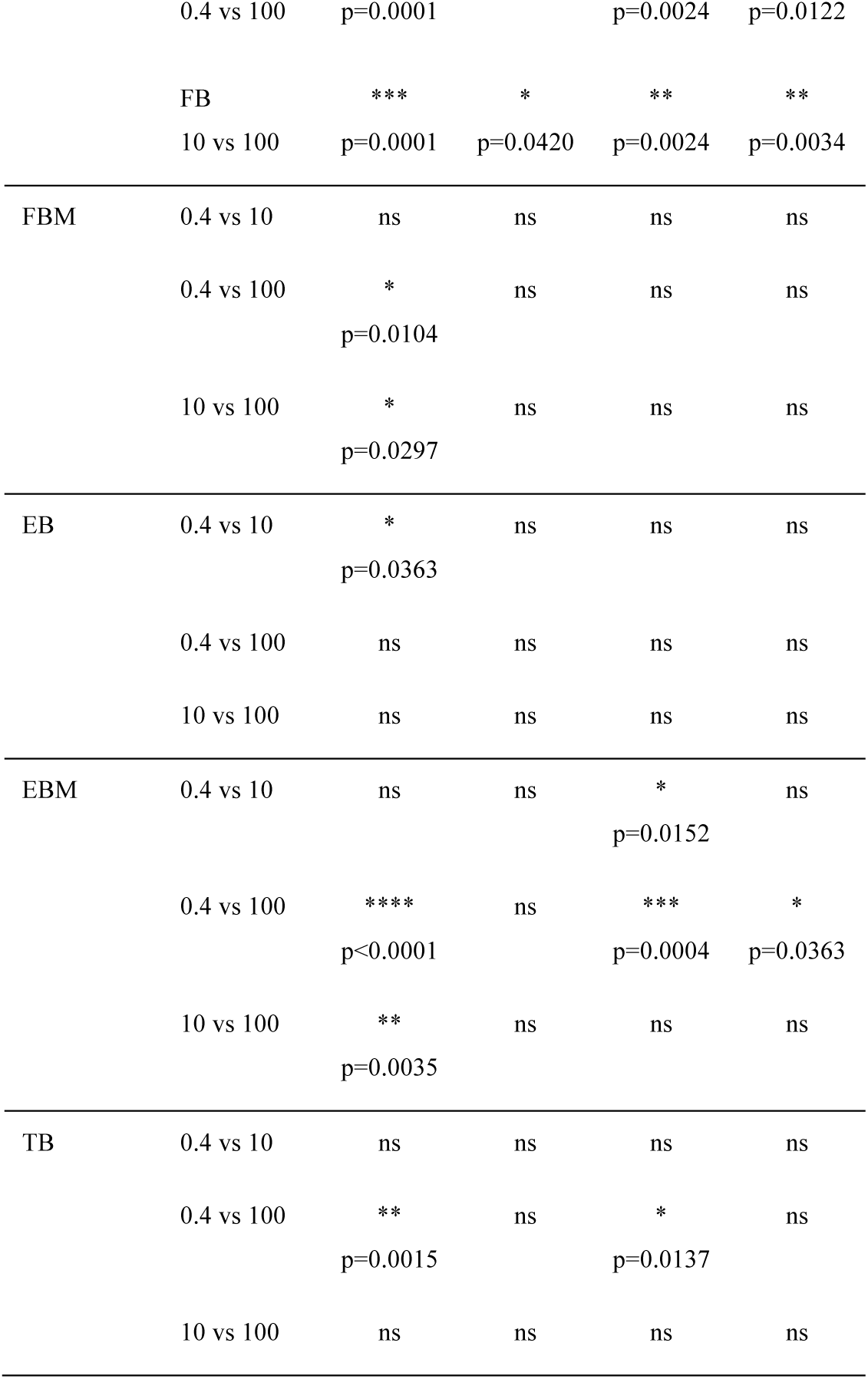
The significance of differences of the inflection points properties of bundles at different strain-rates assessed with a one-way ANOVA followed by a Tukey post hoc (ns p>0.05, *p≤0.05, **p≤0.01, ***p≤0.001, ****p≤0.0001).

| Scaffold | Strain-rate<br>[% s<sup>-1</sup>] | IFF<br>[N] | IFε<br>[%] | IF$\sigma_{App.}$<br>[MPa] | IF$\sigma_{Net}$<br>[MPa] |
| --- | --- | --- | --- | --- | --- |
| FB | FB | ns | ns | ns | ns |
|  | 0.4 vs 10 |  |  |  |  |
|  | FB | *** | ns | ** | * |
|  | 0.4 vs 100 | p=0.0001 |  | p=0.0024 | p=0.0122 |
|  | FB | *** | * | ** | ** |
|  | 10 vs 100 | p=0.0001 | p=0.0420 | p=0.0024 | p=0.0034 |
| FBM | 0.4 vs 10 | ns | ns | ns | ns |
|  | 0.4 vs 100 | * | ns | ns | ns |
|  |  | p=0.0104 |  |  |  |
|  | 10 vs 100 | * | ns | ns | ns |
|  |  | p=0.0297 |  |  |  |
| EB | 0.4 vs 10 | * | ns | ns | ns |
|  |  | p=0.0363 |  |  |  |
|  | 0.4 vs 100 | ns | ns | ns | ns |
|  | 10 vs 100 | ns | ns | ns | ns |
| EBM | 0.4 vs 10 | ns | ns | * | ns |
|  |  |  |  | p=0.0152 |  |
|  | 0.4 vs 100 | **** | ns | *** | * |
|  |  | p<0.0001 |  | p=0.0004 | p=0.0363 |
|  | 10 vs 100 | ** | ns | ns | ns |
|  |  | p=0.0035 |  |  |  |
| TB | 0.4 vs 10 | ns | ns | ns | ns |
|  | 0.4 vs 100 | ** | ns | * | ns |
|  |  | p=0.0015 |  | p=0.0137 |  |
|  | 10 vs 100 | ns | ns | ns | ns |

**Table S9.** Evolution of the orientation of actin filaments of hMSCs grown in the different regions of EB and EBM. In the table 0° represents the axial orientation while 90° the transversal one with respect to the axis of EB or EBM.

| Scaffolds, regions, time points | Frequency (%)<br>(range: 0°-18°) | Frequency (%)<br>(range: 78°-90°) |
| --- | --- | --- |
| **EB** |  |  |
| 7 days TB | 73 ± 3 | 1.2 ± 0.3 |
| 14 days TB | 67 ± 6 | 1.7 ± 0.5 |
| 28 days TB | 74 ± 2 | 1.0 ± 0.1 |
| 7 days J | 70 ± 13 | 1.6 ± 0.3 |
| 14 days J | 63 ± 7 | 2.2 ± 0.2 |
| 28 days J | 58 ± 6 | 2.6 ± 0.6 |
| 7 days FB | 38 ± 2 | 4.9 ± 0.3 |
| 14 days FB | 37 ± 4 | 6.5 ± 1.6 |
| 28 days FB | 49 ± 7 | 5.6 ± 3.5 |
| **EBM** |  |  |
| 7 days TB | 69 ± 10% | 1.9 ± 0.8 |
| 14 days TB | 68 ± 10% | 2.2 ± 0.9 |
| 28 days TB | 68 ± 9% | 2.1 ± 0.6 |
| 7 days J | 64 ± 2% | 2.0 ± 0.2 |
| 14 days J | 58 ± 10% | 2.5 ± 0.4 |
| 28 days J | 59 ± 8% | 3.2 ± 1.4 |
| 7 days FBM | 35 ± 5% | 7.9 ± 1.2 |
| 14 days FBM | 64 ± 10% | 2.1 ± 0.5 |
| 28 days FBM | 50 ± 11% | 3.8 ± 1.8 |

**Table S10.** Mean ± SD of the percentage of surface area of the ECM produced by hMSCs on EB and EBM in the different time points and regions (7, 14, 28 days; TB, J, FB) in terms of COL-I, COL-II, TNC and Hydroxyapatite on EBM.

| Scaffold | Region | Days of Culture | COL-I | COL-II | TNC | Hydroxyapatite |
| --- | --- | --- | --- | --- | --- | --- |
| **EB** | TB | 7 | 81±2 | 16±7 | 3.3±0.7 | - |
| | | 14 | 85±18 | 22±5 | 5.0±1.2 | - |
| | | 28 | 76±26 | 28±1 | 5.4±0.7 | - |
| | J | 7 | 65±23 | 20±7 | 1.9±1.1 | - |
| | | 14 | 77±11 | 24±3 | 2.9±0.7 | - |
| | | 28 | 61±17 | 29±2 | 4.4±0.2 | - |
| | FB | 7 | 89±6 | 24±11 | 2.1±0.5 | - |
| | | 14 | 91±16 | 21±2 | 2.3±0.4 | - |
| | | 28 | 101±23 | 44±2 | 2.4±0.7 | - |
| **EBM** | TB | 7 | 43±10 | 57.2±4.7 | 15±1 | 0.0±0.0 |
| | | 14 | 79±2 | 28±17 | 8.0±0.02 | 0.01±0.01 |
| | | 28 | 59±6 | 22±17 | 15±3 | 0.02±0.03 |
| | J | 7 | 71±34 | 28±2 | 14±0.4 | 1.48±1.23 |
| | | 14 | 49±29 | 21±1 | 9±2 | 0.18±0.03 |
| | | 28 | 65±3 | 31±6 | 11±2 | 1.50±1.00 |
| | FB | 7 | 33±17 | 30±6 | 11±2 | 3.95±1.72 |
| | | 14 | 50±24 | 20±3 | 8±1 | 1.34±0.33 |
| | | 28 | 12±11 | 9.8±4.9 | 13±2 | 1.01±0.77 |

**Table S11.**
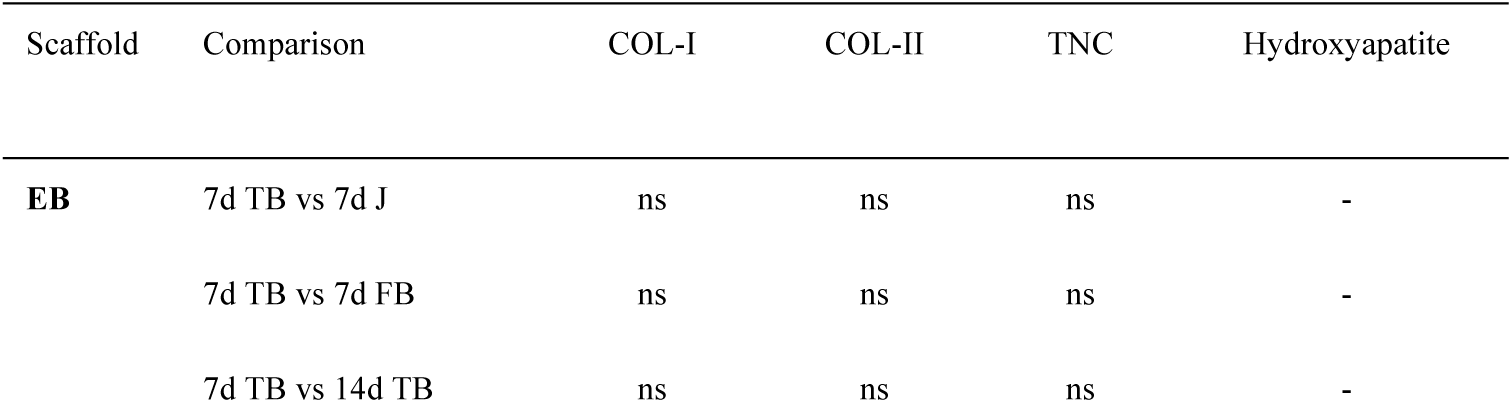
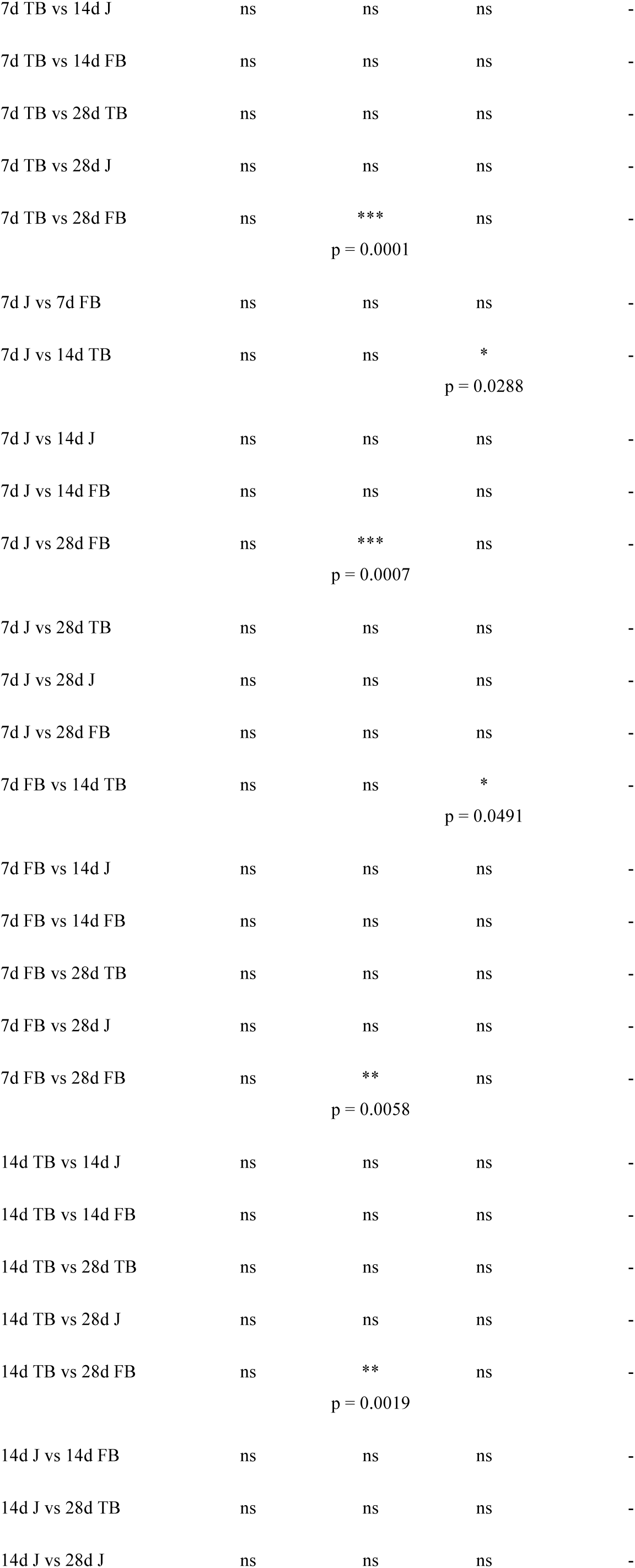
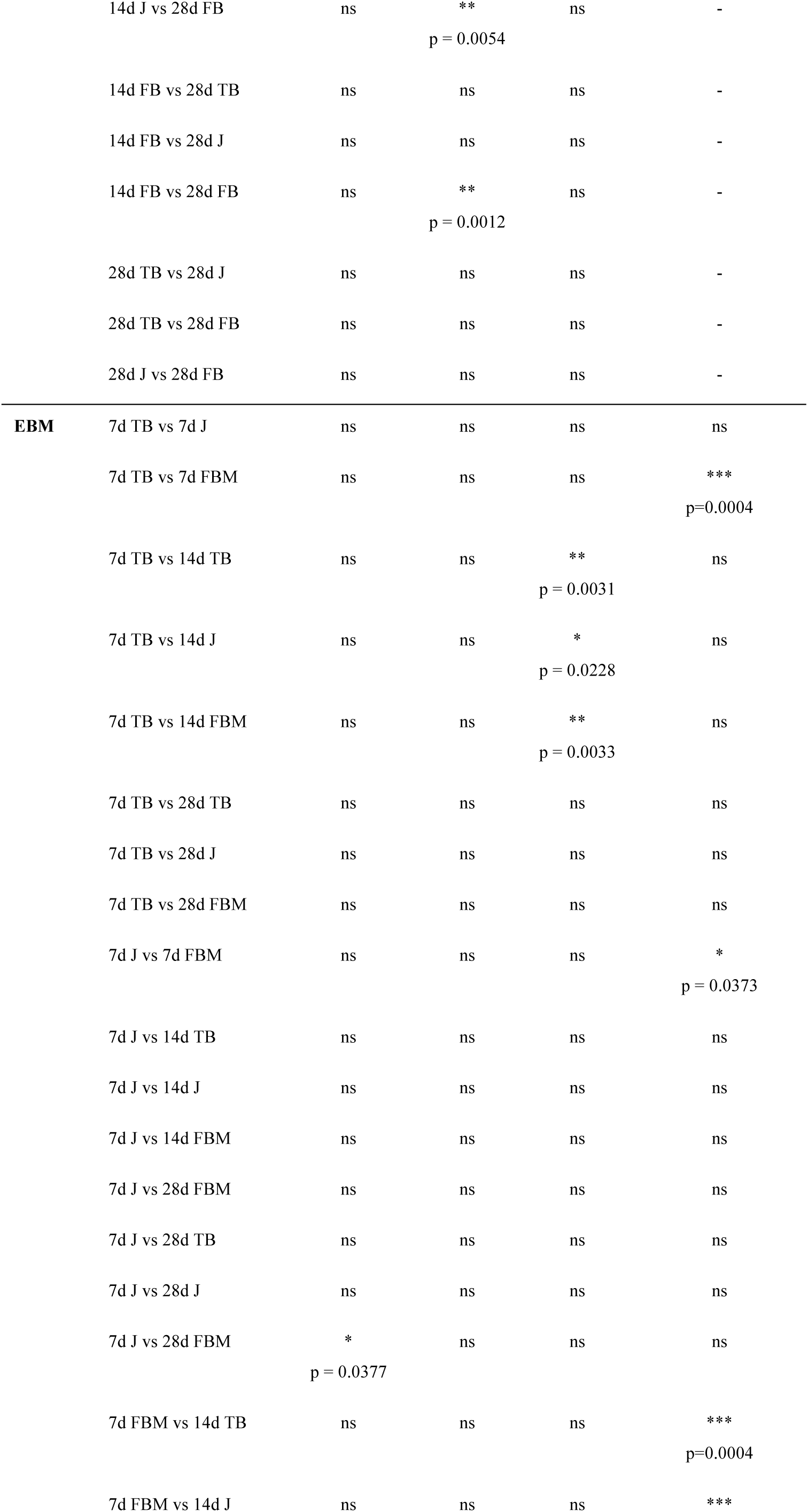
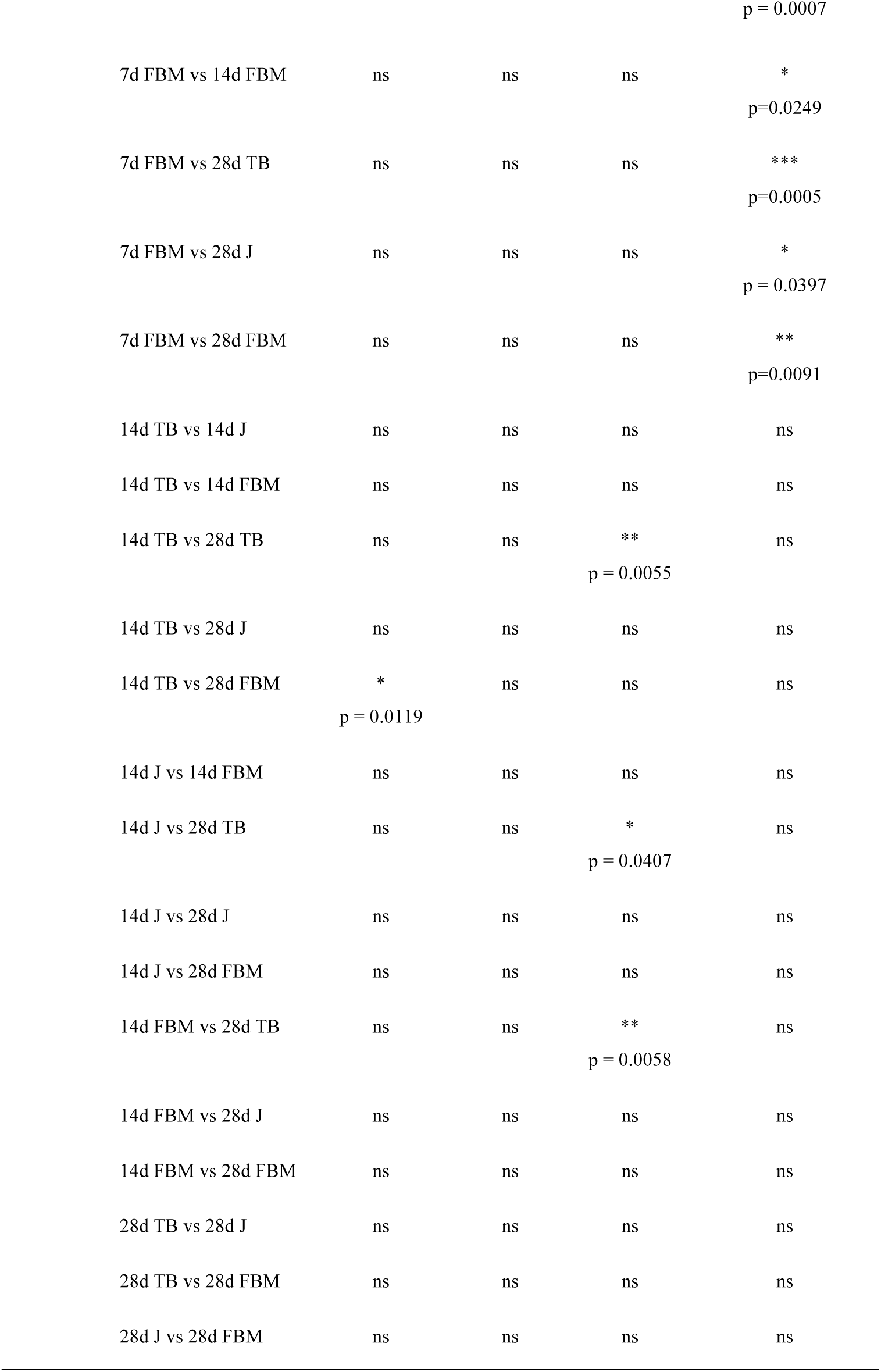
The significance of differences of the ECM quantifications and OsteoImage of EB and EBM between the different time points and regions (7, 14 and 28 days; TB, J, FB), assessed with a one-way ANOVA followed by a Tukey post hoc (ns p>0.05, *p≤0.05, **p≤0.01, ***p≤0.001, ****p≤0.0001).

**Table S12.** Gene expression of EB and EBM after 7, 14 and 28 days of hMSCs culture detected via RT-qPCR. Data expressed in fold change at day 1.

| Scaffold | Days of Culture | *Col-I*<br>[2<sup>-ΔΔCT</sup>] | *Scx*<br>[2<sup>-ΔΔCT</sup>] | *Tnc*<br>[2<sup>-ΔΔCT</sup>] | *Sox9*<br>[2<sup>-ΔΔCT</sup>] | *Col-II*<br>[2<sup>-ΔΔCT</sup>] | *Opn*<br>[2<sup>-ΔΔCT</sup>] | *Osx*<br>[2<sup>-ΔΔCT</sup>] | *Runx2*<br>[2<sup>-ΔΔCT</sup>] |
| --- | --- | --- | --- | --- | --- | --- | --- | --- | --- |
| **EB** | 7 | 67±22 | 4.7±0.5 | 6.3±1.0 | 1.4±0.3 | 3.2±1.2 | 2.4±1.2 | 4.4±1.5 | 3.4±0.8 |
|  | 14 | 38±2 | 5.1±0.8 | 2.8±0.1 | 1.9±0.1 | 2.8±0.1 | 2.0±0.8 | 7.2±1.8 | 2.7±0.6 |
|  | 28 | 58±2 | 6.8±1.9 | 2.8±1.3 | 1.7±0.2 | 3.1±0.4 | 1.5±0.4 | 13±1 | 2.4±0.5 |
| **EBM** | 7 | 0.9±0.1 | 4.7±1.7 | 0.6±0.3 | 39±13 | 22±1 | 5.9±0.4 | 7.9±2.4 | 8.9±0.1 |
|  | 14 | 1.1±0.3 | 6.2±1.3 | 0.4±0.1 | 45±14 | 50±9 | 4.4±0.7 | 9.0±4.1 | 8.3±0.4 |
|  | 28 | 0.6±0.1 | 13±1 | 0.2±0.04 | 74±11 | 70±8 | 2.3±0.2 | 9.2±2.2 | 8.9±0.5 |

**Table S13.** The significance of differences of each gene expressed by hMSCs on EB and EBM at the different time points of culture (7, 14 and 28 days), detected via RT-qPCR, assessed with a one-way ANOVA followed by a Tukey post hoc (ns p>0.05, *p≤0.05, **p≤0.01, ***p≤0.001, ****p≤0.0001).

| Scaffold | Days of Culture | *Col-I* | *Scx* | *Tnc* | *Sox9* | *Col-II* | *Opn* | *Osx* | *Runx2* |
| --- | --- | --- | --- | --- | --- | --- | --- | --- | --- |
| **EB** | 1 vs 7 | ***<br>p=0.0004 | *<br>p=0.0133 | ***<br>p=0.0002 | ns | *<br>p=0.0136 | ns | ns | **<br>p=0.0011 |
|  | 1 vs 14 | *<br>p=0.0152 | **<br>p=0.071 | ns | **<br>p=0.0029 | *<br>p=0.0368 | ns | **<br>p=0.0021 | *<br>p=0.0327 |
|  | 1 vs 28 | **<br>p=0.0011 | ***<br>p=0.0007 | ns | *<br>p=0.0108 | *<br>p=0.0169 | ns | ****<br>p<0.0001 | ns |
|  | 7 vs 14 | ns | ns | **<br>p=0.0036 | ns | ns | ns | ns | ns |
|  | 7 vs 28 | ns | ns | **<br>p=0.0031 | ns | ns | ns | ***<br>p=0.0002 | *<br>p=0.0436 |
|  | 14 vs 28 | ns | ns | ns | ns | ns | ns | **<br>p=0.0023 | ns |
| **EBM** | 1 vs 7 | ns | *<br>p=0.0269 | ns | *<br>p=0.0115 | *<br>p=0.0130 | ****<br>p<0.0001 | *<br>p=0.0491 | ****<br>p<0.0001 |
|  | 1 vs 14 | ns | **<br>p=0.0040 | **<br>p=0.0090 | **<br>p=0.050 | ****<br>p<0.0001 | ****<br>p<0.0001 | *<br>p=0.0241 | ****<br>p<0.0001 |
|  | 1 vs 28 | *<br>p=0.0196 | ****<br>p<0.0001 | **<br>p=0.0027 | ***<br>p=0.0002 | ****<br>p<0.0001 | *<br>p=0.0243 | *<br>p=0.0221 | ****<br>p<0.0001 |
|  | 7 vs 14 | ns | ns | ns | ns | **<br>p=0.0020 | ns | ns | ns |
|  | 7 vs 28 | ns | ***<br>p=0.0001 | ns | *<br>p=0.0210 | ****<br>p<0.0001 | ***<br>p=0.0003 | ns | ns |
|  | 14 vs 28 | **<br>p=0.0071 | ***<br>p=0.0006 | ns | ns | *<br>p=0.0168 | **<br>p=0.0014 | ns | ns |

**Table S14.**
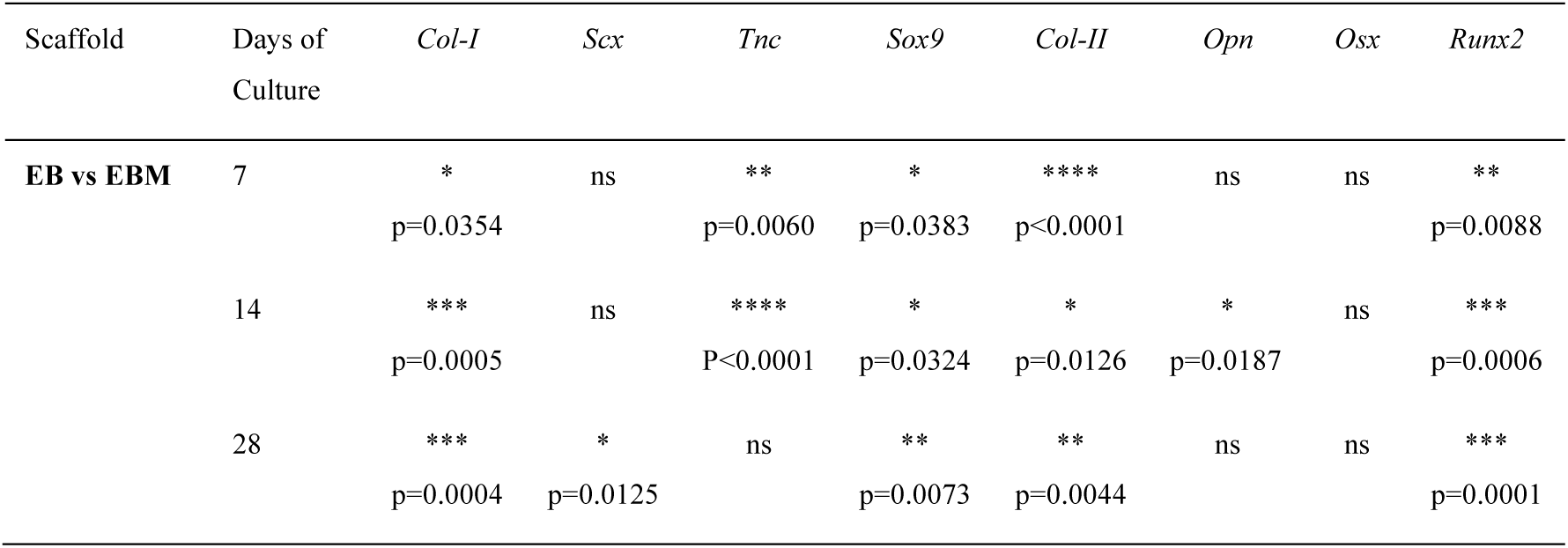
The significance of differences between EB and EBM of each gene expressed by hMSCs at the different time points of culture (7, 14 and 28 days), detected via RT-qPCR, assessed with an unpaired parametric t-test with Welch’s correction (ns p>0.05, *p≤0.05, **p≤0.01, ***p≤0.001, ****p≤0.0001).

**Table S15.** Elisa assay of EB and EBM after 7, 14 and 28 days of hMSCs culture.

| Scaffold | Days of Culture | COL-I<br>[ng $\mu\text{l}^{-1}$] | COL-II<br>[ng $\mu\text{l}^{-1}$] | TNC<br>[ng $\mu\text{l}^{-1}$] | TNMD<br>[ng $\mu\text{l}^{-1}$] |
| --- | --- | --- | --- | --- | --- |
| **EB** | 7 | 0.91±0.09 | 0.005±0.001 | 1.88±0.01 | 0.003±0.0009 |
|  | 14 | 1.17±0.25 | 0.005±0.001 | 1.82±0.02 | 0.004±0.0009 |
|  | 28 | 0.83±0.14 | 0.007±0.002 | 1.78±0.04 | 0.007±0.001 |
| **EBM** | 7 | 1.09±0.16 | 0.003±0.001 | 1.79±0.14 | 0.003±0.0004 |
|  | 14 | 1.29±0.16 | 0.007±0.0001 | 1.64±0.08 | 0.005±0.0005 |
|  | 28 | 0.24±0.01 | 0.009±0.001 | 1.79±0.05 | 0.005±0.003 |

**Table S16.** The significance of differences of each ECM protein expressed by hMSCs on EB and EBM at the different time points of culture (7, 14 and 28 days), detected via Elisa test, assessed with a one-way ANOVA followed by a Tukey post hoc (ns p>0.05, *p≤0.05, **p≤0.01, ***p≤0.001, ****p≤0.0001).

| Scaffold | Days of Culture | COL-I | COL-II | TNC | TNMD |
| --- | --- | --- | --- | --- | --- |
| **EB** | 7 vs 14 | ns | ns | ns | ns |
|  | 7 vs 28 | ns | ns | *<br>p=0.0113 | *<br>p=0.0184 |
|  | 14 vs 28 | ns | ns | ns | *<br>p=0.0267 |
| **EBM** | 7 vs 14 | ns | **<br>p=0.0084 | ns | ns |
|  | 7 vs 28 | ***<br>p=0.0005 | ***<br>p=0.0008 | ns | ns |
|  | 14 vs 28 | ***<br>p=0.0001 | ns | ns | ns |

**Table S17.** The significance of differences between EB and EBM of each ECM protein expressed by hMSCs at the different time points of culture (7, 14 and 28 days), detected via Elisa test, assessed with an unpaired parametric t-test with Welch’s correction (ns p>0.05, *p≤0.05, **p≤0.01, ***p≤0.001, ****p≤0.0001).

| Scaffold | Days of Culture | COL-I | COL-II | TNC | TNMD |
| --- | --- | --- | --- | --- | --- |
| **EB vs EBM** | 7 | ns | ns | ns | ns |
|  | 14 | ns | ns | ns | ns |
|  | 28 | * | ns | ns | ns |
| | | $p = 0.0176$ | | | |

**Table S18.** Calcium assay of EB and EBM after 7, 14 and 28 days of hMSCs culture compared with the control medium.

| Scaffold |  | Day 7 | Day 14 | Day 28 |
| --- | --- | --- | --- | --- |
| **Control medium**<br>(no cells) | Calcium<br>[nmol $\mu\text{l}^{-1}$] | 0.54±0.07 | 0.40±0.005 | 0.41±0.12 |
| **EB (cells)** | Calcium<br>[nmol $\mu\text{l}^{-1}$] | 0.44±0.25 | 0.20±0.01 | 0.25±0.05 |
| **EBM (cells)** | Calcium<br>[nmol $\mu\text{l}^{-1}$] | 0.96±0.15 | 0.42±0.07 | 0.79±0.14 |

**Table S19.**
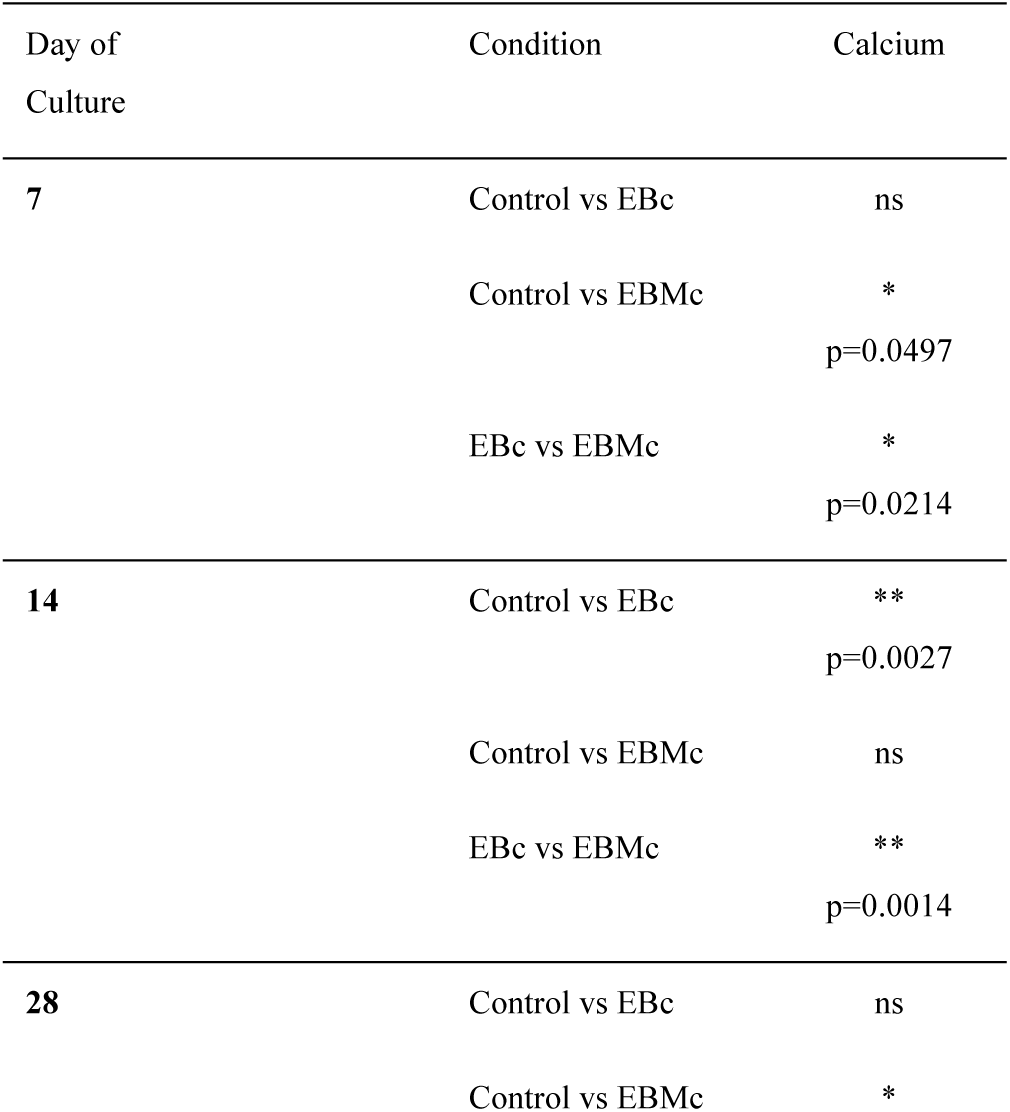
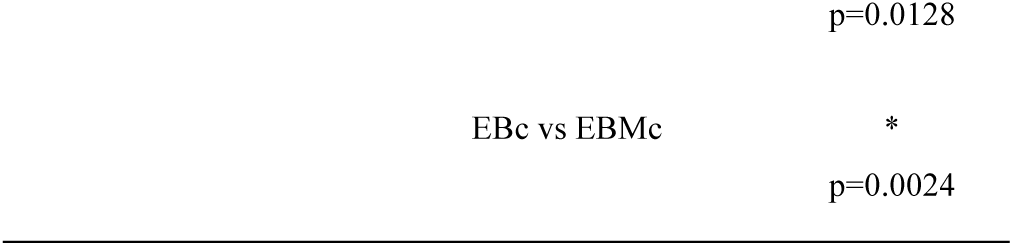
The significance of differences between the calcium assays of control medium, EB cellularized and cellularized with and without cells at different time points (7, 14 and 28 days) assessed with a one-way ANOVA followed by a Tukey post hoc (ns p>0.05, *p≤0.05, **p≤0.01, ***p≤0.001, ****p≤0.0001).

**Table S20.**
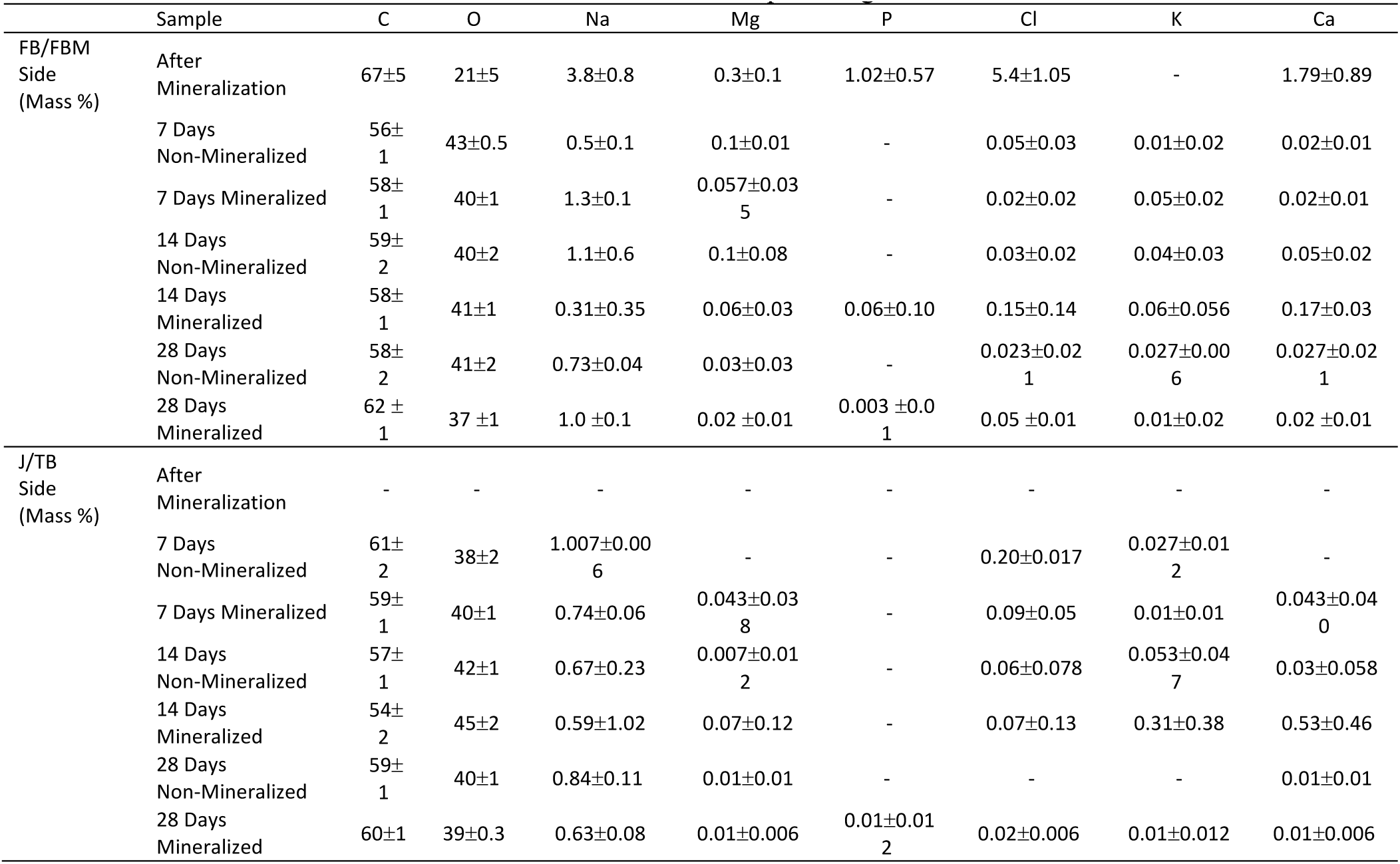
EDS investigations on the mineralized and non-mineralized bundles at the different time points of the hMSCs cultures. Amount of the different elements in terms of percentage of mass.

**Table S21.**
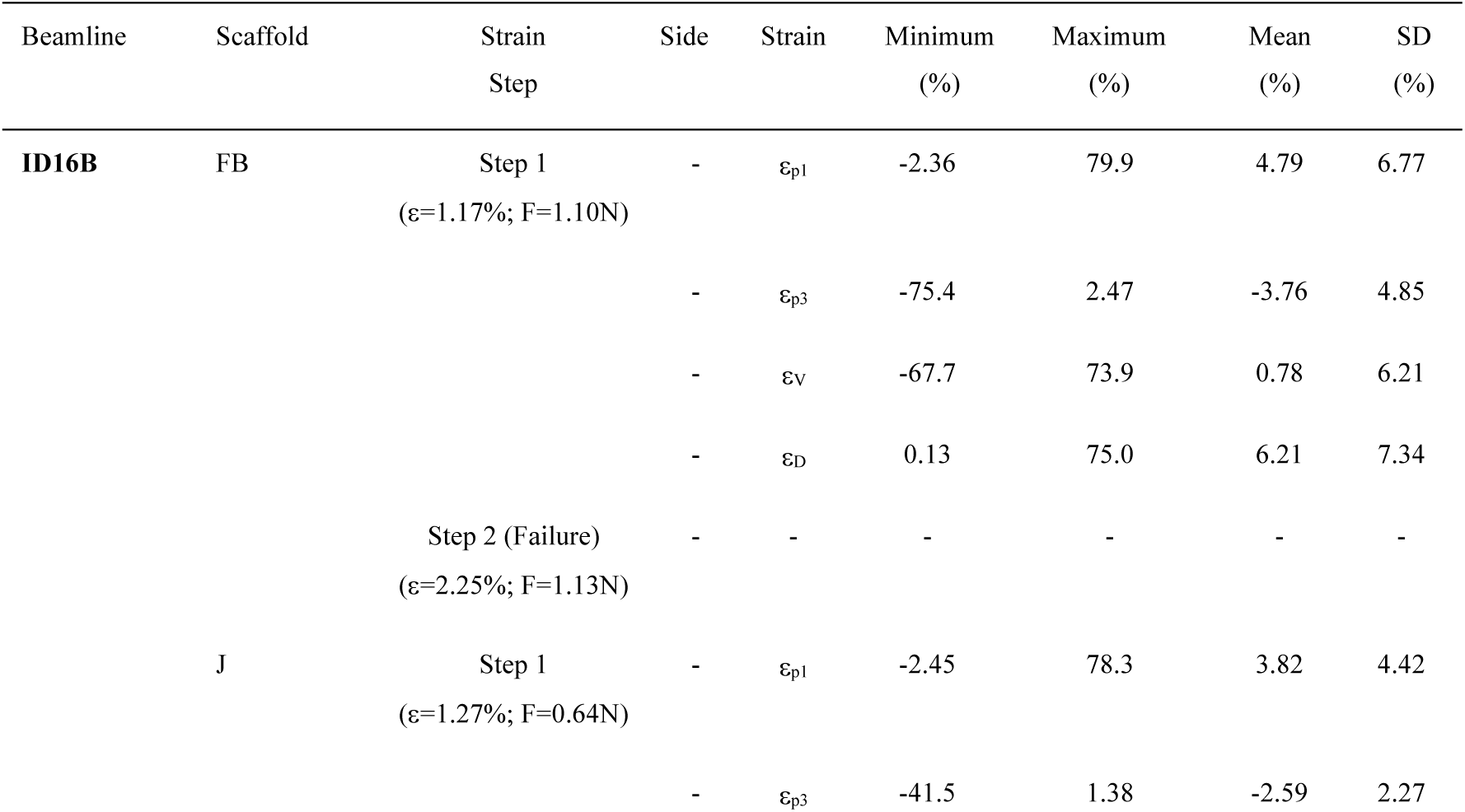
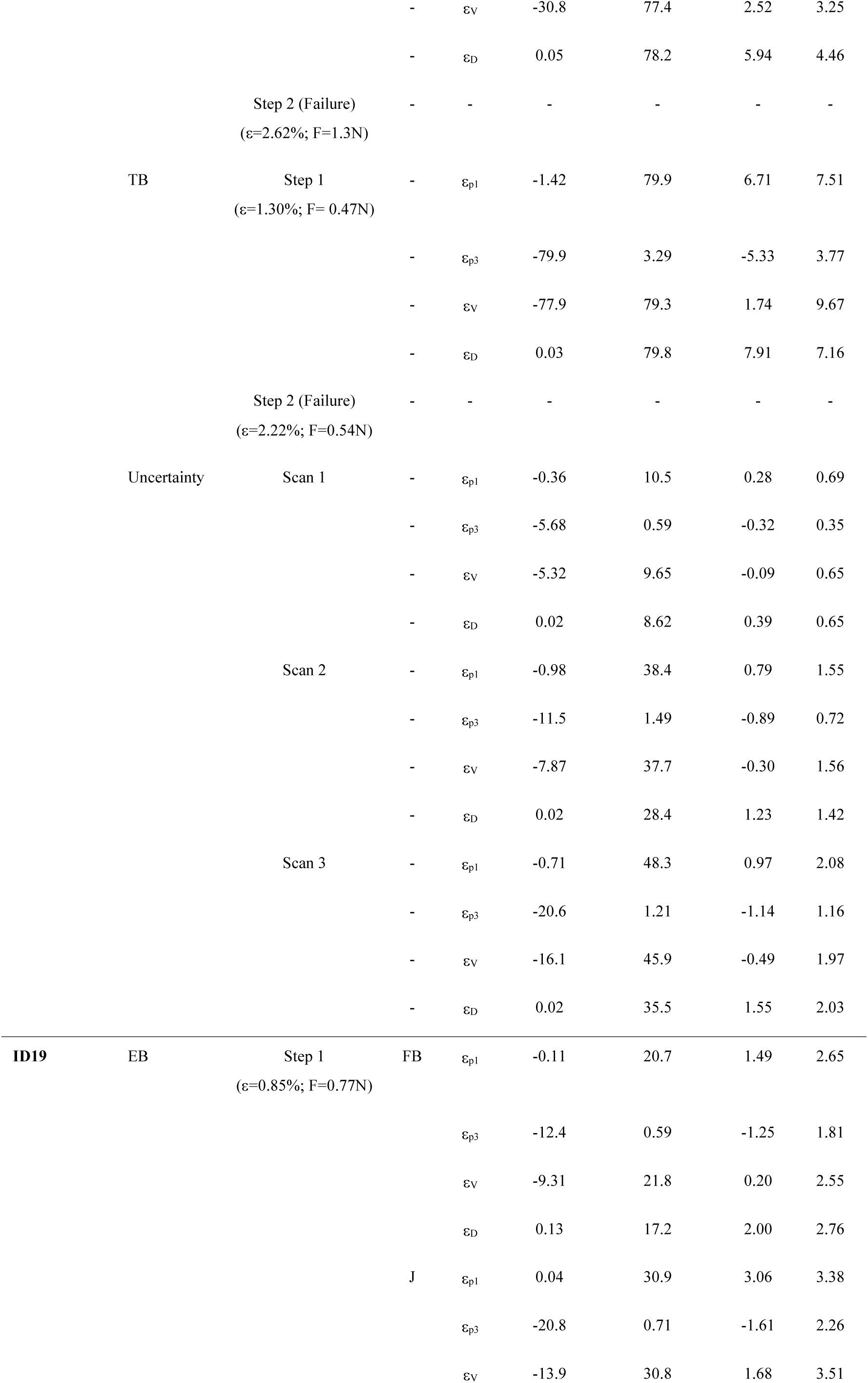
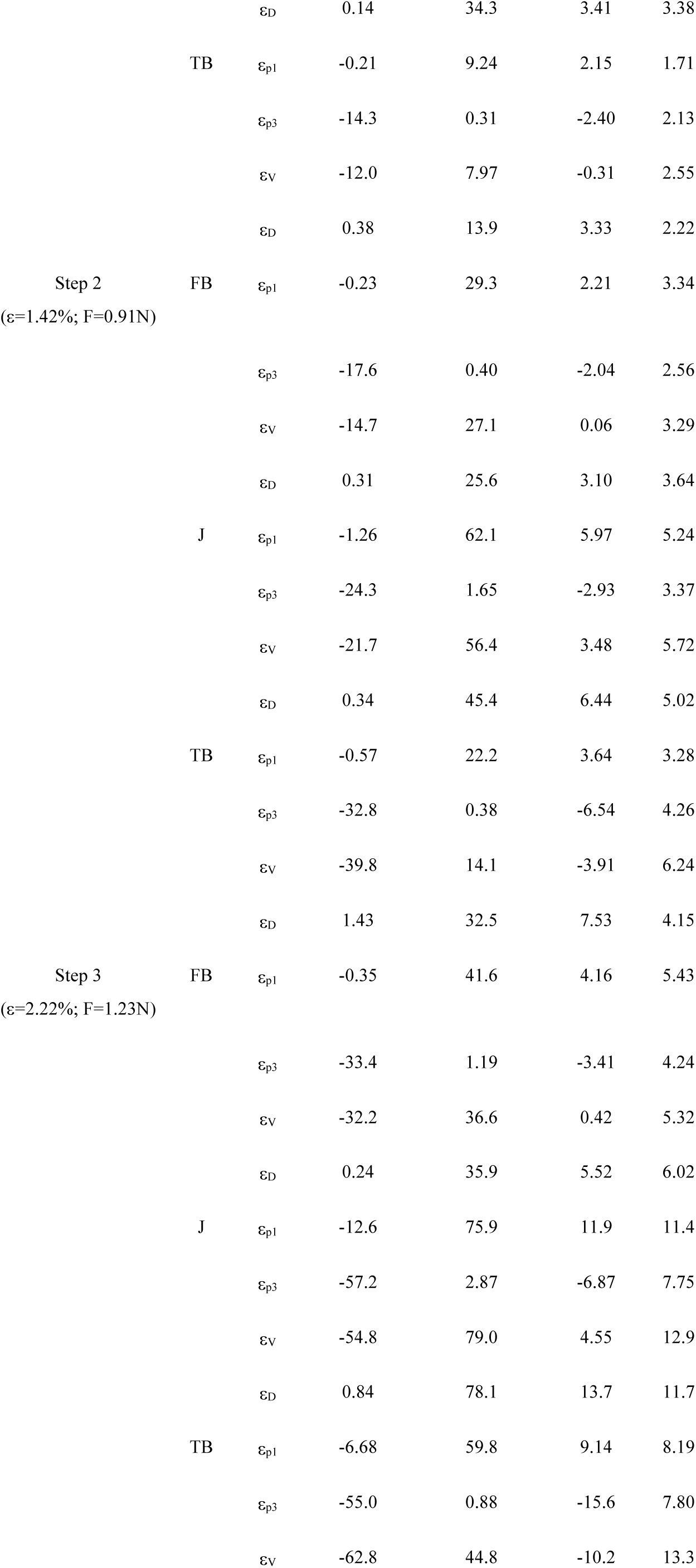
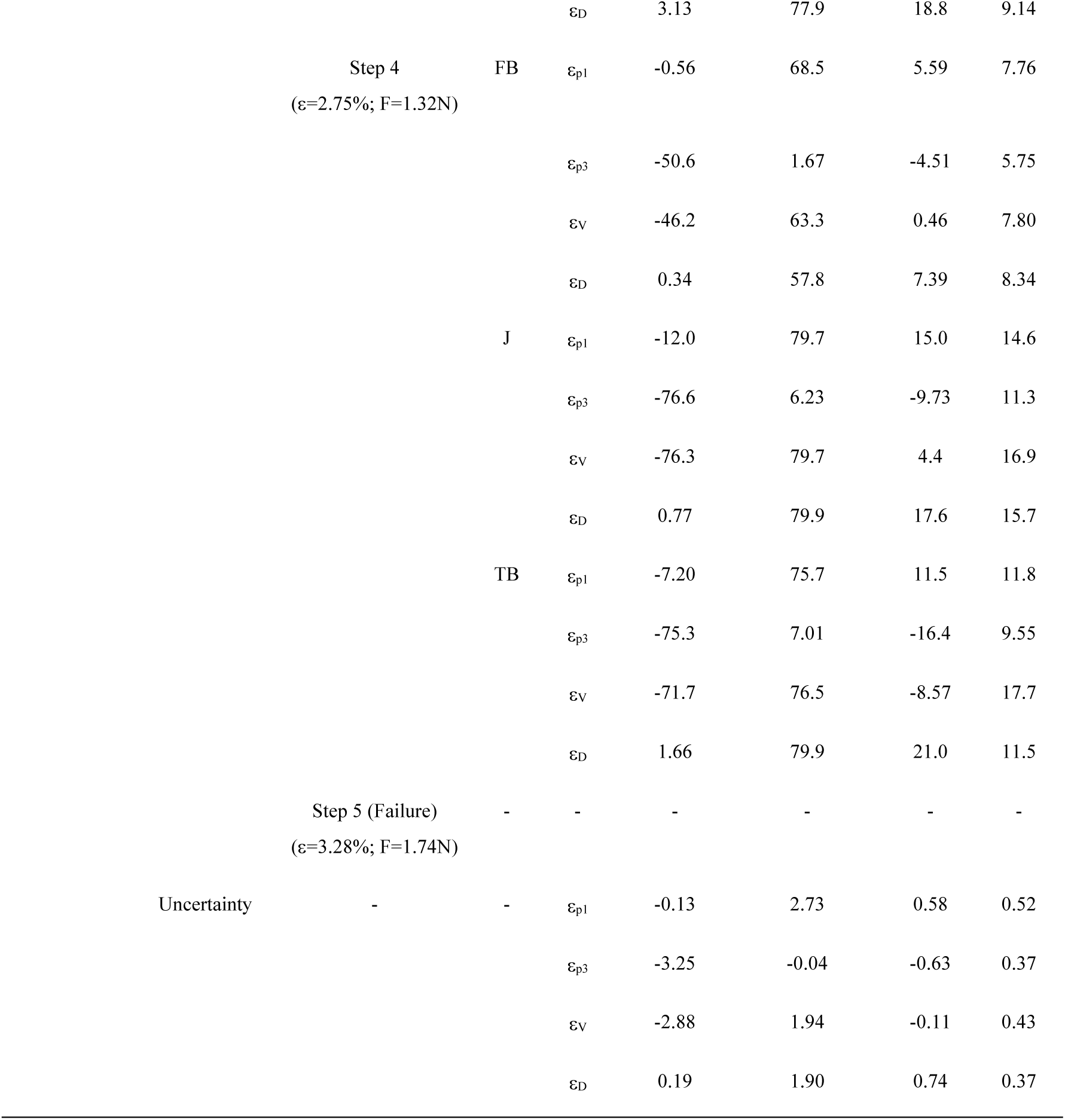
DVC mean, minimum, maximum and standard deviation values of χ_p1_, χ_p3_, χ_V,_ χ_D_ for FB, J, TB and EB sca<u>ffolds in the *in situ* tests at ID16B and ID19.</u>.

**Video S1.**
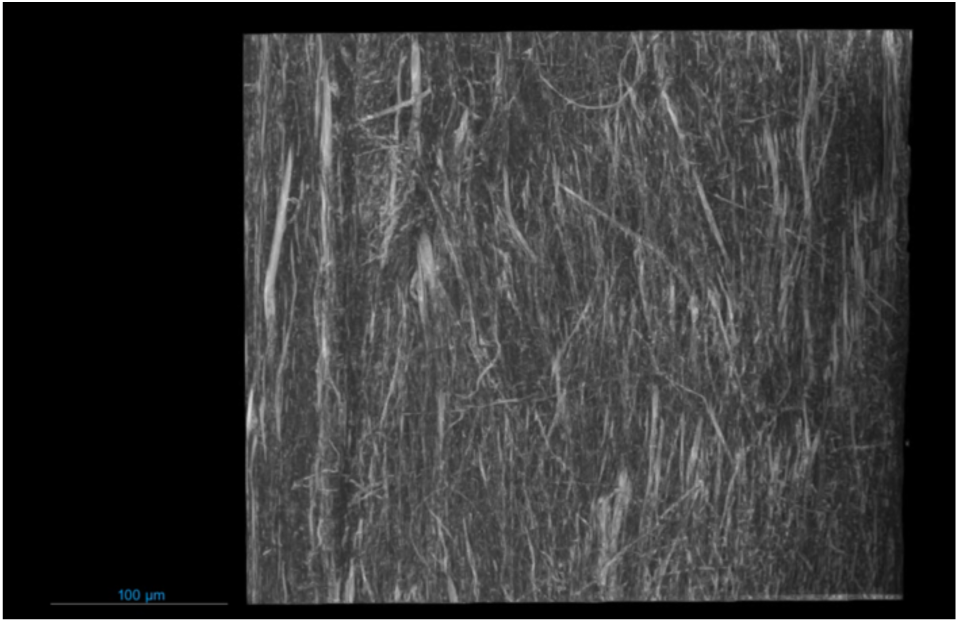
SnCT rendering of a FB at 300 nm of voxel size.

**Video S2.**
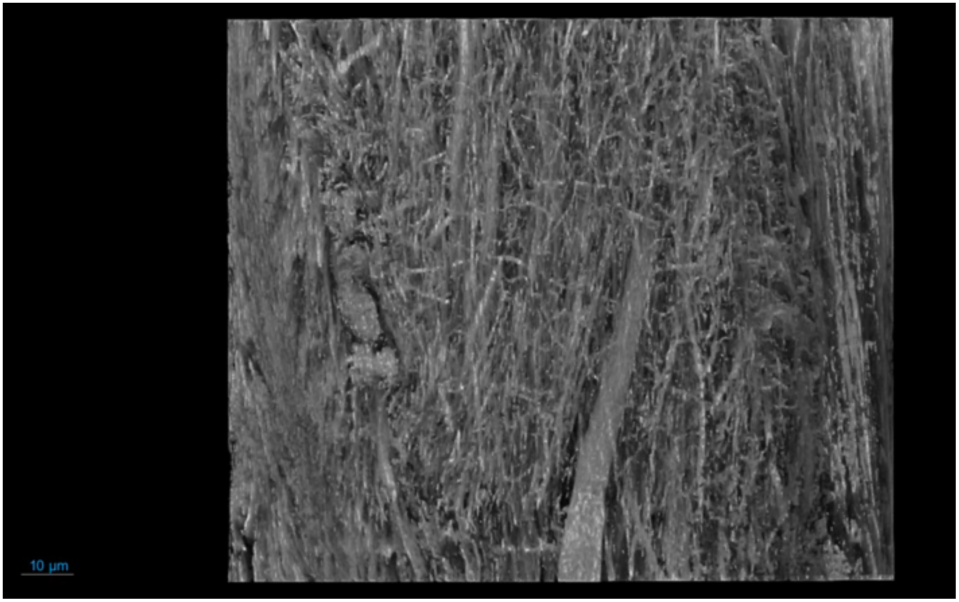
SnCT rendering of a FB at 100 nm of voxel size.

**Video S3.**
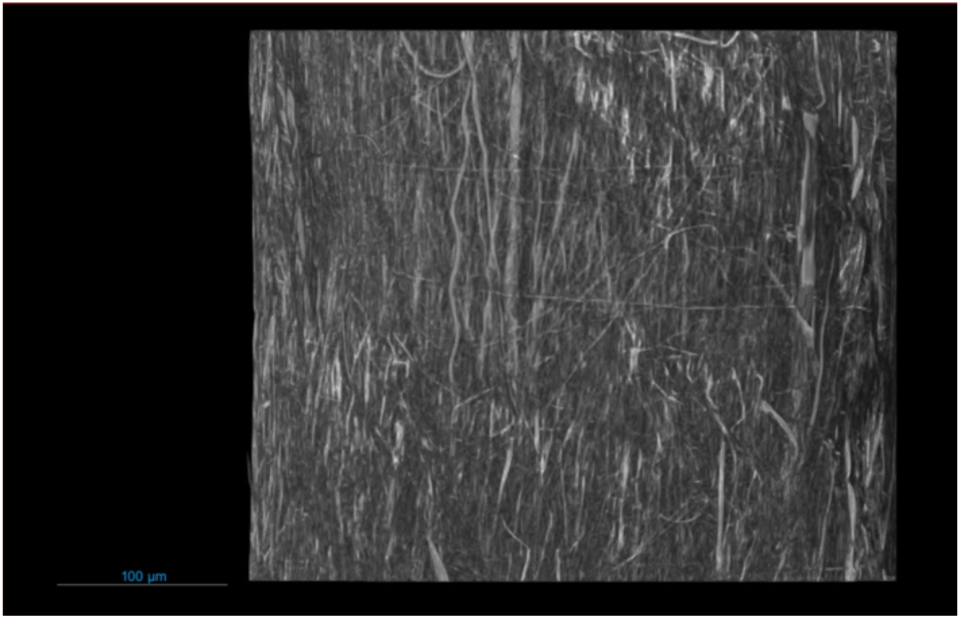
SnCT rendering of a TB at 300 nm of voxel size.

**Video S4.**
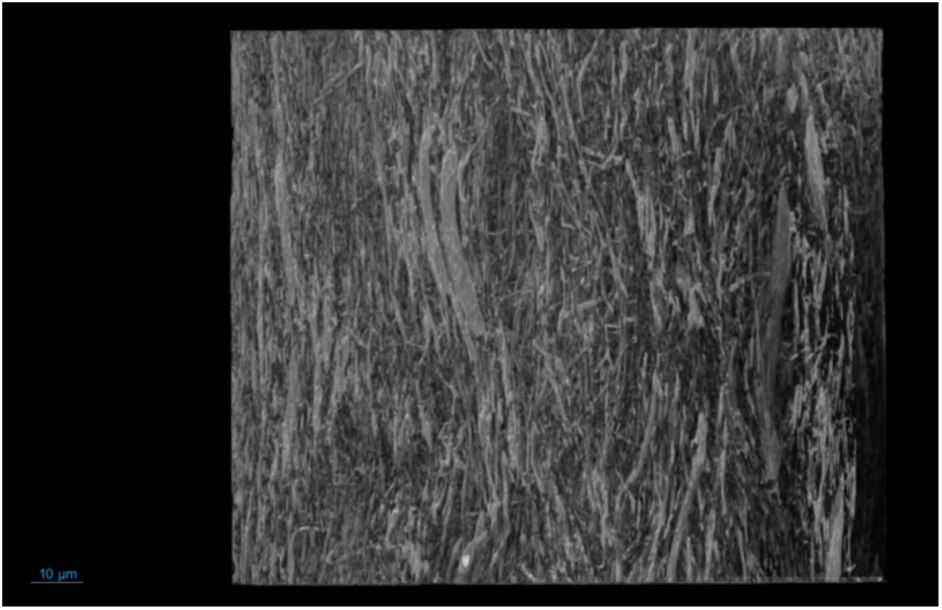
SnCT rendering of a TB at 100 nm of voxel size.

**Video S5.**
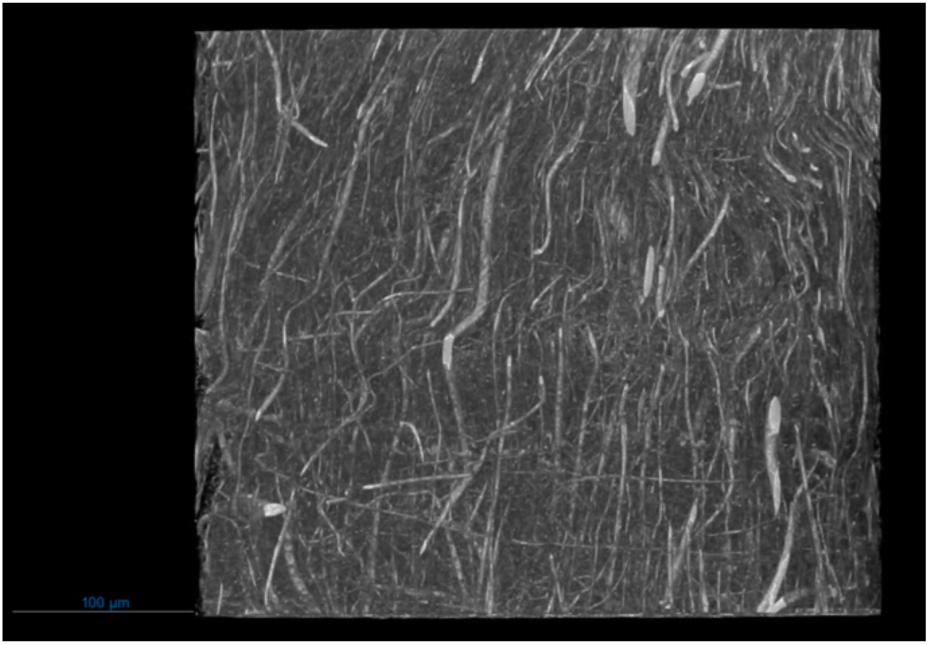
SnCT rendering of a J at 300 nm of voxel size.

**Video S6.**
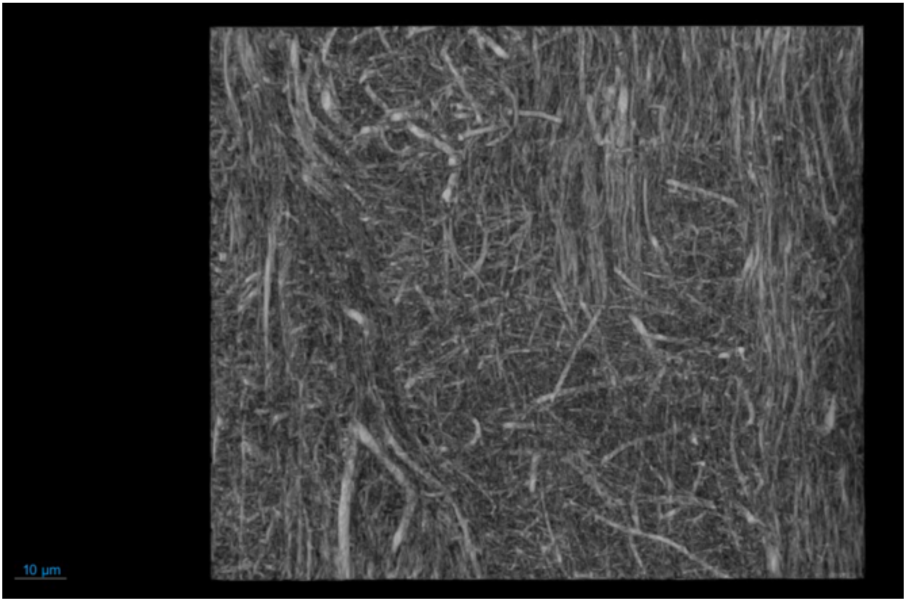
SnCT rendering of a J at 100 nm of voxel size.

**Video S7.**
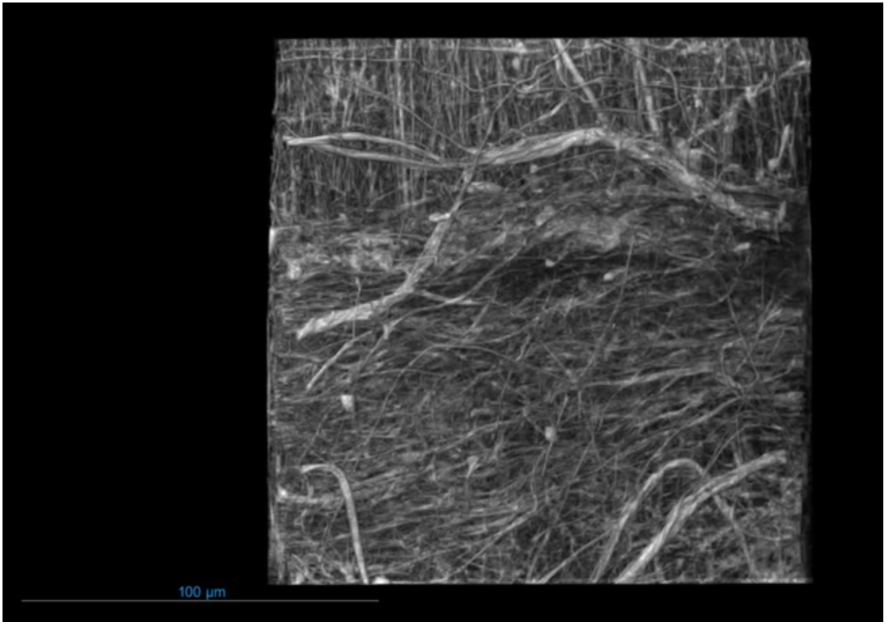
SnCT rendering of a J at 150 nm of voxel size.

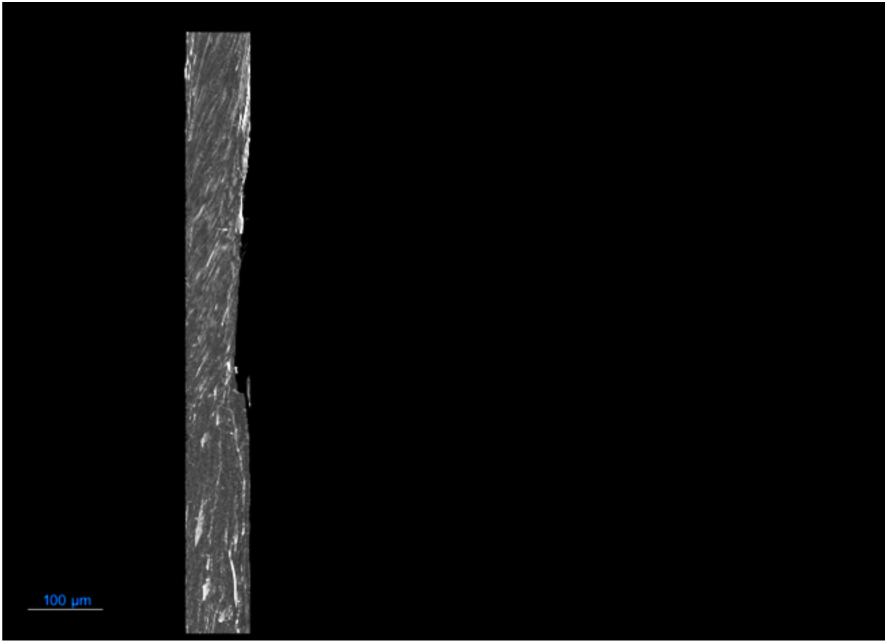

## Notes

### Competing Interest Statement

The authors have declared no competing interest.

